# FastaLake selects compact metaproteomic databases from large sequence reservoirs by de novo peptide lookup

**DOI:** 10.64898/2026.10.08.757395

**Authors:** Peter V. Treit, Florian A. Rosenberger, Philip Hartout, Mikhail Lebedev, Jinqiu Xiao, Maximilian Zwiebel, Philipp E. Geyer, Ankit Sinha, Georg Wallmann, Vincenth Brennsteiner, Kathrin Korff, Edwin H. Rodriguez, Wen-Feng Zeng, Maximilian T. Strauss, Marisa I. Keller, Michael Kuhn, Jonel Trebicka, Christophe Junot, Mani Arumugam, Jessica A. Grembi, Dari Shalon, Kerwyn Casey Huang, Karsten Borgwardt, Johannes B. Müller-Reif, Matthias Mann

**Affiliations:** Department of Proteomics and Signal Transduction, Max Planck Institute of Biochemistry, Martinsried, Germany; Department of Medical Biochemistry and Biophysics, Karolinska Institutet, Stockholm, Sweden; Department of Machine Learning and Systems Biology, Max Planck Institute of Biochemistry, Martinsried, Germany; Computational Systems Biochemistry Research Group, Max Planck Institute of Biochemistry, Martinsried, Germany; ions.bio GmbH, Martinsried, Germany; School of Engineering and School of Life Sciences, Westlake University, Hangzhou, China; Resolute Bio, Copenhagen, Denmark; Structural and Computational Biology Unit, European Molecular Biology Laboratory (EMBL), Heidelberg, Germany; Department of Internal Medicine B, University Hospital Münster, University of Münster, Münster, Germany; European Foundation for the Study of Chronic Liver Failure (EF-Clif), Barcelona, Spain; Université Paris-Saclay, CEA, Institut Joliot, Gif-sur-Yvette, France; Novo Nordisk Foundation Center for Basic Metabolic Research, Faculty of Health and Medical Sciences, University of Copenhagen, Copenhagen, Denmark; Department of Veterinary and Biomedical Sciences, Pennsylvania State University, University Park, PA, USA; Envivo Bio, Inc., San Francisco, CA, USA; Departments of Bioengineering and Microbiology and Immunology, Stanford University, Stanford, CA, USA

## Abstract

Metaproteomics analyses require protein sequence databases, yet the microbial proteins present in complex communities are not known in advance. Searching reference catalogs containing hundreds of millions of candidate sequences reduces identification sensitivity, while generating sample-specific databases through metagenomic sequencing adds time and cost. We introduce FastaLake, a computational workflow that uses full-length de novo peptide predictions by AlphaNovo to retrieve complete proteins from broad sequence resources and selects compact databases for conventional searching. In a defined eight-member community, FastaLake approached reference-search peptide yields without knowing its members. Using public sequence catalogs, FastaLake identified about as many peptides as matched metagenomic databases in two stool cohorts and 20–35% more in intestinal fluid at the same nominal 1% peptide false discovery rate. The resulting proteomes reveal personal functional profiles that persist across intestinal regions despite differences in leading species between intestinal fluid and stool. DNA, RNA and protein profiles agreed on broad differences between patients but only weakly on individual functions. All 726 acquisition-specific databases were smaller than the human reference proteome. Compact databases allowed non-specific searches of all 347 CapScan acquisitions, revealing histidine enrichment at peptide C termini in intestinal fluid. FastaLake enables metaproteomics analysis without matched metagenomic sequencing by constructing search databases from sequence resources and de novo peptide sequencing.

## Introduction

Mass spectrometry (MS)-based proteomics identifies peptides primarily by matching tandem mass spectra (MS/MS) to candidate sequences in a protein database [1,2]. The protein database therefore defines the peptide sequences that can be identified from the measurement. In human proteomics, this search space is well defined by a reference proteome of approximately 20,000 protein sequences that can be applied consistently across samples and individuals. In metaproteomics, however, the organisms and strains present in a microbial community cannot generally be specified a priori, and therefore neither can the relevant sample-specific protein sequence space. Large-scale metagenomic sequencing has generated comprehensive gut and global microbial sequence catalogs that provide broad sequence coverage but now contain hundreds of millions of proteins [3–5]. Searching these catalogs directly is computationally demanding and introduces large numbers of competing target and decoy sequences, reducing identification sensitivity [1,6]. Database construction is therefore a central challenge in metaproteomics: it determines both which microbial sequences can be identified and which alternatives compete to explain each spectrum. This challenge is particularly pronounced in complex communities where closely related organisms contribute large numbers of homologous protein sequences [1,2].

Current approaches aim to make this sequence space tractable by reducing it before the final database search. Matched metagenomic sequencing provides the most direct sample-specific resource but is also the most laborious and expensive, requiring DNA collection, sequencing, assembly, and gene prediction, with sequences potentially lost throughout the process. Reference database selection and protein sequence clustering reduce large catalogs computationally but can remove strain-specific sequence variation and thereby exclude peptides unique to individual strains. Iterative, two-stage approaches use an initial database search to select proteins or genomes for a second search [7,8]. When whole genomes are selected, all predicted proteins encoded by those genomes enter the second search, leaving the refined database comparatively large [9]. Combining these databases across samples can therefore produce very large study-level search spaces. Moreover, using the same spectra both to construct the reduced database and subsequently to identify peptides can bias conventional target–decoy estimation. The first search enriches the reduced database not only for genuine proteins but also for target sequences that happen to match those spectra well by chance. If the second-pass decoys have not undergone an equivalent spectrum-dependent selection process, false target matches and decoy matches are no longer statistically exchangeable, potentially leading to underestimation of the true false discovery rate (FDR) [10–12].

De novo peptide sequencing derives peptide sequence information directly from fragmentation spectra and therefore provides an approach to database construction that is independent of database searching [13]. MetaNovo maps short sequence tags to a broad protein resource and ranks the resulting proteins to construct a reduced database for conventional searching [14], while the uMetaP workflow uses filtered novoMP de novo peptide predictions for homology-based protein retrieval [15]. AlphaNovo predicts peptide sequences and is trained on MetaPTCM, a large corpus that includes non-tryptic peptides and diverse post-translational and chemical modifications alongside conventional tryptic spectra [16]. This coverage is particularly relevant to gut metaproteomics, where endogenous proteolysis may generate peptide termini beyond those expected in a conventional tryptic search.

We introduce a de novo-driven pipeline for metaproteomics raw data analysis called FastaLake. The name refers to the data lake of data engineering, in which raw data are stored unprocessed and structured only when queried. FastaLake likewise keeps context-specific protein sequences in a reservoir without sequence-similarity clustering and selects from it only when a sample is analyzed. Unlike tag ranking in MetaNovo, it uses complete AlphaNovo predictions as exact sequence queries to select compact protein databases from this sequence reservoir. We search the selected databases with Sage [17] and validate the results with a second search engine. We first establish the workflow, performance and robustness of FastaLake across technical and biological variation in stool and in a defined eight-member community from the Critical Assessment of MetaProteome Investigation (CAMPI) [18]. We then apply it to two clinical studies, benchmarking public-sequence resources against matched metagenomes in PREDICT, a cohort of patients with acutely decompensated cirrhosis [19], and CapScan, which sampled intestinal fluid with ingestible capsules alongside stool [20,21]. Beyond increasing sequence coverage, these analyses reveal how microbial functions vary between individuals and along the gastrointestinal tract. The compact search spaces further make non-specific searches practical, enabling us to investigate non-tryptic peptide termini directly in the gut.

## Results

### De novo peptide lookup selects compact search databases from broad protein collections

We first tested the FastaLake concept in a healthy-stool variability series of 58 liquid chromatography (LC)– MS/MS acquisitions spanning repeated injections, independent preparations, and sampling within and between individuals. The starting reservoir contained 233.5 million protein sequences from the Unified Human Gastrointestinal Genome (UHGG), the gut subsets of the Global Microbial Gene Catalog (GMGC) and the Global Microbial smORFs Catalog (GMSC) [3–5], together with human Swiss-Prot [22]. We retained all supplied distinct sequences without sequence-similarity clustering. This corresponds to a search space more than 10,000-fold larger than the human reference proteome.

Pooled across the 58 acquisitions, AlphaNovo predicted approximately 250,000 distinct peptides, treating isobaric isoleucine and leucine (I/L) as equivalent, and exact lookup of these peptides retrieved 18 million study-supported proteins from the reservoir, from which the acquisition-specific databases were selected. In a representative acquisition, 25,000 queries retrieved 6.7 million candidate proteins. Deterministic parsimony then assigned each query to one compatible protein using a fixed evidence-based ranking and retained only the proteins receiving evidence (Methods). This reduced the candidates to 12,600 complete sequences, approximately 18,500-fold fewer than the starting reservoir. A conventional database search against this compact sequence set identified 29,700 target peptides of 5–50 residues at peptide *q*≤0.01 (Fig. 1a). We use peptide *q*≤0.01 for database searches throughout.

**Figure 1.**
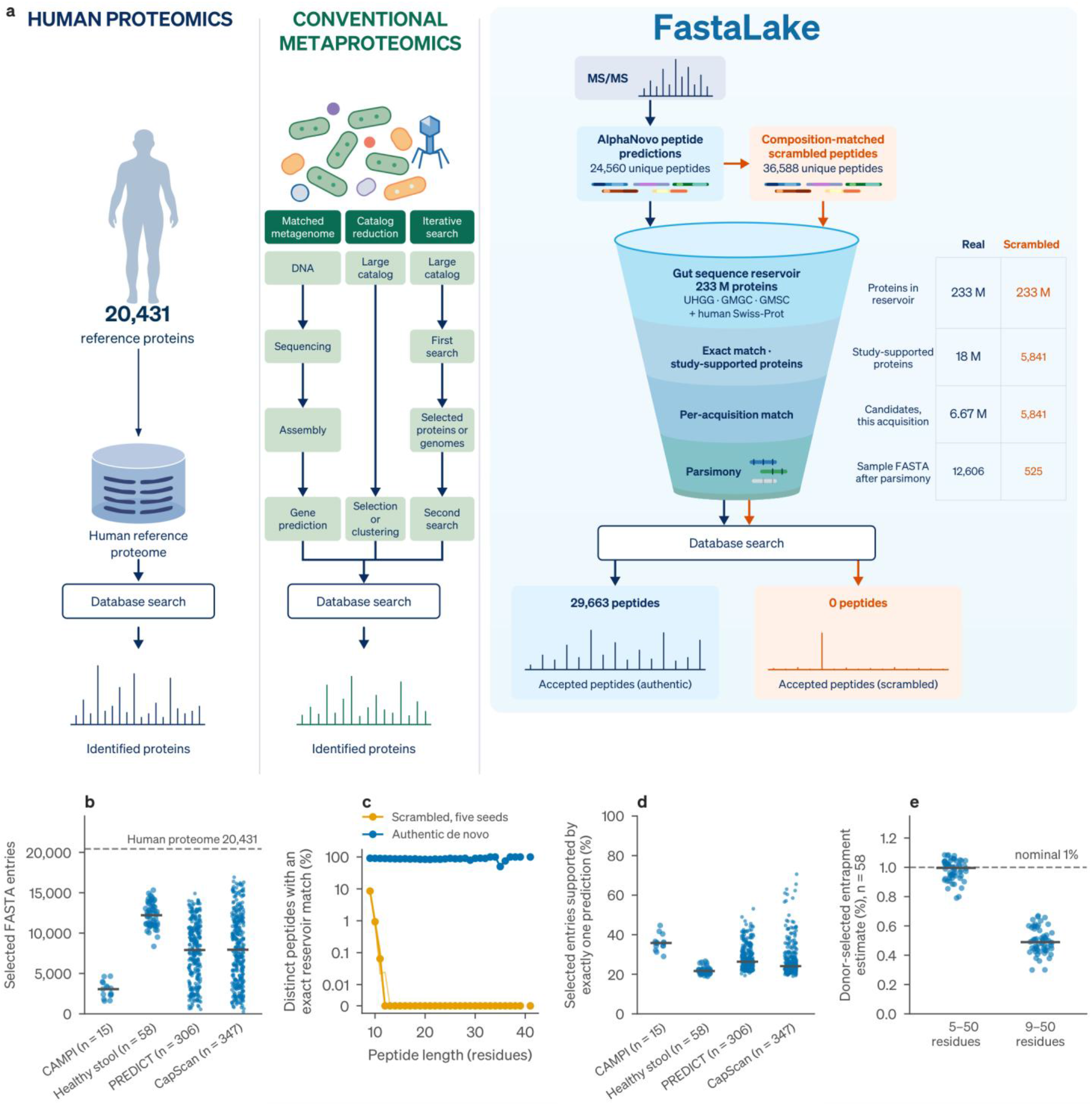
Exact peptide lookup selects compact search databases. **a**, Database strategies. Human proteomics searches one reference proteome (20,431 proteins). Conventional metaproteomics builds sample-specific databases from matched metagenomes, by reducing or clustering large catalogs, or by iterative two-stage searches. FastaLake pools de novo peptide queries from the 58 healthy-stool acquisitions to look up study-supported proteins in the gut sequence reservoir, then selects an acquisition-specific FASTA by exact matching and parsimony, shown for one acquisition. Each of the 36,589 predictions retained for this acquisition was also scrambled with its amino-acid composition kept, giving 36,588 distinct scrambled queries (24,560 distinct authentic peptides). The scrambled queries followed the same lookup, parsimony and search without a study-level pool, so their study-supported proteins and candidates are the same 5,841 proteins. **b**, Selected FASTA sizes across 726 acquisitions. Dashed line, human reference proteome. **c**, Fraction of distinct authentic or scrambled queries with an exact reservoir match, by peptide length (five scrambling seeds; split logarithmic–linear axis). **d**, Fraction of selected proteins supported by exactly one distinct production query. **e**, Donor-selected entrapment estimates in 58 healthy-stool acquisitions for accepted peptides of 5–50 or 9–50 residues. Points are acquisitions and bars show medians.

This reduction was consistent across the datasets studied in this paper. All 726 acquisition-specific databases contained fewer entries than the 20,431-entry human reference proteome, with median sizes ranging from 3,100 proteins in CAMPI to 12,200 in healthy stool (Fig. 1b), despite full study reservoirs containing 233.5 million sequences for healthy stool, 272.9 million for PREDICT, 264.9 million for CapScan and 375.3 million for CAMPI (Supplementary Fig. 1). The PREDICT and CapScan study reservoirs included cohort assemblies, whereas their public-sequence benchmarks used the 233.5-million-sequence reservoir. The CAMPI reservoir additionally included the complete Swiss-Prot and TrEMBL sections of UniProtKB.

To assess how often peptide sequences matched reservoir proteins by chance, we scrambled the amino-acid order within the selected AlphaNovo peptides while preserving their lengths and amino-acid compositions. The fraction of scrambled queries matching a reservoir protein fell from 8.1–8.9% for 9-residue queries to 0.7– 1.1% at 10 residues, with no match in 86,000 query–seed evaluations at ≥13 residues (Fig. 1c, Supplementary Fig. 2). Scrambled queries selected 525 proteins but yielded no accepted target peptides in the final Sage search, compared with 12,600 proteins and 29,700 peptides of 5–50 residues for authentic predictions (Fig. 1a, Supplementary Methods 6). The same held for scrambles that kept the C-terminal residue fixed, in all 10 control workflows (five seeds per design; Supplementary Methods 6 and Source Data).

Across cohorts, a fifth to over a third of selected proteins were supported by a single production query (Fig. 1d), but nearly all (96–98%) contained at least one predicted peptide that was also identified in the final search. The selected databases also supported identifications beyond the predictions used to build them: in the representative acquisition, 14% of accepted peptides had not been predicted by AlphaNovo (Source Data).

We first estimated the error rate of the search step with donor-selected entrapment. Each healthy-stool acquisition was searched against a database selected from another acquisition, so that selection did not use the searched spectra, and supplemented with shuffled entrapment peptides (Methods). The median entrapment estimate was 1.0% for accepted 5–50-residue peptides and 0.49% for the 9–50-residue subset (Fig. 1e). Unless stated otherwise, peptide counts and identities below refer to accepted peptides of 9–50 residues, a subset of the 5–50-residue search results (Methods). In five acquisitions for which the database was instead selected from the same spectra used for identification, the corresponding median estimates were 1.03% and 0.51% (Source Data). Because entrapment peptides were added after selection, these estimates do not capture selection from the searched spectra, which the fusion-entrapment and simulation controls below address.

AlphaNovo also records unresolved local residue order as bags, groups with a predicted composition but no fixed order. An exploratory bag-aware retrieval route gave small net gains or losses in peptide identifications depending on the orderings admitted, and gained identifications were rarely supported by a fragment ion distinguishing the two orders (Supplementary Fig. 2e,f). Production analyses used exact peptide lookup.

### FastaLake recovers a defined microbial community without prior knowledge of its composition

To test whether FastaLake selects a database that reflects the sample’s composition, we used 15 acquisitions from the CAMPI benchmark of the simplified human intestinal microbiota (SIHUMIx), a defined eight-member community [18]. Community-member identities, reference proteomes and specimen-matched metagenomic proteins were not used for database selection. Instead, FastaLake selected proteins from a 375.3-million-sequence reservoir combining broad gut catalogs with UniProtKB to include sequences from cultured isolates [22] (Fig. 2a), which contained 91–100% of each supplied member reference proteome based on exact I/L-equivalent sequence matching (Source Data). We searched the supplied community reference database and the multi-omic co-assembly proteins (hereafter MG) with identical settings.

**Figure 2.**
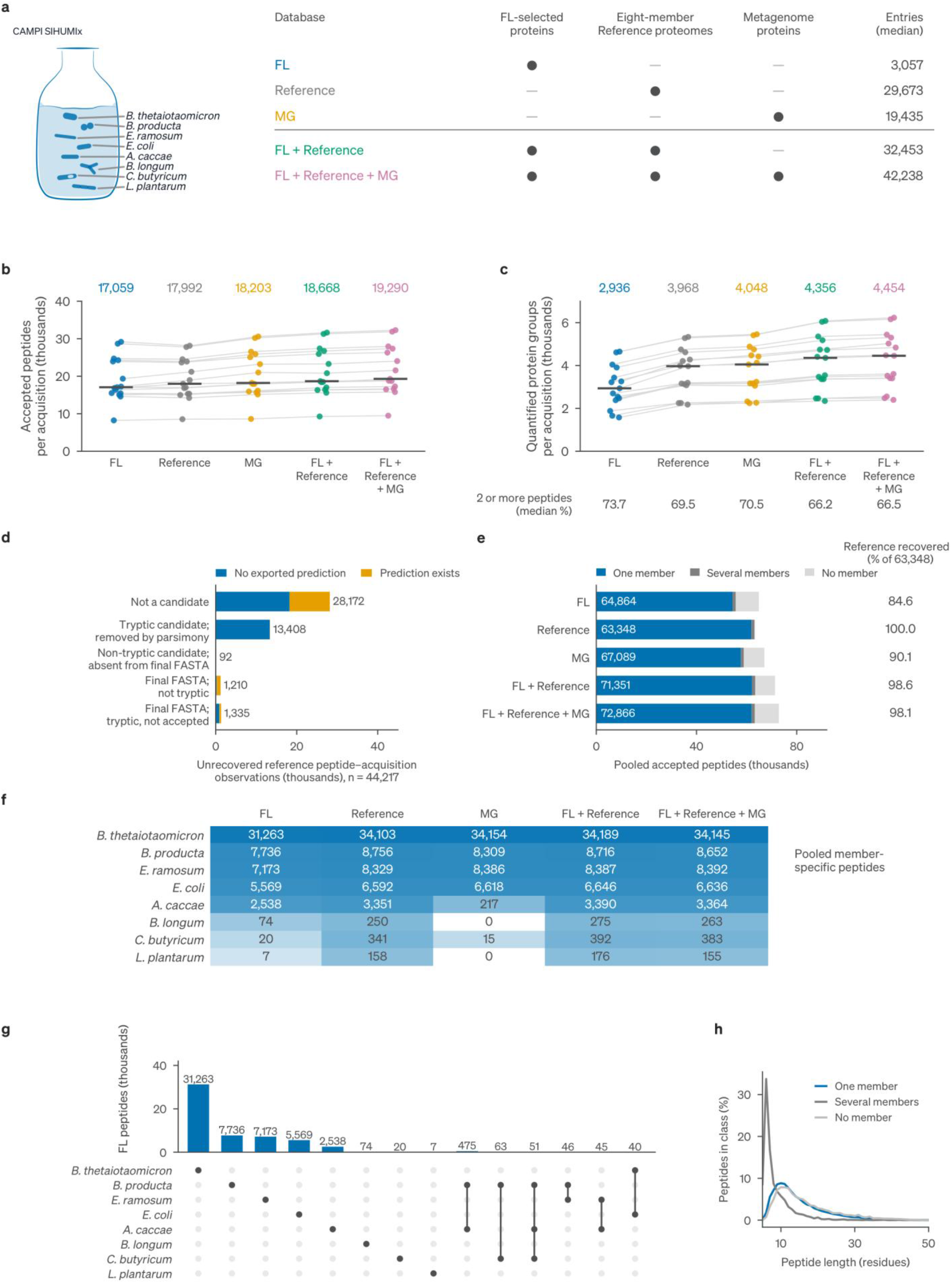
Recovery of the defined CAMPI community. **a**, Fifteen CAMPI SIHUMIx acquisitions and five search databases: FastaLake (FL), supplied reference proteomes (Reference), multiomic co-assembly proteins (MG), and FL with the reference proteomes appended alone or with MG. **b,c**, Accepted peptides (**b**) and quantified protein groups (**c**). Points are acquisitions, lines pair databases and bars show medians. Percentages in **c** give groups with ≥2 assigned peptides. **d**, Stage at which reference peptide–acquisition observations missed by FL were lost, split by whether a de novo prediction existed for the spectrum. **e**, Pooled peptides assigned to one, several or none of the eight members. Right, recovery of the 63,348 pooled reference peptide identities. **f**, Pooled member-specific peptides per member and database. **g**, Sharing of FL peptides among members. **h**, Peptide lengths by member-assignment class. Protein groups in c and peptide lengths in h use all accepted 5–50-residue peptides; all peptides are at peptide *q*≤0.01.

FastaLake produced substantially smaller databases while approaching the peptide yield of the reference search. The selected databases contained a median 3,100 protein entries, compared with 29,700 for the reference database and 19,400 for MG, yet identified a median 17,100 peptides per acquisition, compared with 18,000 and 18,200, respectively (Fig. 2a,b). Paired by acquisition, FastaLake identified as many peptides as the reference database (median ratio 1.01, more in 9 of 15 acquisitions) and 0.95 times as many as MG (fewer in all 15). At the protein level, FastaLake quantified fewer groups, with a median 2,936 per acquisition compared with 3,968 for the reference database and 4,048 for MG (Fig. 2c). Adding the complete reference proteomes to the selected FastaLake databases after parsimony increased this to 4,356 groups, and additionally including MG increased it to 4,454. Both augmented databases exceeded the reference database and MG alone in all 15 acquisitions. Most of this gain came from missing sequences. Of the additional accepted 5–50-residue peptide observations, 98.4% could not have been identified from the FastaLake database before the reference proteomes were added.

Similar peptide counts from FastaLake and the reference database did not imply recovery of identical peptide sequences. FastaLake recovered a median 85% of reference peptide identities within acquisitions and 84.6% of the 63,348 pooled reference identities. Most losses arose during database selection. Of 44,217 unrecovered reference peptide–acquisition observations, 94.2% lacked the sequence in the selected database (Fig. 2d). Appending reference proteins after selection restored pooled recovery to 98.6% (Fig. 2e). Adding MG proteins increased total peptide yield but did not further improve recovery of reference identities (98.1%).

For identified CAMPI spectra, the top-ranked AlphaNovo prediction exactly matched the reference-database identification for 51.8% of 801,613 spectra (per-acquisition median 67.7%, range 26.3–79.2%). Among the 271,469 spectra whose predictions passed the top-30% score filter used for database selection, this increased to 85.3%. Retaining the correct sequence in the selected database did not require an accurate prediction from the same spectrum. Among 40,097 reference peptide observations for which the retained de novo prediction disagreed with the reference identification, 84.8% (34,019) remained searchable in the selected database. Only 20.3% of these discrepant predictions directly retrieved candidate proteins, indicating that most of the corresponding sequences were retained through predictions from other spectra (Supplementary Methods 6).

FastaLake recovered member-specific peptides for all eight community members, whereas MG recovered member-specific peptides for only six, missing *Lactiplantibacillus plantarum* and *Bifidobacterium longum* (Fig. 2f, Supplementary Figs. 3 and 4d). For four of the five most represented members, FastaLake recovered 84– 92% of reference-search peptides and 84–93% of MG peptides. For *Anaerostipes caccae*, which MG largely missed, FastaLake recovered 2,538 member-specific peptides (76% of reference), versus 217 with MG. Recovery was lower for the three least represented members but remained at least as high as MG and increased to reference levels when the reference proteomes were appended (Source Data). Overall, 84% of FastaLake peptides mapped uniquely to one community member, while shared peptides were shorter than member-specific peptides (median 7 versus 13 residues among 5–50-residue peptides; Fig. 2g,h).

A second search engine gave the same picture. With MSFragger and Percolator [23,24], FastaLake yielded as many peptides as the reference database (median ratio 1.00) and 0.94 times as many as MG, while recovering a median 85.3% of reference and MG peptide identities (Supplementary Fig. 3d and Source Data). In healthy stool, MSFragger and Sage identified similar numbers of peptides (median ratio 1.00; Source Data).

Clustering candidate proteins before parsimony consistently reduced peptide identifications. Across all 15 CAMPI acquisitions, MMseqs2/Linclust clustering [25,26] reduced accepted peptide counts by a median 0.9– 8.8% (5–50-residue peptides) as the identity threshold decreased from 99% to 50%, while retaining 96.4– 99.0% of annotated KEGG functions (Supplementary Table 1 and Source Data).

In a CAMPI fusion-entrapment control [27], every reservoir protein was paired with a shuffled control that stayed linked to it through selection and parsimony. At peptide *q*≤0.01, the pooled diagnostic averaged 1.6% for 9–50-residue peptides and 2.7% for 5–50 residues over two randomizations, above the nominal 1%. It was highest at 5–8 residues, mostly because of unresolved target–control pairing (Supplementary Fig. 5 and Supplementary Methods 6).

We also simulated spectra from known CAMPI reference peptides to directly measure false identifications among accepted 5–50-residue peptides. Selecting the database from the searched spectra did not increase the false-discovery proportion under baseline conditions (1.01% versus 1.25% with a fixed reference). Under more stringent stress conditions, including increased noise and removal of half the generating peptides from the reference, the proportion increased to 1.91%, 0.71 percentage points above selection from an independent replicate (Supplementary Fig. 5 and Supplementary Methods 6).

### Selected databases transfer across replicates and can be pooled

The 58 healthy-stool acquisitions form five series of increasing biological variability, from repeated injections of the same sample to samples from different individuals (Fig. 3a). Within each series, we compared protein-group and functional profiles across acquisitions searched with their respective FastaLake databases. Among technical replicates, searching an acquisition against a database selected from another injection yielded a median 96.0% of the peptide count and retained 89.6% of the peptide identities obtained with its own database. Transfer was lower between individuals, retaining 55.8% of the peptide count and 50.2% of peptide identities (Fig. 3b,c).

**Figure 3.**
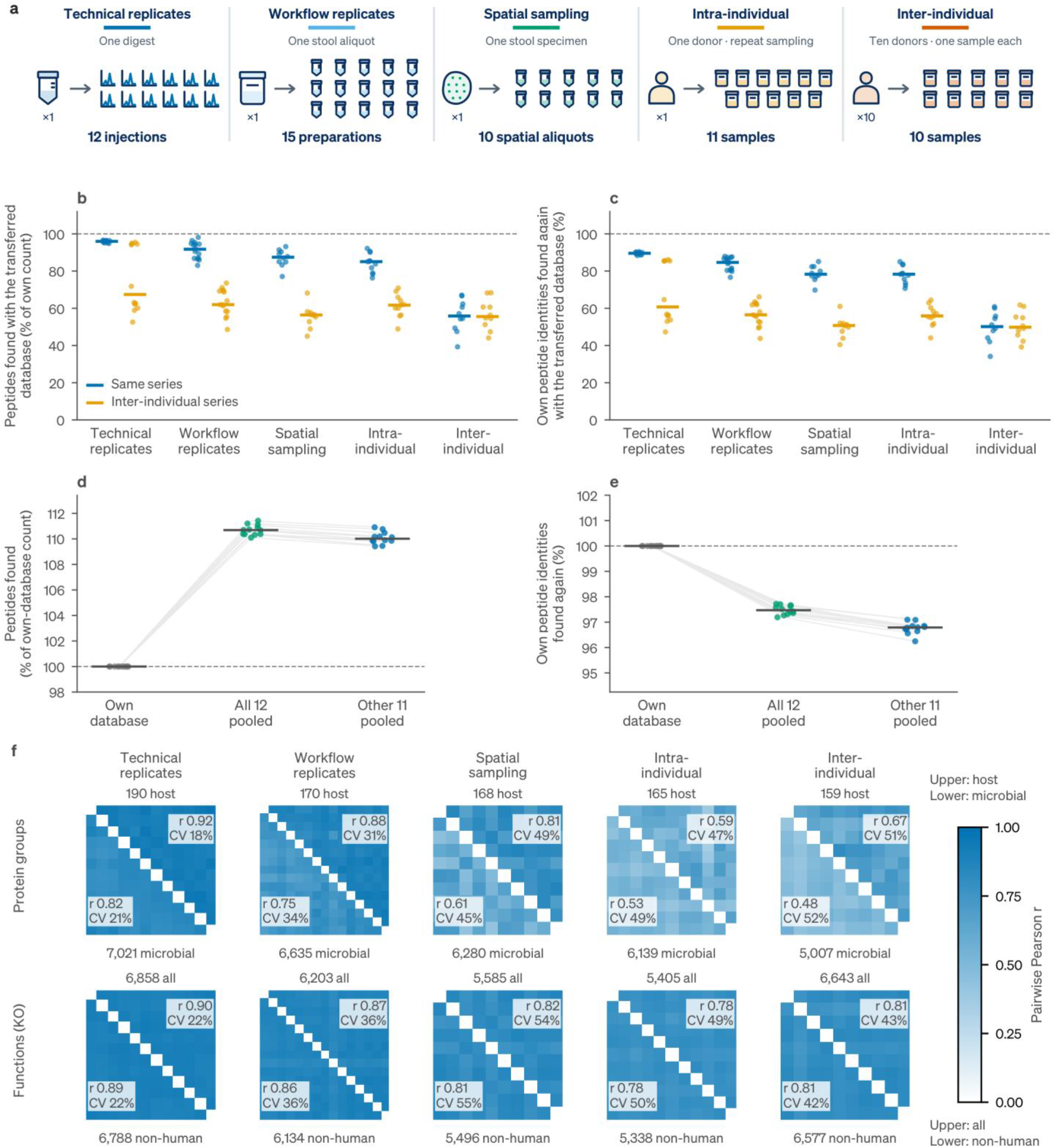
Database transfer, pooling, and quantitative agreement. **a**, Sampling design of the five healthy-stool series. **b,c**, Peptide counts (b) and recovery of peptide identities (c) with a database selected from another acquisition of the same series (blue) or from the inter-individual series (orange), relative to the own-database result. **d,e**, Relative peptide count (d) and recovery of own-database identities (e) for technical replicates searched against their own database, the union of all 12 replicate databases, or the union of the other 11. Points, acquisitions; lines, pairs; bars, medians; dashed lines, own-database result. **f**, Pairwise agreement across the five series for protein-group intensities (host above, microbial below) and KEGG-ortholog intensities (all origins above, human-assigned signal excluded below), using all accepted 5–50-residue peptides. Labels give median Pearson correlation, raw-intensity coefficient of variation (CV) and jointly quantified features per acquisition pair.

To test how much identification depended on using the same spectra for database selection and scoring, we selected protein databases using all predictions, odd-scan predictions or even-scan predictions, and searched the same odd-scan spectra against each database. Across 726 acquisitions, the median ratios of peptide counts with disjoint selection to those with same-half selection were 92.4% in CAMPI, 82.9% in healthy stool, 86.5% in PREDICT and 79.9% in CapScan. Disjoint selection yielded fewer accepted peptides in every acquisition. Cohort medians for recovery of the same peptide identities were lower, ranging from 70.0% to 83.7% (Supplementary Fig. 6 and Source Data).

Given this transfer across related measurements, we asked whether pooling selected sequence spaces could increase peptide recovery. Across the 12 technical replicates, searching each acquisition against the union of databases selected from the other 11 increased peptide identifications in every acquisition by a median 10.0%, while retaining 96.8% of identities from the acquisition-specific search (Fig. 3d,e). Including the acquisition’s own database changed this little (+10.7%, 97.5% retained), indicating that a single database pooled across technical replicates performed similarly to unions that excluded the searched acquisition.

We also used the acquisition-specific searches to compare quantitative variability across the five healthy-stool series. For microbial protein groups, median pairwise coefficient of variation (CV) increased from 21% between technical replicates to 52% between individuals, while Pearson correlation decreased from 0.82 to 0.48 (Fig. 3f). KEGG ortholog (KO) profiles built from all accepted 5–50-residue peptides were more strongly correlated than microbial protein-group profiles in every sampling design, with Pearson correlations of 0.89 between technical replicates and 0.81 between individuals with human-assigned signal excluded (Fig. 3f).

We then asked whether evidence could be combined at the study level with a common protein database. Study-level protein grouping aligned peptide quantities under shared representative accessions, but these accessions were often absent from the FASTA of the acquisition in which the peptide was quantified (Supplementary Methods 7). We therefore merged the 58 acquisition-specific FASTAs and removed exact sequence duplicates, giving a common study database of 153,267 proteins. We searched each acquisition separately against this database, and we also processed all 58 acquisitions jointly in Sage, combining peptide–spectrum match (PSM) rescoring, peptide-confidence estimation and cross-run label-free quantification (LFQ). In the joint search, peptide confidence was estimated either within each acquisition (per-acquisition confidence, used below unless stated) or once across all 58 acquisitions (study-level confidence).

Relative to acquisition-specific searches, the common FASTA increased median peptide counts by 11.4% in separate searches and 11.6% in the joint search, and both retained about 94% of the original peptide identities (Supplementary Fig. 7a and Source Data). Among the 12 technical replicates, joint processing increased the number of peptide features identified in all of them from 10,493 to 13,149, counting only features with an accepted identification (Methods). Study-level confidence raised the median gain to 34.2% on the identical sequence space, so this additional gain reflects how confidence was estimated, not the database, and is not at the per-acquisition 1% peptide FDR used elsewhere.

Joint processing improved quantitative agreement between technical replicates but not between independent preparations, and it did not improve agreement in the leading species assigned to each KEGG ortholog (Supplementary Fig. 7c,d, Supplementary Methods 7 and Source Data).

### Public-sequence FastaLake approaches matched metagenomes in stool and exceeds them in intestinal fluid

We compared public-sequence FastaLake with specimen-matched metagenome databases in two contrasting clinical settings. The PREDICT cohort comprised 306 stool LC–MS/MS acquisitions from patients with decompensated cirrhosis or acute-on-chronic liver failure [19,28], with companion DNA, RNA and metabolite measurements (Fig. 4a). A second cohort comprised sequential intestinal fluid samples collected from healthy participants using CapScan ingestible sampling capsules, together with matched stool [20,21]. The four intestinal sampling regions (regions 1–4) are device-defined and correspond on average to increasingly distal locations. FastaLake selected only from public-sequence resources for these benchmarks. Both search strategies used identical identification settings and independent peptide *q*≤0.01 thresholds.

**Figure 4.**
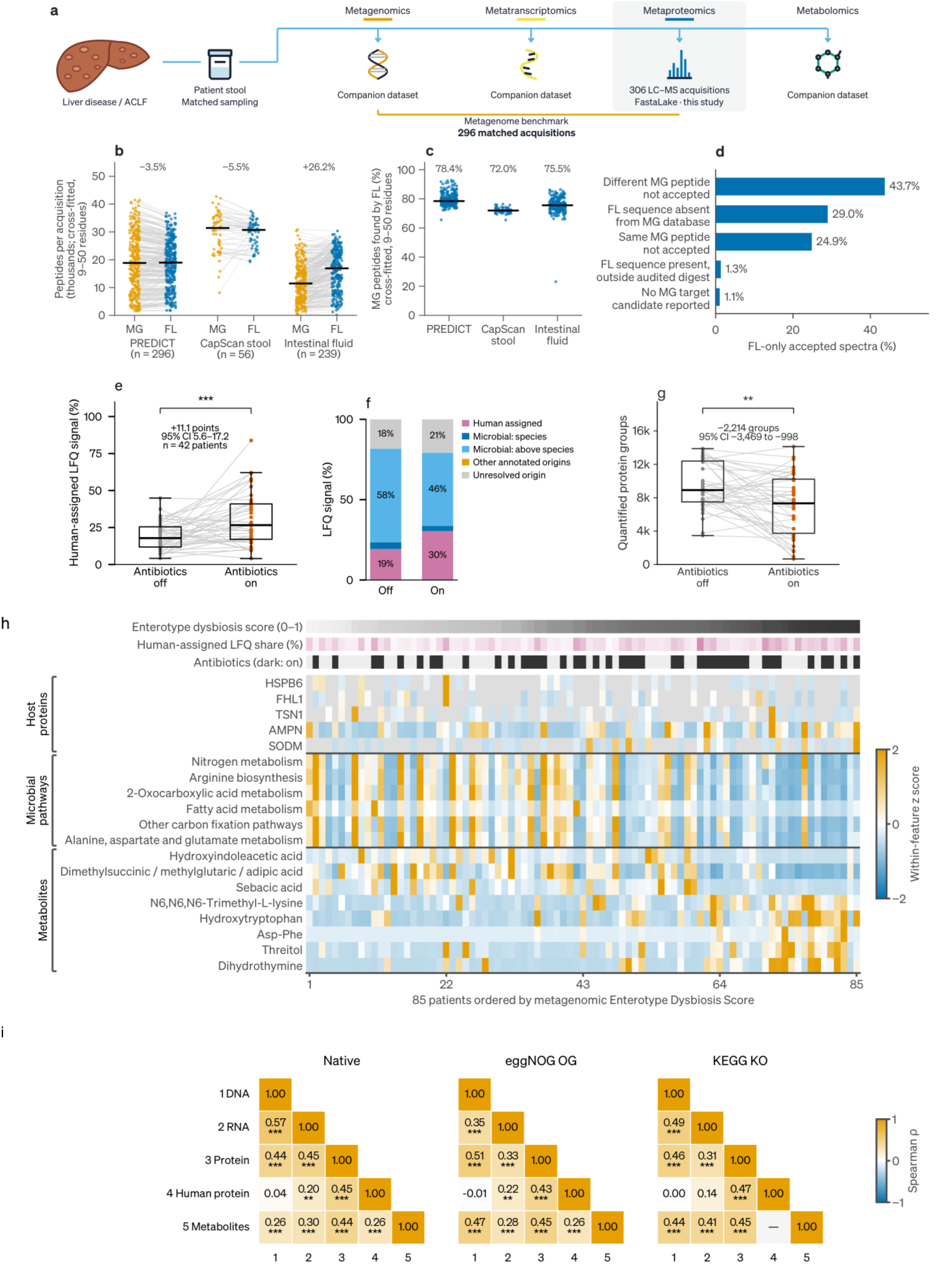
Public-sequence benchmarking and PREDICT molecular profiles. **a**, PREDICT study design and matched molecular measurements. **b,c**, Peptides per acquisition (b) and fraction of metagenome-search peptide identities recovered by FastaLake in the same acquisition (c) in matched metagenome (MG) and FastaLake (FL) searches with 10-fold spectrum cross-fitting, in which each spectrum was searched against a FastaLake database selected from the predictions of the other nine folds (296 PREDICT, 56 CapScan stool and 239 CapScan intestinal-fluid acquisitions; values with databases selected from the scored spectra in Supplementary Fig. 6d,e). Numbers above the points in b are median paired changes in peptides per acquisition (FL relative to MG), as reported in the text. **d**, Sequence availability and MG-search outcomes for 6.69 million PREDICT spectra assigned 5–50-residue peptides only by FastaLake, using same-spectrum database selection and the highest-scoring target among up to five reported candidates (Supplementary Methods 6). **e–g**, Human-assigned LFQ share (e), conditional origin composition (f) and protein-group depth (g) in 42 patients off and on antibiotics. Boxes, median and interquartile range (IQR); whiskers, 1.5 IQR. Numbers in e,g give mean paired differences and 95% CIs. **h**, Host proteins, microbial KEGG pathways and metabolites across 85 patients ordered by metagenomic Enterotype Dysbiosis Score (EDS); values are within-feature z scores. Alternative identities of the ambiguous organic-acid feature (dimethylsuccinic acid) are retained. **i**, Patient-profile distance correlations (77 patients) for native, eggNOG orthologous group (OG) and KEGG ortholog (KO) features; the human protein–metabolite comparison is tested once, under OG (—, KO). Lines pair acquisitions (b) or patient states (e,g); bars, medians. Stars (e,g,i), \**q*<0.05, \*\**q*<0.01, \*\*\**q*<0.001.

FastaLake databases are selected from the spectra that are subsequently searched, whereas metagenome databases are not. Selecting from one half of each acquisition and scoring the other half reduced FastaLake peptide counts by a median 13.5% in PREDICT and 20.1% in CapScan relative to same-half selection (Supplementary Fig. 6). We therefore compared the two databases on spectra not used for FastaLake selection, using 10-fold cross-fitting: the FastaLake database for each fold was selected from predictions in the other nine folds, and the held-out spectra were searched against this database and against the matched metagenome database (Methods). Repeated observations of a peptide in other folds could still support its selection (Supplementary Methods 7).

With confidence estimated jointly across folds, FastaLake identified 3.5% fewer peptides than the metagenome search in PREDICT (95% CI −4.8 to −1.4%) and 5.5% fewer in CapScan stool (95% CI −8.4 to −2.3%). In intestinal fluid, however, FastaLake identified 26.2% more peptides (95% CI 17.0–34.3%), with gains of 20–35% across the four intestinal sampling regions. FastaLake recovered a median 78.4% (PREDICT), 72.0% (CapScan stool) and 75.5% (intestinal fluid) of metagenome-search peptide identities (Fig. 4b,c and Source Data). In a linear mixed model with participant as a random intercept, the FastaLake/metagenome peptide-count ratio was 1.37-fold higher in intestinal fluid than in stool (95% CI 1.20– 1.55, *P*=2 × 10⁻⁶), with no detectable difference among the four intestinal regions (*P*=0.59).

To distinguish sequence availability from database selection, we also applied FastaLake selection to each PREDICT specimen metagenome rather than the public-sequence reservoir. In this same-spectrum comparison, these curated specimen-metagenome databases identified a median 1.5% fewer peptides than the unselected metagenome search (fewer in 177 of 296 acquisitions), whereas public-sequence FastaLake exceeded the selected specimen-metagenome databases by a median 10.6%, with higher peptide counts in 292 of 296 acquisitions (Supplementary Fig. 8a). Selection alone therefore did not improve the median yield from specimen metagenomes, whereas public-sequence FastaLake yielded more peptides than the curated metagenome databases.

Selecting the FastaLake database from the scored spectra themselves gave larger gains in every acquisition: a median 9.4% more peptides than the metagenome search in PREDICT and 40.1% in CapScan (14.0% in stool and 46–62% across the four intestinal regions), with the same stool–intestinal-fluid pattern (Supplementary Fig. 6d,e and Source Data). With a minimum peptide length of seven instead of five, the gains for all accepted peptides were 14.5% and 43.8%, compared with 18.8% and 49.3% for all accepted 5–50-residue peptides (Supplementary Fig. 4a). Much of the 5–50-residue gain came from 5–8-residue peptides, the lengths with the highest fusion diagnostic (Supplementary Fig. 5). Using the same-spectrum FASTAs, MSFragger and Percolator gave median FastaLake/metagenome ratios of 1.05 in 291 PREDICT acquisitions and 1.35 in 295 CapScan acquisitions, compared with 1.10 and 1.40 with Sage on the same acquisition subsets. In the 5–50-residue same-spectrum sensitivity, additionally requiring PSM posterior error probability (PEP)≤0.01 retained a median 85.7% of FastaLake peptides in PREDICT and 84.7% in CapScan and increased the cohort-median same-spectrum gains from 18.8% to 23.5% in PREDICT and from 49.3% to 59.9% in CapScan (Supplementary Methods 4).

Intestinal-fluid samples yielded less DNA than stool (median 54–168 ng across regions versus 687 ng; 99 acquisitions with recorded quantities) and smaller metagenome databases (median 105,000 versus 272,000 predicted proteins). All 15 participants with databases from both compartments had smaller intestinal-fluid databases. The median intestinal-fluid peptide yield was 36% of the median stool yield with metagenome searching and 54% with same-spectrum public-sequence FastaLake (239 and 56 acquisitions). Same-spectrum FastaLake gains were 70%, 53% and 42% across increasing thirds of intestinal-fluid metagenome database size (Supplementary Fig. 9 and Source Data).

In PREDICT, FastaLake-only identifications were not explained solely by sequences absent from the specimen-matched metagenome databases. Among 6.69 million PREDICT spectra accepted only with public-sequence FastaLake, the assigned peptide sequence was absent from the metagenome database in 29%. In approximately 69% (43.7% and 24.9% in the two categories of Fig. 4d in which a metagenome peptide was not accepted), the sequence was present in the allowed theoretical digest, but the highest-scoring reported metagenome target failed the peptide-level threshold (Fig. 4d).

### The cirrhotic metaproteome captures holobiont shifts and multi-omics layer agreement

We next characterized host and microbial changes in the cirrhotic metaproteome and their relationship to DNA, RNA and metabolite profiles from the same patients. For these analyses, we added proteins from the cohort metagenome assemblies to the public-sequence reservoir before FastaLake selection (assembly-inclusive databases). In 42 patients sampled both off and on antibiotics, the human-assigned share of total LFQ intensity was on average 11.1 percentage points higher during exposure (95% CI 5.6–17.2, Benjamini– Hochberg *q*=6.8 × 10⁻⁴). The number of quantified protein groups was on average lower by 2,214 (95% CI 998–3,469, *q*=0.0013, Fig. 4e–g). Both differences held when rifaximin for hepatic encephalopathy was counted as antibiotic exposure (40 patients; Supplementary Table 2). Fig. 4h displays host proteins, microbial pathways and metabolites across 85 patients ordered by the metagenomic Enterotype Dysbiosis Score (EDS) [29]. Three metabolite–enzyme pairs selected post hoc from the full exploratory biochemical screen, with their adjustments, are shown in Supplementary Fig. 10 and Supplementary Table 3.

We then asked whether patients with dissimilar protein profiles also differed across other molecular layers. Among 77 patients with complete DNA, RNA, protein and metabolite data, we computed distances between patients within each layer (Hellinger distances for DNA, RNA and protein profiles, and Euclidean distances for standardized log-transformed metabolites) and correlated the resulting distance matrices. Protein distances correlated with DNA, RNA and metabolite distances (Spearman *ρ*=0.44, 0.45 and 0.44, respectively; all permutation *q*<0.001; Fig. 4i), indicating that patients who differed in their protein profiles also tended to differ in the other molecular layers. These profiles excluded human-assigned signal but retained signal of unresolved origin. Agreement persisted after grouping measurements by shared functional annotations, although pathway-level RNA–protein agreement was weaker (*ρ*=0.16, *q*=0.024; Supplementary Fig. 11a and Source Data). This profile-level agreement did not extend to individual functions. Across patients, median DNA–protein and RNA–protein correlations for individual functions were only 0.03–0.08, compared with 0.24– 0.36 for DNA–RNA (Supplementary Table 4). Thus, the abundance of an individual function in DNA or RNA was only weakly predictive of its protein abundance.

### Personal functional profiles persist across intestinal sampling regions

The CapScan study established a technology for noninvasive sampling of the human intestinal environment during normal digestion [20]. Its proteomic analysis was previously limited to host proteins [20]. We used FastaLake to revisit these data and characterize microbial proteins and their annotated functions alongside the host proteome in intestinal fluid. Participants swallowed capsule devices designed to collect fluid at different points during intestinal transit. We analyzed 347 existing LC–MS/MS acquisitions from 308 specimens of 15 healthy participants, comparing four device-defined intestinal sampling regions with stool (Fig. 5a). We asked whether each participant’s characteristic microbial functional profile persisted across these sampling regions, and whether the same organisms contributed those functions in each region.

**Figure 5.**
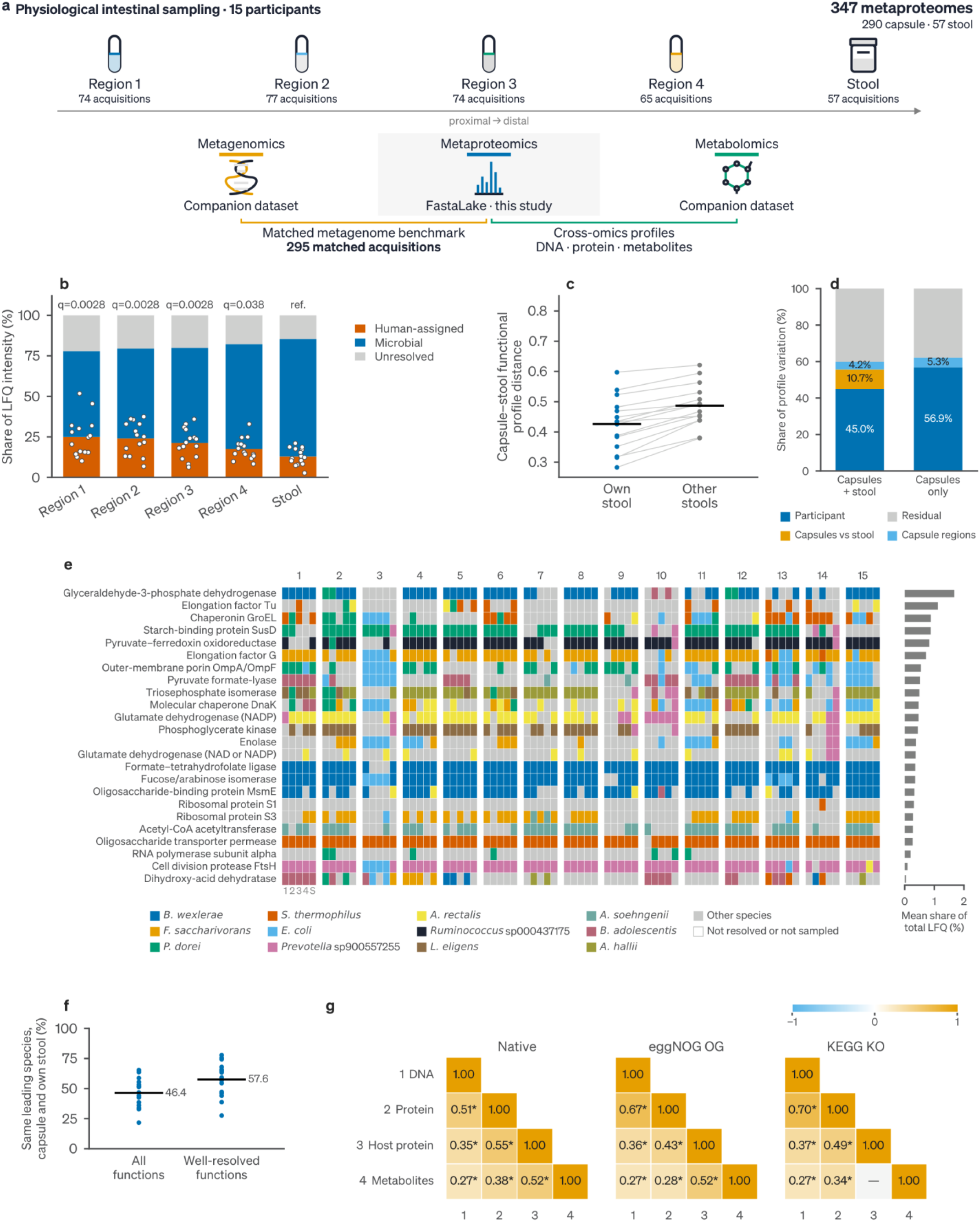
Personal functional profiles across intestinal sampling regions. **a**, Study design: four sequential intestinal sampling regions and stool, with matched data layers. **b**, Human-assigned, microbial and unresolved-origin shares of LFQ intensity (15 participants). Points, each participant’s human-assigned share; *q* values, each region versus the same participant’s stool. **c**, Distance from each participant’s intestinal functional profiles to own versus other stools (73,316 microbial eggNOG orthologous groups). Points, participants; lines pair each participant’s own- and other-stool distances; bars, means. **d**, Variance partition of functional profiles (participant, capsule–stool, among-region, residual), with and without stool. **e**, Leading annotated species for 24 selected functions. Columns, regions 1–4 and stool (S) within each participant; right, mean share of total LFQ intensity per function. Gray, other species; white, unresolved or unsampled. **f**, Leading-species agreement between intestinal regions and own stool (direct peptide assignment) for all functions and for well-resolved functions, with at least 20% species-resolved non-host signal and a leading species at least five percentage points ahead. Points, participants; bars, means. **g**, Spearman correlations between distances of region-adjusted native, eggNOG orthologous group (OG) and KEGG ortholog (KO) profiles in 13 participants with complete profiles; the host protein– metabolite comparison is tested once, under OG (—, KO). \**q*<0.05.

Host proteins contributed a larger share of measured protein intensity in all four intestinal sampling regions than in stool. Using assembly-inclusive databases, which add the CapScan metagenome assemblies to the reservoir, the fraction of total LFQ intensity assigned to human proteins averaged 25.0% in region 1, 17.5% in region 4 and 12.8% in stool among 15 participants with measurements in all five sampling groups (regions 1–4 and stool). Regions 1, 2 and 3 had a higher human-assigned share than stool, by 12.1, 11.1 and 8.4 percentage points (95% CI 6.6–18.7, 5.8–16.4 and 4.5–12.5, Benjamini–Hochberg *q*=0.0028 each, Fig. 5b). Region 4 exceeded stool by a smaller 4.7 percentage points (95% CI 0.8–9.1, *q*=0.038).

We then asked whether variation across intestinal sampling regions erased the microbial functional pattern characteristic of each person. For each of the 15 participants, we calculated the Hellinger distance, a standard ecological measure for comparing relative-abundance profiles that ranges from 0 for identical profiles to √2 for completely non-overlapping profiles. The mean distance from each participant’s four intestinal-region profiles to their own stool was smaller than the corresponding mean distance to other participants’ stools (mean difference 0.061, 95% CI 0.047–0.075, *P*=1 × 10⁻⁵, Fig. 5c). Across intestinal fluid and stool, participant identity accounted for 45.0% of functional-profile variation and sampling group for 14.8%. Of the total variation, 10.7% came from the intestinal fluid–stool contrast and 4.2% from differences among intestinal sampling regions. In the separate intestinal-fluid-only analysis, participant identity accounted for 56.9% of variation and sampling region for 5.3% (Fig. 5d).

The same functions were not necessarily carried by the same species (Fig. 5e). For each function, we identified the species contributing most of its signal in each intestinal region and in the participant’s own stool (Methods). Across all eligible functions, the leading species agreed in about half of the comparisons (46%), rising to 58% for functions with good species coverage and a clear leader (Fig. 5f, Supplementary Fig. 12).

For the 24 displayed functions, the same annotated species led in 50.4% of individual intestinal-fluid–own-stool comparisons (726 of 1,440). Requiring the stool leader to also lead in every resolved intestinal sampling region gave agreement in 111 of 360 participant–function combinations (30.8%, Fig. 5e). These comparisons describe only the taxonomically resolved portion of the community, because species labels accounted for a median of only 29.1% of non-host functional intensity in eligible profiles (Supplementary Fig. 12). Single species also dominated more strongly in intestinal fluid: the leading species accounted for a median 23% of the species-resolved signal of an acquisition, compared with 13% in stool (13 of 15 participants; paired Wilcoxon *P*=6 × 10⁻⁴). A standard evenness index gave the same result (Source Data).

We next asked whether intestinal protein profiles agreed with other molecular measurements from the same participants. After removing average differences among the four intestinal regions and stool, we compared distances between profiles within each molecular layer (Hellinger distances, or Euclidean distances for standardized log-transformed metabolites). Across the 13 participants with complete data, non-host protein distances correlated with DNA, host protein and metabolite distances (Spearman *ρ*=0.51, 0.55 and 0.38, respectively; all *q*<0.001; Fig. 5g). These relationships persisted after grouping microbial features by function. Host and non-host protein profiles derive from the same acquisitions, so their agreement includes shared measurement effects. All Hellinger-distance comparisons passed multiple-testing correction, whereas 9 of 24 rank-based comparisons did not, mostly those involving metabolites (Supplementary Fig. 11f and Source Data).

### Compact databases enable non-specific searches and reveal histidine-rich internal termini

Conventional tryptic searches restrict candidate peptides to those expected from tryptic digestion, whereas non-specific searches can identify peptides with non-tryptic termini at the cost of a much larger search space. The compact FastaLake databases allowed such searches of all sequential intestinal-fluid and stool acquisitions. Across all 347 CapScan acquisitions, non-specific searches scored a median of about 20,200 candidates per spectrum, compared with 218 in tryptic searches of the same assembly-inclusive selected FASTAs (median paired increase, 90-fold; Source Data). Peptides ending in neither lysine nor arginine (non-K/R termini) accounted for 5.1–5.6% of peptide sequences in non-specific searches across the four intestinal regions, compared with 3.1% in stool, with higher fractions in intestinal fluid for all 15 paired participants (Fig. 6a). Semi-tryptic searches and de novo predictions showed the same enrichment. By comparison, the conventional tryptic search recovered only 0.34% non-K/R peptides, all at permitted protein termini.

**Figure 6.**
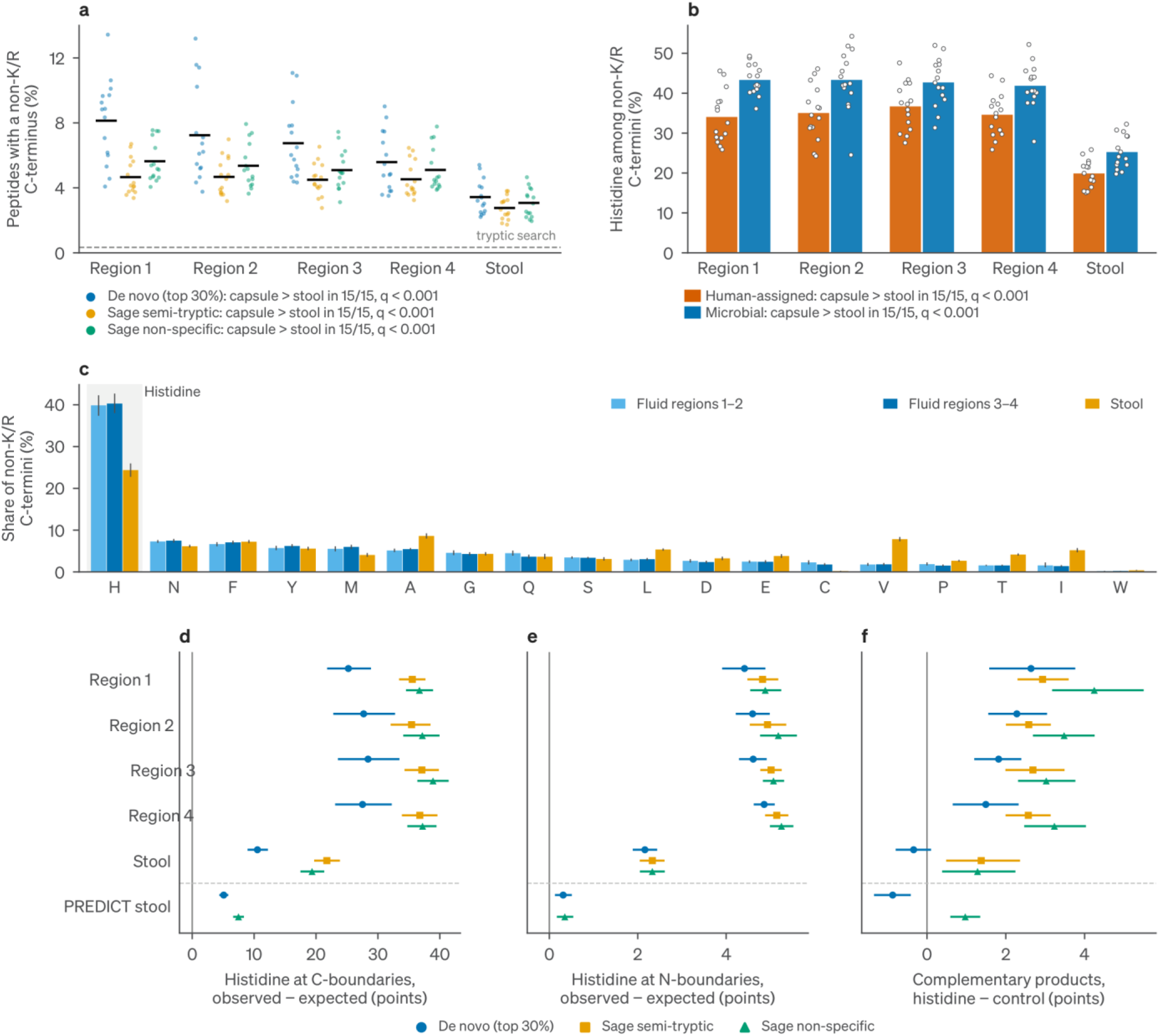
Histidine-rich internal peptide termini in intestinal fluid. **a**, Fraction of peptides with non-K/R C termini in regions 1–4 and stool for de novo, semi-tryptic and non-specific search routes. Dashed line, conventional tryptic-search baseline. **b**, Histidine fraction among non-K/R C termini for human- and microbial-assigned peptides from the non-specific search. **c**, Residue composition of non-K/R C termini in regions 1–2, regions 3–4 and stool. **d,e**, Histidine enrichment at uniquely localized internal C-terminal (d) and N-terminal (e) boundaries relative to the same source proteins, including PREDICT stool (93 patients). **f**, Complementary-product frequency at histidine minus matched non-histidine boundaries. In a,b, points are participants and horizontal lines (a) or bars (b) are means across participants. Bars (c) and symbols (d–f) show participant or patient means with 95% bootstrap intervals. *q* values (a,b) are from paired sign-flip tests of each participant’s mean across the four intestinal regions against their stool (*n*=15), Benjamini– Hochberg adjusted.

Histidine was strongly enriched among these non-K/R C termini. In non-specific searches, histidine accounted for 39.9% and 40.3% of non-K/R C termini in regions 1–2 and regions 3–4, respectively, compared with 24.4% in stool (Fig. 6b,c). We observed the higher histidine share in intestinal fluid than in stool separately among human- and microbial-assigned peptides in all 15 participants, with mean paired differences of 15.2 and 17.6 percentage points, respectively.

This enrichment was not explained by the histidine content of the source proteins. At uniquely localized internal boundaries, approximately 40% of observed non-K/R C-terminal sites in intestinal fluid were histidines in database searches, compared with approximately 2.4% of eligible positions in the same proteins (Fig. 6d). Enrichment was greater at C-terminal than N-terminal boundaries and occurred in every intestinal sampling region (Fig. 6d,e). The C-terminal histidine enrichment was also observed in the independent PREDICT stool cohort: all 93 patients showed positive enrichment with both de novo predictions and non-specific database searches (Supplementary Fig. 13).

We then asked whether peptides terminating at histidine were accompanied by products on the opposite side of the same protein boundary. A complementary product was a second peptide identified from a different spectrum that began at the residue immediately following the terminal histidine. In intestinal fluid, histidine boundaries were more often accompanied by such complementary products than matched non-histidine boundaries across de novo, semi-tryptic and non-specific routes (Fig. 6f). At least some of the histidine-enriched termini are therefore likely to be protein cleavage boundaries. In stool, non-specific searches showed the same pattern, whereas de novo predictions showed no excess of complementary products at histidine boundaries in CapScan stool and fewer in PREDICT stool. Supplementary Figs. 13 and 14 show sequence-context and selected-spectrum checks.

## Discussion

Metaproteomic database construction requires defining a protein sequence space for communities whose taxonomic and strain composition are not known a priori. FastaLake uses de novo peptide sequences from the proteomic measurement to select this search space directly from broad sequence databases. Full-length predictions serve as exact queries for protein retrieval, separating the sequence resource from the database subsequently searched against the spectra. The underlying sequence reservoir can therefore expand without a proportional increase in the size of the database searched against the spectra. Across the datasets studied here, all 726 acquisition-specific databases selected by FastaLake were smaller than the human reference proteome.

This strategy has become possible with recent advances in de novo peptide sequencing. Exact lookup requires full-length predictions that are sufficiently accurate to retrieve the relevant proteins from hundreds of millions of candidate sequences. Earlier workflows instead used de novo output as short sequence tags or as queries for homology search [14,15]. With AlphaNovo, the top-ranked prediction matched the database identification for about half of identified CAMPI spectra and for 85% of those passing the score filter used for selection. Errors in individual spectra were partly compensated by predictions from other spectra of the same proteins, allowing the selected databases to retain most reference peptides even when the prediction for a given spectrum was incorrect. Further improvements in de novo sequencing should improve retrieval, particularly for proteins represented by weak or sparse spectral evidence.

The matched-metagenome comparisons show that sequence coverage alone does not determine search performance. With separate spectra for database selection and peptide identification, public-sequence FastaLake identified slightly fewer peptides of 9–50 residues in stool and substantially more in intestinal fluid than matched metagenome databases. In the same-spectrum PREDICT comparison, most FastaLake-only spectra were assigned peptides already present in the metagenome database but not accepted in that search. Metagenome searches also scored more candidates per spectrum: on identical spectra, metagenome searches scored a median of 6,083 candidates per spectrum in PREDICT and 3,318 in CapScan, compared with 277 and 276 for public-sequence FastaLake. In the same-spectrum curated-metagenome comparison, selection did not improve the median peptide yield, whereas public-sequence FastaLake outperformed the curated databases. Matched metagenomics remains valuable as an independent molecular layer and a source of sequences absent from public catalogs. When sequencing is performed specifically to construct a proteomic search database, our results indicate that public sequence catalogs can replace it for stool and intestinal fluid, with slightly lower peptide yields in stool and higher yields in intestinal fluid, avoiding the added cost and turnaround of specimen-specific sequencing, assembly and gene prediction [30–33].

The difference in database size between public-sequence selection and matched metagenomes grows when databases are combined across a study. Pooled across specimens, matched metagenome databases accumulated 44.8 million distinct protein sequences in PREDICT and 29.2 million in CapScan and continued to grow as specimens were added. The corresponding public-sequence FastaLake databases contained only 0.54 and 0.42 million sequences, 69- to 84-fold fewer, with growth slowing as additional specimens were included. In separate common-FASTA sensitivities using the original clinical rosters, searching each acquisition against a study-wide FastaLake union increased the sequence space by 57- to 70-fold, with median peptide gains across participants below 5%.

These results argue for systematically depositing metagenome-derived proteins in public catalogs such as SPIRE, with 35 billion predicted proteins from 99,146 metagenomes [34]. As catalogs become more complete, the added value of matched sequencing for database construction is likely to decline for well-studied human sample types such as stool and intestinal fluid.

The stool–intestinal-fluid comparison illustrates how metagenome coverage depends on DNA recovery, assembly and gene prediction, as well as on how well the sequenced material represents the organisms contributing proteins to the proteomic sample. Intestinal fluid yielded less DNA and smaller metagenome databases than stool, and the larger FastaLake gain in CapScan was concentrated there. Direct selection from the proteomic measurement may therefore be particularly useful where matched metagenomic sequence coverage is limited or unavailable, including low-biomass or host-rich specimens, anatomical sites from which representative DNA is difficult to obtain, and existing proteomic datasets without accompanying sequencing.

FastaLake differs from existing database reduction strategies in three respects. First, its reservoir retains strain-level sequence variants without similarity clustering, and parsimony chooses among them by peptide evidence. Exact lookup returns complete reservoir proteins, and clustering candidate proteins before parsimony reduced accepted CAMPI peptides across all sequence-identity thresholds tested. Second, selection uses de novo predictions rather than hits of an initial database search. De novo predictions select the database directly, whereas iterative two-stage strategies select proteins from hits of an initial database search, tying selection to its scoring and search space. Third, parsimony prioritizes discriminating sequence evidence among otherwise compatible proteins. In the representative healthy-stool acquisition, accepted peptides occurred in a median of 201 reservoir sequences, and only 771 of 24,430 occurred in a single sequence. Proteins removed during parsimony are not lost to interpretation because identified peptides can subsequently be mapped back to every reservoir protein containing them. The reservoir can also retain sequence classes that may be lost through conventional clustering or gene prediction. Sequences from the Global Microbial smORFs Catalog comprised 3.5–3.8% of selected proteins and supplied 1.3–1.5% of identified peptides that matched no other selected protein, of which 30% (PREDICT) and 49% (CapScan) were absent from the matched metagenomes (Source Data).

The resulting proteomes provide a functional view that is not interchangeable with taxonomic or genomic measurements. In CapScan, microbial functional profiles remained participant-specific along the intestine even when the leading annotated species changed between regions. In PREDICT, DNA, RNA and protein profiles captured similar broad differences between patients but corresponded only weakly at the level of individual functions. These observations are consistent with functional redundancy among microbial taxa and with regulation separating protein abundance from genomic potential. Removing the requirement for specimen-matched sequencing makes this functional layer accessible across samples and sampling sites for which parallel metagenomic measurements are not available.

Compact search spaces also extend the range of feasible peptide searches. Non-specific and open-modification searches greatly increase the number of candidate peptides generated from each protein and become difficult to apply to metagenome-scale databases, particularly once databases are combined across a study. Here, non-specific searching revealed a striking enrichment of histidine at peptide C termini within proteins in intestinal fluid, where histidine accounted for about 40% of these boundaries despite representing only about 2.4% of eligible positions in the source proteins. The enrichment occurred across all intestinal sampling regions and was also detectable in stool and in the independent PREDICT cohort. Histidine preference is rare across the substrate-binding positions surveyed in MEROPS [35]. The reproducibility and magnitude of the sequence preference make its origin an interesting target for controlled experiments varying sample handling and protease activity. More generally, reducing the protein search space makes endogenous proteolysis and non-tryptic peptides accessible to systematic analysis in complex microbial samples.

FastaLake remains constrained by both the sequence content of the reservoir and the de novo evidence used for selection. Proteins absent from the reservoir cannot be retrieved, and proteins with insufficient de novo evidence may not enter the selected database, as seen for the least abundant members of the defined CAMPI community. Selecting the database from the searched spectra also raised peptide counts: disjoint selection yielded a median 7.6–20.1% fewer peptides than same-half selection, the CAMPI fusion diagnostic exceeded the nominal 1%, and selection from the searched measurement increased the simulated false-discovery proportion under stress. The comparison with matched metagenomes therefore rests on cross-fitted searches. Peptide length introduces a related sensitivity–specificity trade-off. Short peptides contributed to recovery of low-abundance organisms but had lower sequence specificity and higher entrapment rates. Evidence can also be combined across related acquisitions to increase depth and quantitative completeness, but the level at which confidence is estimated must remain explicit. Study-level confidence is estimated across the pooled study rather than within individual acquisitions, and gains resulting from this procedure are distinct from gains attributable to database composition.

Neither the sequence reservoir nor the downstream search engine is intrinsic to FastaLake. Database selection here used AlphaNovo, but exact lookup requires only sufficiently specific peptide sequence predictions, allowing other de novo models to supply queries. The reservoir can likewise incorporate additional strain variation, small proteins and sequences from less characterized environments [4,5,36] without clustering this diversity or transferring the full expanded resource into the final search space. Because the output is an ordinary FASTA file, database selection is also independent of downstream search and error-control software, and the same-spectrum comparisons were repeated with MSFragger and Percolator.

Computational requirements remained modest: study-level lookup against reservoirs containing 233–375 million sequences required 8–12 min per cohort on a single large-memory node, while a chunked mode completed lookup against the 233-million-sequence reservoir using less than 11 GiB of memory. AlphaNovo inference, outside these timings, had a recorded median of 28 min per CAMPI acquisition on one GPU. The same separation may extend to data-independent acquisition if de novo reconstruction from multiplexed spectra provides sequences sufficiently specific for exact retrieval.

FastaLake changes the relationship between the size of microbial sequence resources and the search space presented to mass-spectrometric data. Comprehensive catalogs need not themselves serve as search databases. Instead, they can function as persistent sequence reservoirs from which each proteomic measurement retrieves the proteins it supports. Sequence resources can therefore grow in size and strain diversity without imposing the same growth on the searched database, while existing datasets can be revisited as catalogs and de novo models improve. For well-cataloged microbial environments, database construction can thus become a computational operation rather than a requirement for matched specimen sequencing, connecting expanding microbial sequence resources directly to the measurement of expressed microbial proteins.

## Methods

### Study design, samples, and experimental measurements

We analyzed four data-dependent acquisition (DDA) metaproteomic datasets: 15 CAMPI SIHUMIx acquisitions from an eight-member community [18], 58 healthy-stool acquisitions, 306 PREDICT stool acquisitions from patients with liver disease [19,28], and 347 CapScan acquisitions from 308 specimens and 15 participants [20,21]. The healthy-stool series covered technical replication, independent preparation (workflow replicates), spatial aliquots, longitudinal (intra-individual) sampling and inter-individual variation. All 15 CapScan participants contributed intestinal-fluid and stool samples. Sampling regions 1–4 are increasingly distal on average but do not anatomically localize individual samples. Supplementary Methods 1 and Supplementary Data 1 give experimental designs, acquisition-to-specimen links, analysis populations and exclusions.

Healthy-stool and PREDICT samples (50 mg stool) were lysed with PreOmics LYSE buffer, digested with trypsin and Lys-C and analyzed by DDA on a Q Exactive HF-X (preparation and acquisition settings in Supplementary Methods 1).

CAMPI preparation and acquisition followed the source study [18]. CapScan used SDS/heat lysis, sonication, bead-based capture, trypsin/Lys-C digestion and EASY-nLC 1200 separation with an Orbitrap Exploris 480 [20]. CapScan metagenomes were processed with Skewer 0.2.2 [37], Bowtie2 2.4.1 [38], MEGAHIT 1.2.9 [39] and Prodigal 2.6.3 [40]. FastaLake used supplied protein FASTAs without rerunning assembly or gene prediction. Supplementary Methods 1 gives protocols and settings recovered from all 347 CapScan RAW files.

PREDICT stool DNA was extracted and sequenced at EMBL as described [29] (Supplementary Methods 1). Metagenome-derived protein FASTAs were produced with the SPIRE pipeline [34] and supplied by EMBL with companion RNA profiles. CapScan metabolomics followed the source studies [20,21]. Supplementary Methods 1 gives available protocols and separates supplied measurements from downstream analysis. The PREDICT metabolomics protocol will be made available upon publication.

### Ethics and consent

Healthy-stool measurements were performed in house using samples from healthy volunteers. PREDICT samples were obtained from the source study described below and measured in house; CapScan and CAMPI data were obtained from the source studies described below. PREDICT stool samples were collected in the MUCOSA-PREDICT sub-study of the PREDICT study (ClinicalTrials.gov NCT03056612) at the University of Debrecen, Hungary. The study was approved by the local ethics committee (35281-2/2017/EKU), and all patients gave written informed consent in accordance with the Declaration of Helsinki [28]. The CapScan study was approved by the WIRB-Copernicus Group institutional review board (study 1186513), and informed consent was obtained from each participant [20,21]. Healthy-stool samples were donated by 10 healthy volunteers (five female, five male), members of the Department of Proteomics and Signal Transduction of the Max Planck Institute of Biochemistry, after informed consent and with institutional permission from the Max Planck Institute of Biochemistry. The CAMPI SIHUMIx data derive from a defined bacterial community [18].

### Reservoirs, de novo prediction, and exact database selection

Reservoir proteins were cleaned to uppercase letters and deduplicated by exact full-length sequence, preserving I/L distinctions and source provenance without additional sequence-similarity clustering. Source priority was UHGG, GMGC, GMSC, study assemblies and Swiss-Prot, followed by TrEMBL for CAMPI. The public-sequence reservoir contained 233,497,702 proteins from the UHGG protein catalog, gut GMGC, gut GMSC and reviewed human Swiss-Prot [3–5,22]. Healthy-stool analyses and public-sequence clinical benchmarks used this reservoir. Biological analyses added all 296 PREDICT or 335 CapScan assembly FASTAs, giving 272,919,212 and 264,925,283 sequences. Assembly-derived accessions combined a cohort prefix with the sequence SHA-256 digest, making identifiers consistent within each cohort namespace; catalog and human accessions were retained according to source priority. CAMPI used gut resources and complete Swiss-Prot and TrEMBL, UniProtKB 2026_03, totaling 375,333,654 sequences. Its community membership, member references and specimen metagenomes were not used for selection (Supplementary Methods 1 and 3).

Archived top-ranked AlphaNovo predictions, one per spectrum, supplied the unmodified peptide sequences and ranking scores for all database selection. Supplementary Methods 2 specifies the checkpoint, preprocessing, and exported fields.

Lookup queries were positive-scoring predictions of 9–50 residues passing the low-complexity rules in Supplementary Methods 2. Complexity was assessed before I/L collapsing. We retained the top 30% of eligible predictions per acquisition, including score ties, and deduplicated their I/L-equivalent sequences. Exact substring lookup, implemented with Aho–Corasick matching, retrieved complete reservoir proteins containing a retained query, including overlapping matches. Lookup treated I and L as equivalent but preserved original protein sequences and headers in the selected FASTAs. It imposed no enzyme, mass-equivalence or approximate-matching rule. A chunked mode matched complete-record reservoir partitions against the same queries and merged results in reservoir order, preserving selected proteins while bounding memory.

Queries pooled across each study first retrieved a study-supported protein space. Each acquisition’s queries then retrieved its candidates from this space under the same rules, equivalently to querying the full reservoir (Supplementary Methods 2).

Selection parsimony used all eligible acquisition-level predictions mapping to candidate proteins, including those below the top-30% lookup threshold. Unique and total distinct I/L-equivalent evidence counts were computed once against the complete candidate set. Each query was assigned to a compatible protein ranked by highest unique count, highest total count, smallest accession hash and then lexical accession. Proteins receiving evidence were retained in full. This fixed ranking preserves mapped evidence without guaranteeing a minimum protein set or a unique origin for shared peptides. It differs from post-search inference (Supplementary Methods 2).

### Reference databases and matched comparisons

The CAMPI reference and metagenome (MG) databases [18] contained 29,673 and 19,435 records, respectively, each including 116 contaminants. Supplied decoys were removed. The reference included genus-level surrogates for *Blautia producta* and was not an exact generating-sequence inventory. FastaLake (FL) + Reference and FL + Reference + MG appended complete comparator targets after parsimony without further selection. Reservoir coverage of the eight supplied member proteomes was assessed with I/L equivalence (Supplementary Methods 3).

The matched benchmark compared public-sequence FastaLake with each specimen metagenome plus human Swiss-Prot in 296 PREDICT and 295 CapScan acquisition pairs, the acquisitions whose specimen had a supplied metagenome (Supplementary Methods 1). Both arms used identical identification settings, with confidence estimated separately. Peptide gain was the acquisition-level percentage change relative to MG, summarized by the cohort median; identity recovery was the median fraction of MG canonical peptides recovered by FastaLake. Candidate competition used Sage scored-candidate counts on shared rank-one spectra, summarized within acquisitions and then across the cohort. The primary cross-fitted regional comparison used a maximum-likelihood mixed model of log(FLᵢ/MGᵢ), with sampling group fixed and participant random intercepts; Wald tests assessed intestinal fluid versus stool and equality across regions. For the curated specimen-MG comparator, the production top-30% lookup and parsimony selected proteins using predictions from the same acquisition subsequently searched, without LFQ. Supplementary Methods 3 gives rosters, estimator details and comparator provenance.

### Database searching and peptide acceptance

Standard searches used the Sage v0.14.7 release [17], whose executable reports 0.14.6, with K/R cleavage except before proline, two missed cleavages, lengths 5–50 residues and masses 500–5,000 Da. Precursor and fragment tolerances were ±20 ppm. Carbamidomethylation of cysteine was fixed and methionine oxidation variable, with at most two variable modifications. Searches allowed isotope errors −1 to +3, required four matched peaks and reported one peptide–spectrum match (PSM) per spectrum unless specified. Sage generated terminal-preserving reversed decoys. FastaLake FASTAs contained selected reservoir proteins without an appended contaminant library. Five truncated PREDICT mzML inputs were replaced by validated RAW reconversions; conversion versions and analysis-specific exceptions are given in Supplementary Methods 4 and 7.

Identification summaries retained rank-one targets at Sage peptide_q≤0.01. Sage fitted a target–decoy linear discriminant and estimated peptide *q* values by picked target–decoy peptide competition. Confidence was estimated within each search invocation, before modification removal and I/L collapsing defined canonical peptide identities. For joint healthy-stool searches, confidence was estimated over combined inputs and additionally within acquisitions from the same results (Supplementary Methods 7). Spectrum, protein and label-free quantification (LFQ) q values were distinct quantities used only where specified; CAMPI accession-support analyses additionally required protein *q*≤0.01.

The additional-confidence sensitivity required peptide *q*≤0.01 and PSM posterior error probability (PEP)≤0.01 on the same target PSM before canonical peptide union. Missing PEP estimates were excluded only from that sensitivity; *q* values were not re-estimated. Cross-fitted searches used Sage’s per-fold PSM PEP. Peptide *q*≤0.01 remained the primary acceptance rule (Supplementary Methods 4).

Main-text peptide counts use 9–50 residues; indicated figures and Source Data also report 5–50 residues. These counts are a subset of the accepted 5–50 lists without q-value re-estimation. Full-length selected proteins can yield 5–8-residue identifications although these lengths were ineligible for lookup. Pooled identities are set unions; peptide–acquisition observations retain repeats. Study grouping adds no study-wide or protein-group FDR calibration. A minimum-seven-residue sensitivity repeated production, healthy-stool common/joint and CAMPI fusion searches with confidence re-estimated and direct peptide assignment (Supplementary Fig. 4; clinical gains also in Supplementary Methods 7).

### Protein inference, quantification, and annotations

Post-search inference used accepted canonical peptides and their memberships in the searched proteins, separately by study and database strategy. A deterministic greedy cover repeatedly selected the protein explaining most uncovered peptides, with lexical tie-breaking. Each peptide was assigned to the first selected representative explaining it. LFQ features were Sage rows representing modified peptides with charge states combined; acceptance required the canonical peptide at peptide *q*≤0.01 in the same acquisition. Positive finite intensities were assigned once to the representative and summed without normalization, imputation or an LFQ-q filter. Representatives are quantification labels, not unique biological origins. A study-wide cover keeps assignments consistent across acquisitions (Supplementary Methods 5).

Groups were identified within an acquisition by an accepted assigned peptide and quantified by positive finite intensity. Study-level grouping used 58 healthy-stool, 306 PREDICT, 347 CapScan, or 15 CAMPI acquisitions. Matched comparisons used identical 296- or 295-acquisition rosters in each arm.

Peptide features were human-assigned when annotated compatible proteins agreed on *Homo sapiens*; unannotated compatible proteins did not veto assignment. Compatible proteins included the study’s searched proteins and full local UHGG catalog. Microbial, other and unresolved features remained in total-intensity denominators. Fig. 3f and histidine analyses used the distinct group-level and accession-based origin rules in Supplementary Methods 5 and 9. Functional annotations combined eggNOG-mapper 2.1.12 [41], eggNOG 5.0.2 and DIAMOND 2.0.11 [42] with MGnify/UHGG and earlier GMGC/GMSC annotations. MGnify assignments required alignment to the seed ortholog at ≥50% of the reported bitscore. Peptide intensity was split equally among distinct eligible terms; unannotated intensity was retained and total intensity conserved. Pathways excluded map011xx global maps; microbial orthologous group (OG) profiles required exactly one bacterial or archaeal OG.

Functions comprised KEGG orthologs (KOs) [43], microbial OGs and pathways. A species label required peptide-compatible support for both function and species; other signal remained unresolved. The leading annotated contributor was selected with lexical tie-breaking. Leader eligibility and stricter species-resolution and margin criteria are specified in Supplementary Methods 5 and 8, alongside matched comparisons and the separate fixed-dictionary diagnostic.

### Validation, error controls, and spectrum inspection

Composition-preserving scrambled predictions underwent the full selection/search workflow. In a representative healthy-stool acquisition, five seeds measured length-dependent lookup fractions using all eligible distinct queries as denominators, and a terminal-preserving variant repeated the complete workflow. Adjacent swaps, candidate clustering, bag-aware retrieval and mass substitutions were separate sensitivities. MSFragger 4.1 with Percolator 3.06.5 searched all 15 CAMPI, 291 PREDICT, 295 CapScan and 58 healthy-stool acquisitions using the production same-spectrum FASTAs and mzML inputs. Confidence was estimated separately by acquisition and engine. The five reconverted PREDICT inputs failed in this engine and were excluded from paired engine summaries. Supplementary Methods 2 and 6 specify routes, settings, missing searches and fragment-audit criteria.

CAMPI member-specific peptides mapped to exactly one supplied member proteome, excluding contaminants. Pooled and acquisition-level counts retained zeros and both length windows. Recovery was assessed within acquisitions and after set union. Unrecovered reference peptide observations were traced through retrieval, parsimony, compatible context and final acceptance. Same-spectrum prediction accuracy was evaluated separately from sequence retention through other predictions (Supplementary Methods 6).

In CAMPI fusion-entrapment controls [27], linked target/control pairs entered before selection and remained linked through parsimony. In donor-selected controls, the transferred database was fixed before FDRBench v0.0.4 [44] generated entrapments and decoys. Paired estimates weighted accepted controls according to acceptance and ranking of their paired targets; unresolved fusion identifications contributed an additional term. These are control diagnostics rather than calibrated FDR estimates for ordinary searches. Supplementary Methods 6 specifies denominators and tie handling. Restricted-reference simulations used AlphaPeptDeep [45] spectra from known peptides and retained all baseline and stressed-mixture outcomes.

The PREDICT spectrum comparison reused same-spectrum public-sequence and specimen-MG databases with 5–50-residue peptides, reporting up to five candidates and selecting the highest-hyperscore target. FastaLake-only spectra passed its peptide threshold without an accepted best MG target. Literal I/L-distinct sequences were checked against the MG FASTA and an in silico digest. Supplementary Methods 6 defines the categories and their interpretation.

### Transfer, pooling, and common study search spaces

Healthy-stool acquisitions were searched against databases selected from other acquisitions. For own and transferred accepted sets A and B, count retention was |B|/|A| and identity recovery |A∩B|/|A|. Other-11 pools united the remaining technical-replicate FASTAs, excluding the target FASTA but retaining shared upstream cohort evidence. Selection was not repeated on pooled predictions (Supplementary Methods 7).

The spectrum-split sensitivity searched identical odd-scan MS/MS spectra against databases selected from all, odd or even predictions, retaining all precursor (MS1) spectra. Clinical arms used public reservoirs and matched MG comparators. In 10-fold cross-fitting, deterministic native-scan folds withheld each scored spectrum from selection. For each fold, cohort evidence caches were rebuilt from the full reservoir using out-of-fold predictions from every acquisition of the original analysis roster (Supplementary Methods 7), and each acquisition’s database was selected from its own out-of-fold predictions. Held-out spectra were searched against that database, the all-prediction database and matched MG using identical settings. Fold PSMs were pooled per acquisition and database, rescored with a Sage-derived linear discriminant and assigned picked-peptide *q* values. Singular fits in three acquisitions used Sage’s heuristic fallback and remained in the analysis. Gains used acquisition medians with participant-level bootstrap intervals; four PREDICT acquisitions lacking patient IDs were separate resampling units. All 591 matched pairs were retained. Supplementary Methods 7 gives score-reproduction checks, conversion exceptions and sensitivity analyses.

Clinical primary analyses used acquisition-specific databases followed by study-level grouping. Separate searches against cohort-common FASTAs were a sensitivity analysis using the original 306 PREDICT and 331 CapScan acquisitions. The latter predates inclusion of the 16 participant-15 intestinal-fluid acquisitions; its 452,255-protein union differs from the 460,977-protein union of all 347 acquisitions. Supplementary Fig. 7b retains this comparison. Healthy stool compared acquisition-specific searches, separate common-FASTA searches and joint common-FASTA searching. Only joint searching combined confidence estimation and cross-run LFQ. Primary identification and quantitative endpoints required an accepted peptide in the same acquisition, excluding transferred-only signals (Supplementary Methods 7).

Joint searches used a Sage build correcting LFQ q-value mapping, with per-acquisition confidence as primary and study-level confidence as sensitivity. Completeness required accepted peptides and positive quantity in all 12 technical replicates. All-replicate CV used sample standard deviation for features complete in both arms. Pairwise group CV used population standard deviation over mean raw intensity; Pearson correlation used log_2_ intensity. Pairwise comparisons used shared quantified groups without imputation. Acquisition pairs were not independent biological replicates. Supplementary Methods 7 gives preparation-replicate results, alternative feature sets, native-inference comparisons and build provenance.

### Clinical, regional, and cross-omics analyses

For 42 PREDICT patients observed off and on antibiotics, repeated patient-state measurements were combined by mean composition fractions and median group counts. On-minus-off changes weighted patients equally without imposing temporal order. Mean changes used 50,000 patient-bootstrap resamples; two-sided tests used 200,000 sign assignments with plus-one correction and Benjamini–Hochberg adjustment across human share and group depth. Exposure included prophylaxis and treatment; rifaximin for hepatic encephalopathy was added in sensitivity analysis (Supplementary Methods 8).

Enterotype Dysbiosis Score (EDS) and membership fields followed Keller et al. [29], with higher EDS denoting weaker assignment to a reference enterotype. Fig. 4h displays 19 fixed exploratory features across the earliest matched proteomic–metabolomic observations of 85 patients; selection provenance and display procedures are detailed in Supplementary Methods 8. Three post hoc metabolite–KO pairs in Supplementary Fig. 10 were selected from a separate full biochemical screen for opposing EDS associations, inverse pair associations and biochemical interpretability. Two-sided Spearman tests used the full 1,042-pair family for Benjamini– Hochberg correction. Within-feature z scores were clipped at ±2; non-detections remained missing.

Cross-omics comparisons used 77 complete PREDICT patient observations and 13 CapScan participants across regions 1–4 and stool. DNA/RNA genes were matched by identical encoded protein sequence; unresolved transcripts per million (TPM) were excluded and recorded. Native and functional profiles retained all observed features, with functional profiles excluding human-assigned signal. Relative gene/protein abundances were square-root transformed (Hellinger transformation) [46]. Metabolites required ≥70% finite values and were log_10_-transformed, median-imputed and standardized. CapScan profiles were aggregated within participant–region from available specimens and residualized for region. Euclidean profile distances were compared by Spearman correlation, with patient-label or whole-participant-block permutation tests. Individual-function correlations required ≥35 co-detected patients. Benjamini–Hochberg families, prevalence and population sensitivities, and complete transformation rules are given in Supplementary Methods 8.

CapScan summaries averaged acquisitions within specimen and then participant–region. Fifteen participants supported fluid–stool contrasts, assessed by exact paired sign-flip tests with Benjamini–Hochberg correction over four human-share comparisons. Functional comparisons used all 73,316 microbial OGs in 75 participant– region profiles. Each participant’s four fluid regions were compared with their own stool and the other 14 stools by mean Hellinger distance. A two-sided test using 99,999 stool-to-participant permutations assessed the mean within-person advantage, without multiple-testing adjustment. Prevalence filters were sensitivities; participant/region variance partitions were descriptive. DNA profiles used 301 completed metagenome runs, allocating gene TPM equally across distinct eligible annotations before hierarchical averaging. Supplementary Methods 8 gives DNA eligibility and bootstrap details.

### Non-specific searches and histidine boundaries

Non-specific Sage searches of 347 CapScan and 306 PREDICT acquisitions used acquisition-specific FASTAs and standard mass, modification and tolerance settings without quantification. Sage routes required rank-one targets at peptide *q*≤0.01; internal-boundary and complementary-product displays additionally required spectrum *q*≤0.01. De novo predictions used the top-30% filter. Non-K/R C termini ended in neither lysine nor arginine; tryptic searches permitted these only at protein termini. Broad fractions collapsed I/L, whereas residue-composition summaries retained I/L-distinct strings (Supplementary Methods 9).

Origins were assigned from all route-compatible parents in the acquisition FASTA as human, microbial, mixed or unknown; sequence-hash accessions without origin annotation remained unknown. Human/microbial panels excluded mixed/unknown peptides but retained them in Source Data. Summaries averaged within specimen and participant–region; 15 participants supported paired fluid–stool comparisons.

Internal boundaries excluded protein termini and required one compatible protein-sequence/start/end location. Histidine enrichment was the observed histidine indicator minus its expected fraction among eligible internal non-K/R positions of the same source protein, conditional on route-specific length, mass and cleavage constraints. Eligible I/L variants were equally weighted. Participant–region or patient means entered two-sided sign-flip tests and participant/patient bootstrap intervals, with Benjamini–Hochberg adjustment within complete enrichment families (Supplementary Methods 9 and 10).

A complementary product required a distinct peptide and spectrum on the opposite side of the same localized boundary in the same acquisition and route. Histidine and non-histidine boundaries were matched within protein for theoretical peptide availability and spectrum-depth stratum. Intestinal-fluid analyses used de novo, semi-tryptic and non-specific routes; PREDICT used de novo and non-specific routes. Shared spectra and sequence spaces precluded treating routes as independent replications. Supplementary Methods 9 and 10 give complete contrasts and uncertainty calculations.

### Statistical analysis

Patients or participants were the biological units unless stated otherwise. Resampling preserved pairing and repeated regions; spectra, peptides, acquisition pairs and distance entries were not independent biological replicates. Tests were two-sided where applicable, using NumPy, SciPy, pandas and statsmodels in Python; versions are specified in Supplementary Methods 11. Exact sign-flip tests enumerated sign spaces, paired two-endpoint comparisons used two-sided exact Wilcoxon signed-rank tests, Monte Carlo tests used plus-one corrections, and biological bootstrap intervals resampled whole participants. Peptide *q* values describe search confidence; Benjamini–Hochberg [47] *q* values adjust statistical *P* values. Exploratory families used *q*<0.05 and retained non-significant outcomes. Supplementary Methods 6–10 and Source Data define estimands, seeds and complete families.

### Computational resources and provenance

Computational timings excluded queueing and file conversion. Supplementary Fig. 15 and Supplementary Methods 11 report stage-level timings, hardware, software identities and sequencing-cost examples [30–33]. Chunked lookup against the 233-million-protein public reservoir was evaluated within an 11-GiB allocation; complete workflow measurements and input rosters are retained in Source Data. Reporting follows metaproteomics guidance [48].

## Data availability

The healthy-stool mass spectrometry data will be deposited to the ProteomeXchange Consortium via the PRIDE partner repository and made available upon publication. Because they derive from patients, the PREDICT mass spectrometry data, clinical covariates and metabolite data are held in the PREDICT consortium data hub and are available under controlled access through the consortium: requests should be addressed to J.T., who initiates the required data transfer agreement. For five PREDICT acquisitions (MicrobPred_13, 44, 90, 115 and 181), both validated reconversions used in the reported analyses are deposited (ThermoRawFileParser 2.0.0.0 for searches and 1.3.4 for the spectrum-split and cross-fitting analyses). The CAMPI and CapScan mass spectrometry data reanalyzed here are available under PXD023217 and PXD038906, respectively. CAMPI assemblies and metagenomic and metatranscriptomic reads are available from the European Nucleotide Archive under PRJEB42466. CapScan 16S rRNA and shotgun metagenomic reads are available from NCBI under BioProject PRJNA822660, and CapScan metabolomics data from the Metabolomics Workbench under ST002073, ST002075, ST002407, ST002409 and ST002411. PREDICT metagenomic reads are available from the European Nucleotide Archive under PRJEB89898; the remaining metagenomes and the metatranscriptomes are being added to the same project. AlphaNovo prediction exports for all 726 acquisitions and the selected search FASTA files will be made available upon publication. Source data are provided with this paper as FastaLake_Source_Data.zip. Its README.md and SOURCE_DATA_MAP.tsv identify the file, sheet and columns underlying each figure panel, table and numerical statement, and it includes complete test families, non-significant results and sensitivity analyses. PREDICT participant and specimen codes are replaced by random pseudonyms used consistently across all files.

## Code availability

FastaLake is available under the MIT license at https://github.com/MannLabs/fasta-lake. The repository’s Dockerfile builds a container image (ghcr.io/mannlabs/fasta-lake) with the compiled Rust binaries and the official Sage v0.14.7 release executable, so no local Rust or Sage installation is needed. Paths and function names cited in Methods and Supplementary Methods refer to version 1.1.0rc2 (commit 9ad09c9; doi:10.5281/zenodo.23215189), in which the lookup, selection-parsimony and study-grouping functions used here are unchanged apart from added input checks. The analysis and figure code for this study, including study-level protein grouping, functional and origin profiles, and clinical, cross-omics and regional analyses, is available under the MIT license at https://github.com/treitpeter/fastalake-paper. A versioned release will be archived on Zenodo upon publication. Its pipelines folder holds the code of each upstream analysis, its figures, tables and text_statistics folders regenerate every panel, table and number from Source Data, and docs/REVIEWER_GUIDE.md lists every number in the manuscript with the function and Source Data table that produce it. For the Fig. 4h display, Source Data retain the selected-feature list, and the display code reproduces its values. All searches except the joint healthy-stool searches used the unmodified Sage v0.14.7 release (commit 99407db, whose executable reports version 0.14.6). The joint healthy-stool searches used a build of the same source that keeps target and decoy *q* values separate when pooled LFQ *q* values are assigned to peaks. This defect was fixed upstream after the v0.14.7 release in lazear/sage commit 0e2c7a6, and the source, patch, lockfile and checksums of the corrected build are included in the analysis release.

## Acknowledgements

This work was supported by the Max Planck Society for the Advancement of Science and by the European Union’s Horizon 2020 research and innovation programme under grant agreement No 825694 (MICROB-PREDICT). We thank our colleagues at the Department of Proteomics and Signal Transduction for their support and fruitful discussions, especially Igor Paron for his assistance with mass spectrometry measurements and data acquisition. We thank the PREDICT patients, the CapScan participants and the healthy-stool volunteers for their samples, the MICROB-PREDICT consortium for the PREDICT samples and clinical data, and the CAMPI and CapScan teams for making their data available. We thank the computing cluster of the Max Planck Institute of Biochemistry and the Max Planck Computing and Data Facility (MPCDF) for computational resources, and the institute’s mass spectrometry and IT infrastructure staff for their support.

## Author contributions

P.V.T. conceived the study, developed the methodology and software, performed the measurements and analyses, curated the data, made the figures and wrote the manuscript. J.B.M.-R. and M.M. conceived and supervised the study, and M.M. acquired funding. F.A.R. contributed to the conceptualization and supervised parts of the work. P.H. and M.L. developed AlphaNovo, W.-F.Z. contributed de novo and deep-learning methodology, and K.B. supervised the AlphaNovo work. P.E.G. designed the healthy-stool study and collected and measured the samples. J.X. contributed bioinformatic analyses. V.B., K.K., E.H.R. and M.T.S. contributed software and computing infrastructure. J.X., M.Z., A.S. and G.W. contributed to the bioinformatic conceptualization in discussions. M.I.K. and M.K. provided the PREDICT metagenomes and metatranscriptomes, J.T. provided the PREDICT cohort and clinical data, and C.J. provided the PREDICT metabolomics. M.A. contributed metagenomic and functional-annotation expertise and PREDICT data, and interpreted results. D.S., K.C.H. and J.A.G. provided the CapScan cohort and data, and J.A.G. contributed to biological interpretation. All authors reviewed and edited the manuscript.

## Supplementary Information

### Supplementary Figures

**Supplementary Figure 1.**
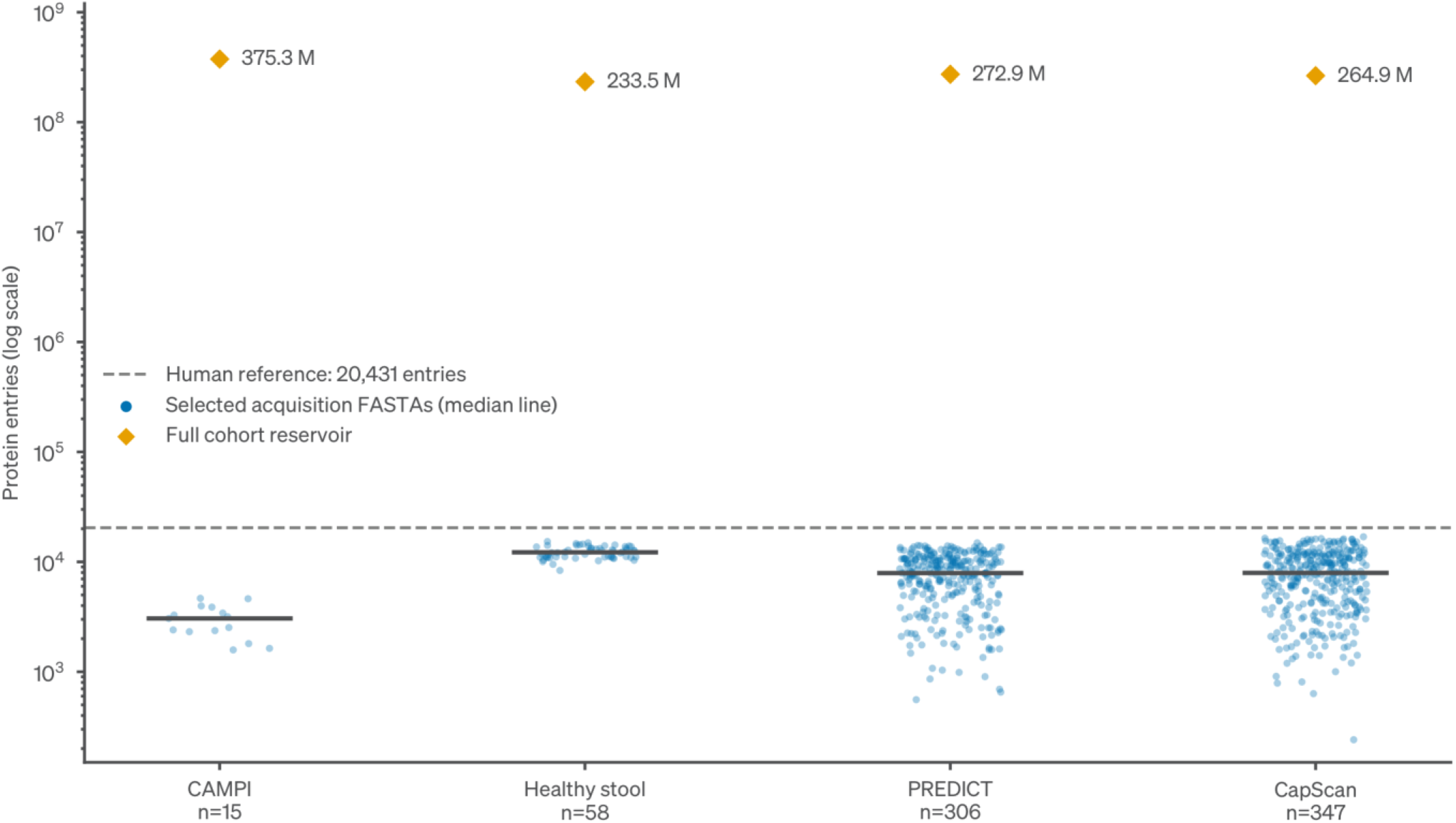
Selected database sizes in the context of the full sequence reservoirs. Blue points show the entry counts of all 726 acquisition-specific FASTAs. Horizontal bars show cohort medians. Orange diamonds show the full reservoirs, and the dashed line marks the 20,431-entry human reference proteome. The entry-count axis is logarithmic. PREDICT and CapScan reservoir diamonds describe assembly-inclusive analyses. Their public-sequence benchmarks used the separate 233,497,702-entry reservoir. Cohorts comprise 15 CAMPI, 58 healthy-stool, 306 PREDICT and 347 CapScan acquisitions.

**Supplementary Figure 2.**
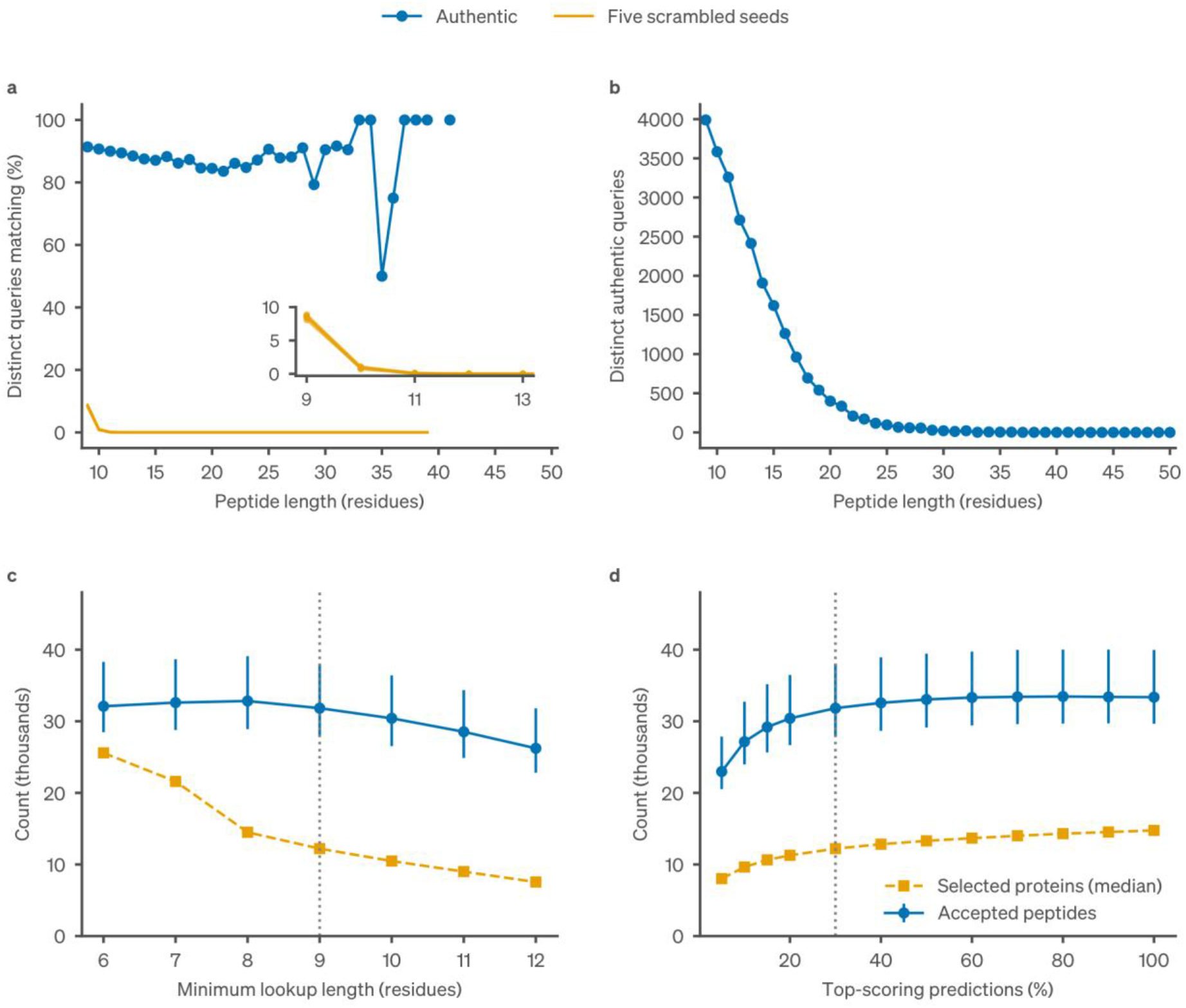
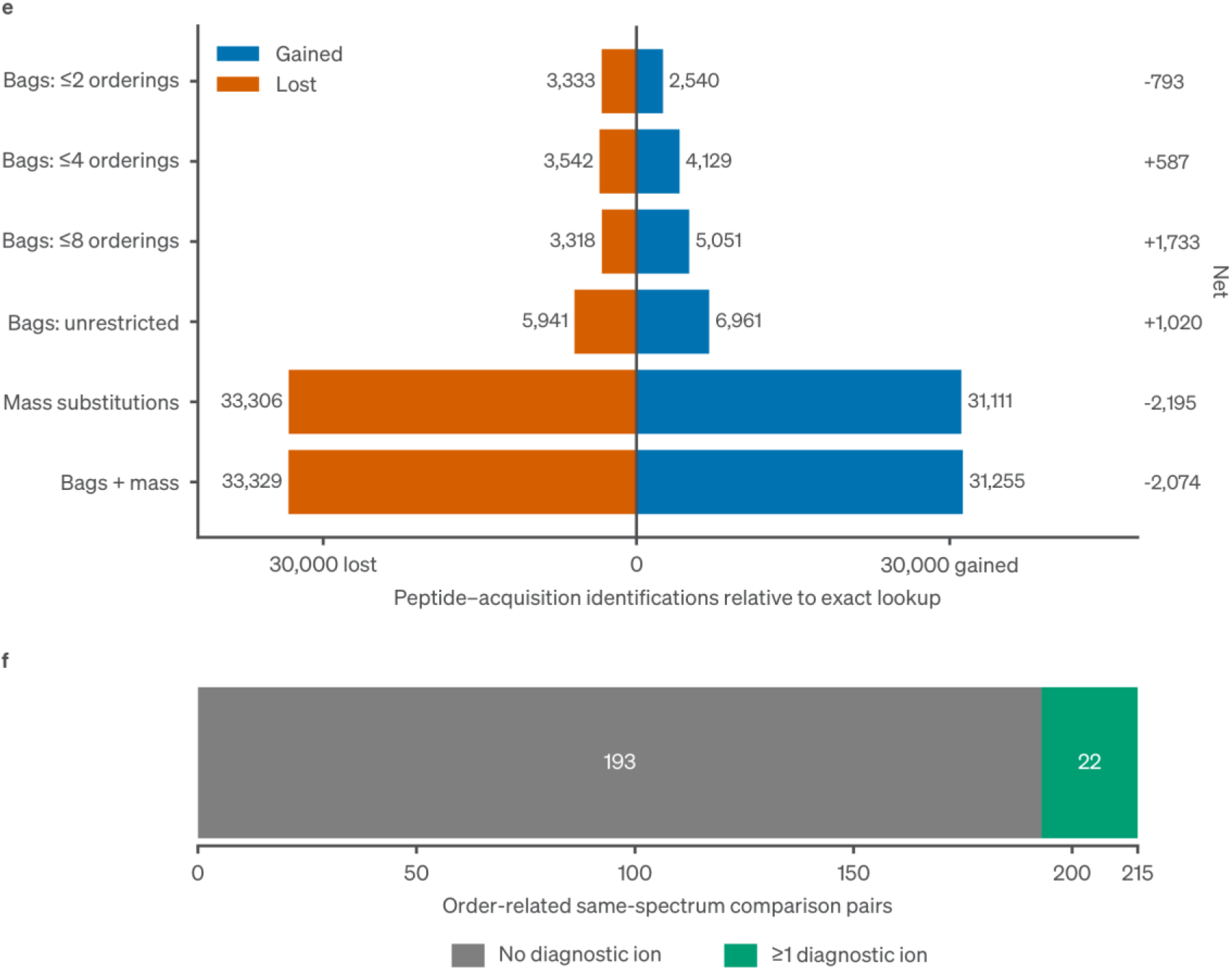
Specificity and sensitivity of peptide-guided database selection. **a,b**, Length-specific lookup fractions and authentic-query counts in the representative healthy-stool acquisition. Orange traces are five composition-preserving scrambled sets, enlarged for lengths 9–13 in the inset. Denominators are distinct queries. **c,d**, Sensitivity of selection to its settings, with 58-acquisition upstream pools and evaluated rosters. The nine-residue point in c reuses the top-30% run in d. Points show median accepted peptide counts with exclusive-quartile ranges and median selected protein counts. Counts strip modifications and keep isoleucine and leucine distinct. Vertical lines mark production settings. Source Data contain all 1,102 arm–acquisition records. Panels e,f, shown in the continuation of this figure, report the separate exploratory bag/mass comparison and fragment audit. Experimental retrieval using residue-order bags and mass substitutions. **e**, Gained (blue) and lost (vermilion) peptide–acquisition identifications relative to exact lookup in the 48-acquisition panel. The right column gives the net difference. Counts of 5–50-residue peptides are summed after within-acquisition peptide deduplication at nominal peptide *q*≤0.01. The ≤2, ≤4 and ≤8 bounds apply to the number of orderings of an eligible query. Unrestricted bags retain the recorded order uncertainty. **f**, Separate six-acquisition fragment audit of gained composition-preserving rearrangement pairs for a diagnostic annotated ion. Diagnostic support distinguishes the accepted sequence from the exact-arm comparator under the stated fragment model. Protocols and all per-acquisition values are provided in Supplementary Methods 2 and Source Data.

**Supplementary Figure 3.**
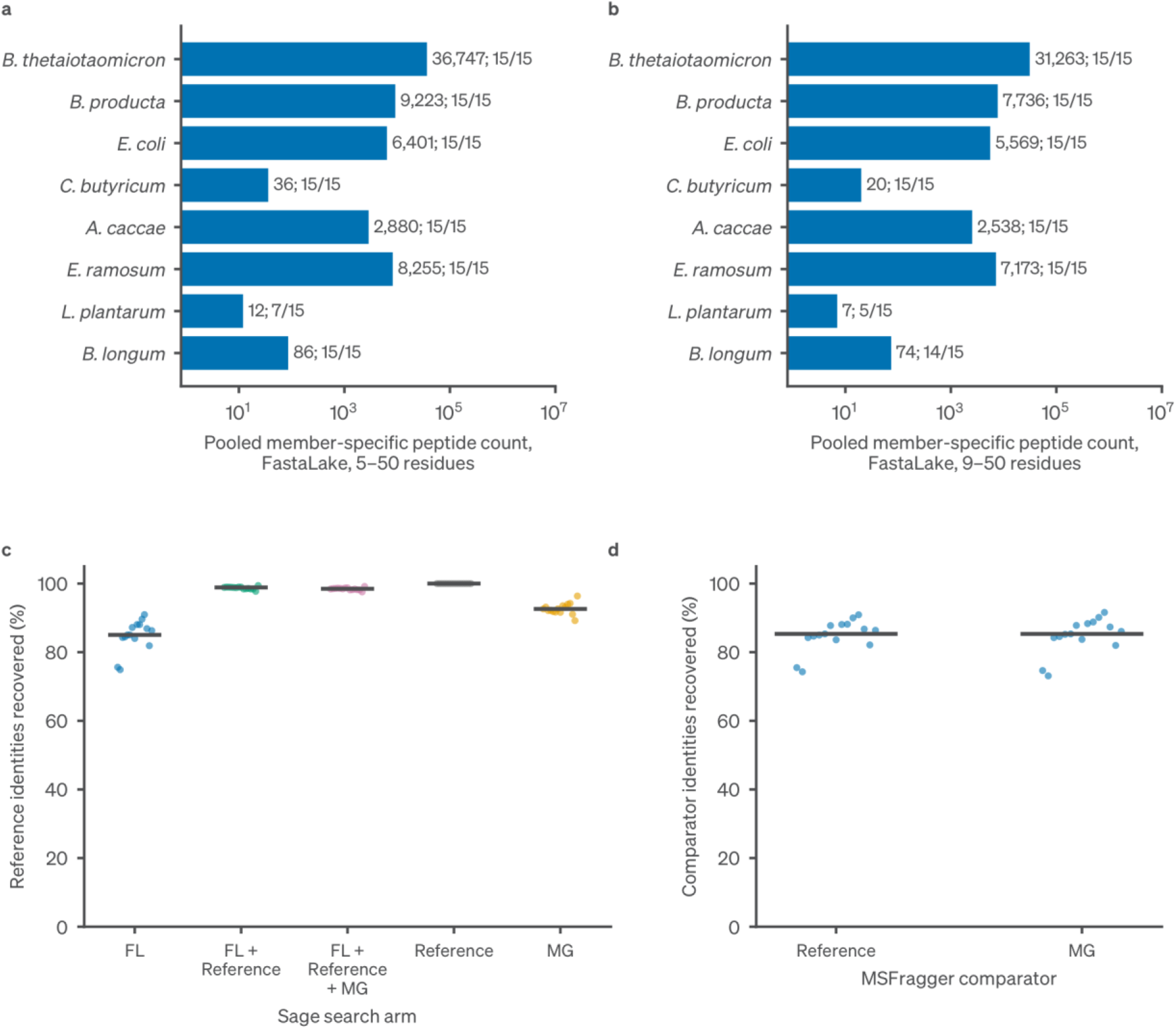
CAMPI member support and identity recovery. **a,b**, Pooled member-specific peptide counts in the FastaLake arm, with the number of acquisitions contributing at least one such peptide (labels give the count and acquisitions/15). Specificity is relative to the eight supplied reference proteomes. **c**, Within-acquisition recovery of reference identities by each Sage search arm. FL + Reference and FL + Reference + MG append the reference proteomes alone or together with the multi-omic co-assembly proteins (MG) to the FastaLake (FL) database after selection. **d**, Within-MSFragger recovery of reference and MG identities by FastaLake. Dots are acquisitions and bars are medians. All five arms, lengths, zeros and acquisition-level member counts are in Source Data.

**Supplementary Figure 4.**
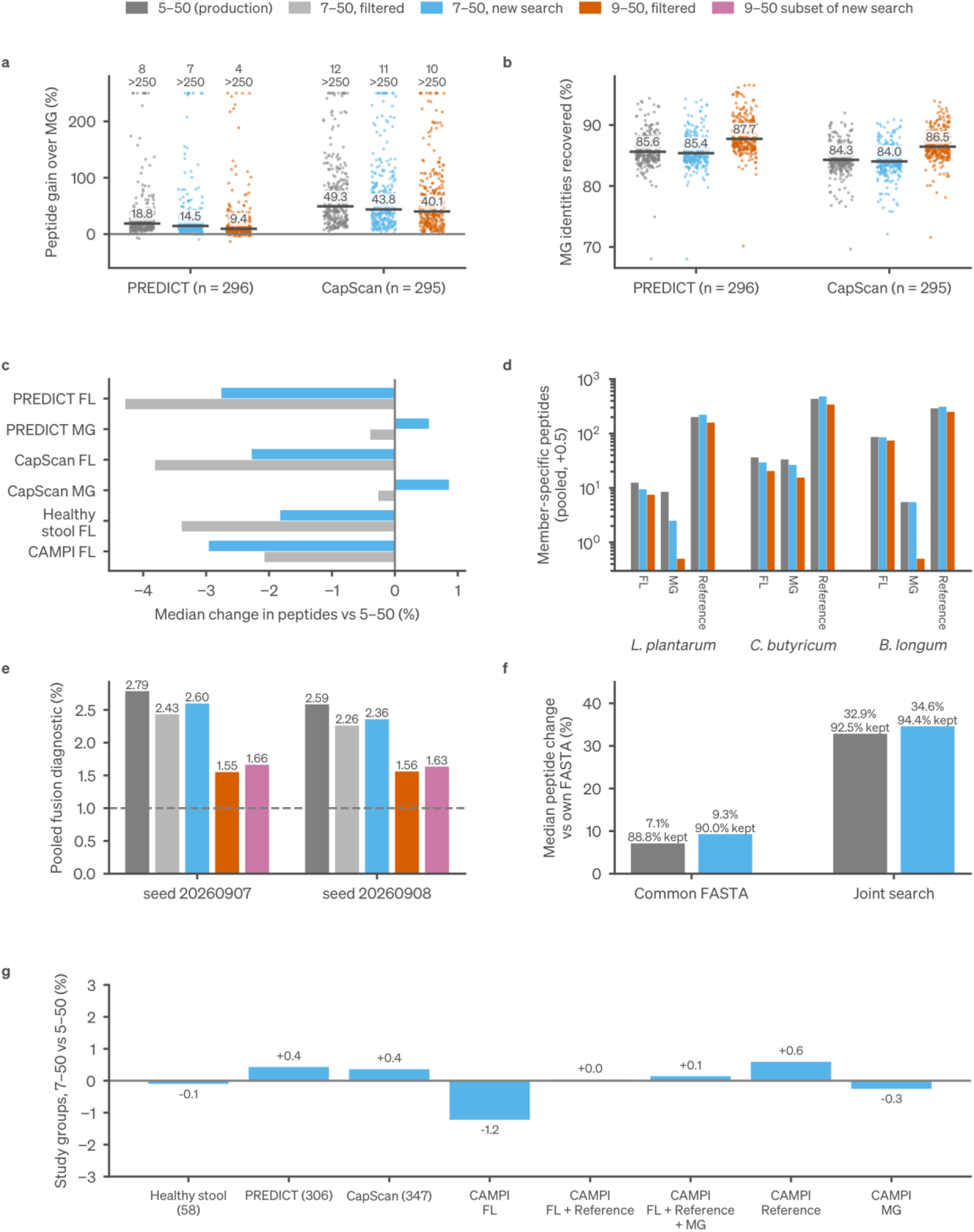
Peptide-length sensitivity of FastaLake results. All 2,035 database searches (2,030 standard plus five repaired PREDICT searches with label-free quantification (LFQ) enabled) were repeated with a minimum peptide length of seven residues and otherwise unchanged settings. Confidence was re-estimated in each new search. Gray, production 5–50-residue searches. Light gray, the production results filtered to ≥7 residues (original *q* values). Blue, new ≥7-residue searches. Orange, production results filtered to ≥9 residues. Purple, the ≥9-residue subset of the new searches. **a**, Per-acquisition peptide gain of public-sequence FastaLake over specimen-matched metagenome (MG) databases in 296 PREDICT and 295 CapScan acquisitions, with FastaLake databases selected from the scored spectra. Bars and labels, median paired gains. Values above 250% are drawn at 250% and counted. **b**, Recovery of MG-search peptide identities. **c**, Median change in accepted peptides relative to 5–50 residues after filtering to ≥7 residues or re-searching at ≥7, by database. **d**, Pooled member-specific peptides for the three least represented CAMPI members in FastaLake (FL), MG and reference searches, with 0.5 added for the logarithmic axis. **e**, CAMPI fusion-control diagnostic (paired estimate plus unpaired worst case), pooled over 15 acquisitions per control seed at peptide *q*≤0.01. Dashed line, 1%. **f**, Healthy-stool searches against the common FASTA or in the joint search with study-level confidence, relative to each acquisition’s own database (58 acquisitions). Labels give median peptide change and median fraction of own-database identities retained. **g**, Change in study-level protein groups under direct peptide assignment.

**Supplementary Figure 5.**
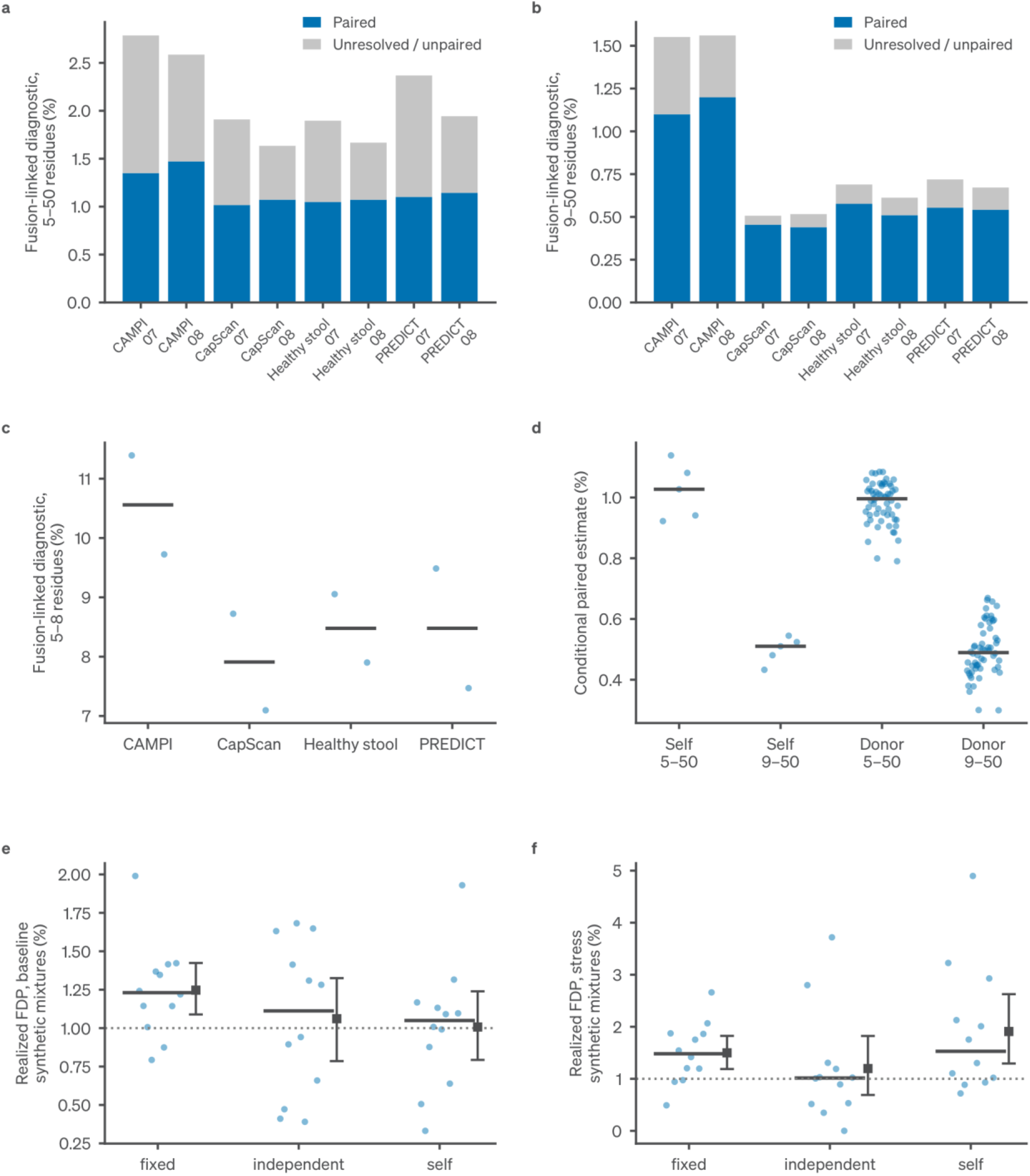
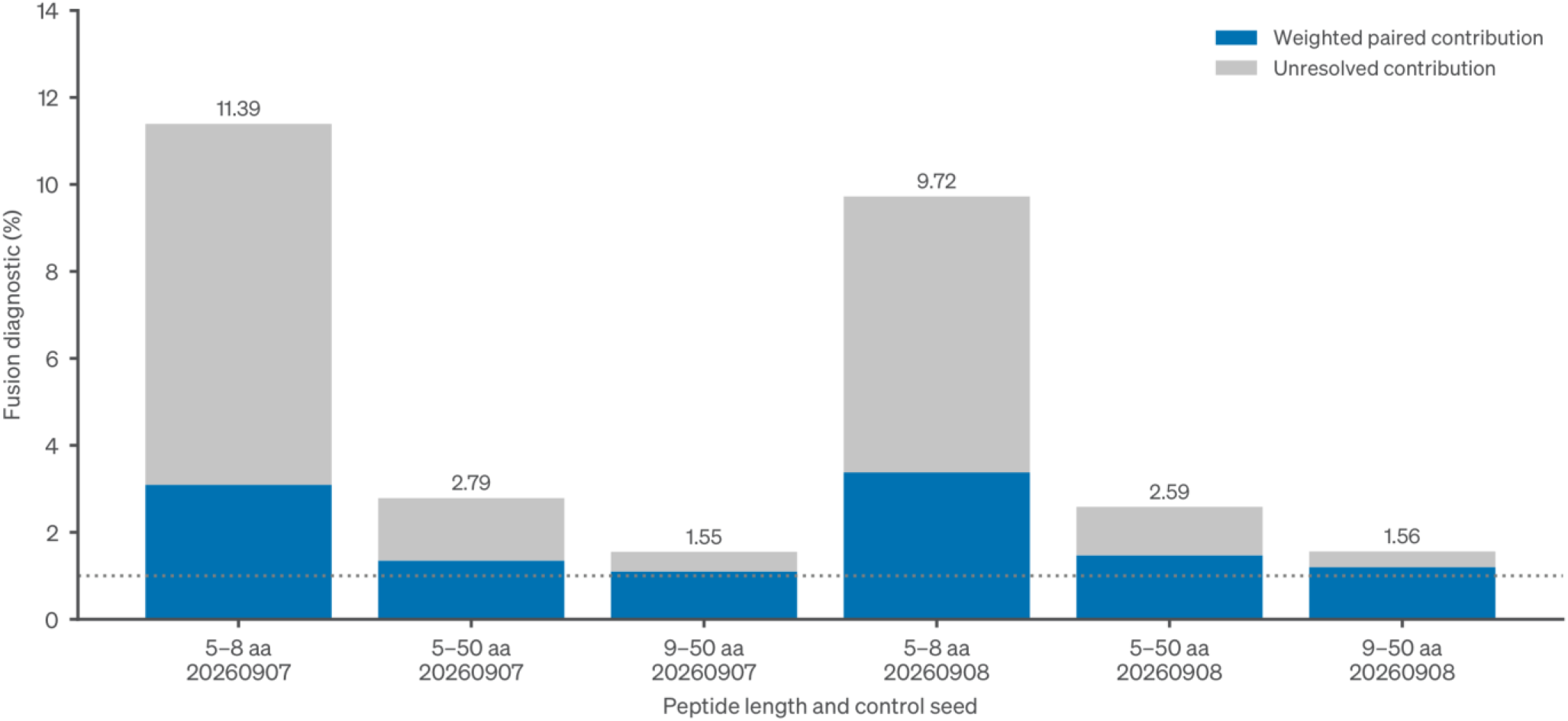
Error controls at different selection stages. **a,b**, Fusion-linked diagnostics at nominal peptide *q*≤0.01, including both paired and unresolved contributions. Each bar is one seed. The CAMPI data use the broad UniProt reservoir. **c**, The nested 5–8-residue population retains the same search *q* values. **d**, Explicit-peptide self-selected (five) and donor-selected (58) acquisition controls, each averaged over three seeds. **e,f**, Known-truth baseline and stress simulations (12 independent mixtures per regime). Dots are mixtures and horizontal bars are medians. Squares and intervals are means and 95% whole-mixture bootstrap intervals. The dotted line marks nominal 1%. Decomposition of the CAMPI fusion diagnostic. Fifteen acquisitions pooled separately for each seed from the 375,333,654-sequence reservoir, peptide *q*≤0.01. Blue, weighted paired contribution (E + E₀ + 2B)/N, with terms defined in Supplementary Methods 6. Gray, unresolved contribution U/N. Labels give the total percentage. Dotted line, nominal 1%. Length subsets retain the original *q* values. The other cohorts, searched with smaller reservoirs, are shown in panels a–c.

**Supplementary Figure 6.**
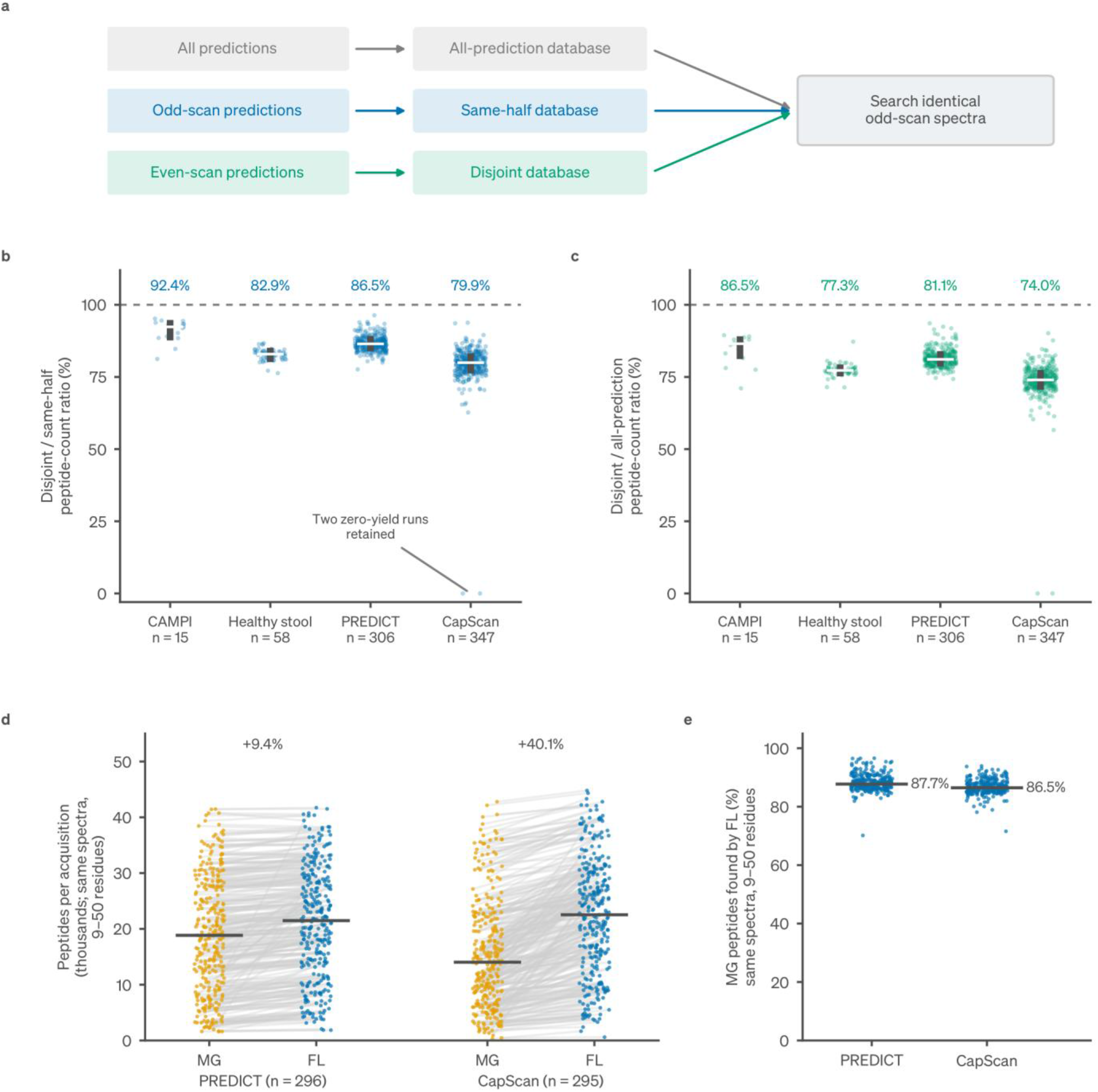
Yield with shared and disjoint spectrum-selection inputs. **a**, Protein databases were selected using all predictions (all-prediction selection), odd-scan predictions (same-half selection) or even-scan predictions (disjoint selection). All three searches scored identical odd-scan MS/MS (MS2) spectra. **b,c**, Disjoint peptide-count yield relative to same-half selection (b) or selection from all predictions (c). Points are acquisitions and bars show medians and interquartile ranges. Dashed lines in b,c mark equal yield. All 726 acquisitions (15 CAMPI, 58 healthy stool, 306 PREDICT and 347 CapScan) are retained, including two CapScan disjoint zeros. **d,e**, Peptides per acquisition (d) and fraction of metagenome-search peptide identities recovered by FastaLake in the same acquisition (e) in matched metagenome (MG) and public-sequence FastaLake (FL) searches, with FastaLake databases selected from the scored spectra (296 PREDICT and 295 CapScan acquisitions). Lines pair acquisitions, and bars show medians. Numbers above the points in d are median paired gains in peptides per acquisition (FL relative to MG), as reported in the text. Cross-fitted values are shown in Fig. 4b,c. Gains with selection from all or even-scan predictions on odd-scan spectra, and 5–50-residue values, are in Source Data. Clinical inputs use public-sequence reservoirs. Input imbalance, confidence fallbacks and five reconverted PREDICT inputs are detailed in Supplementary Methods 7.

**Supplementary Figure 7.**
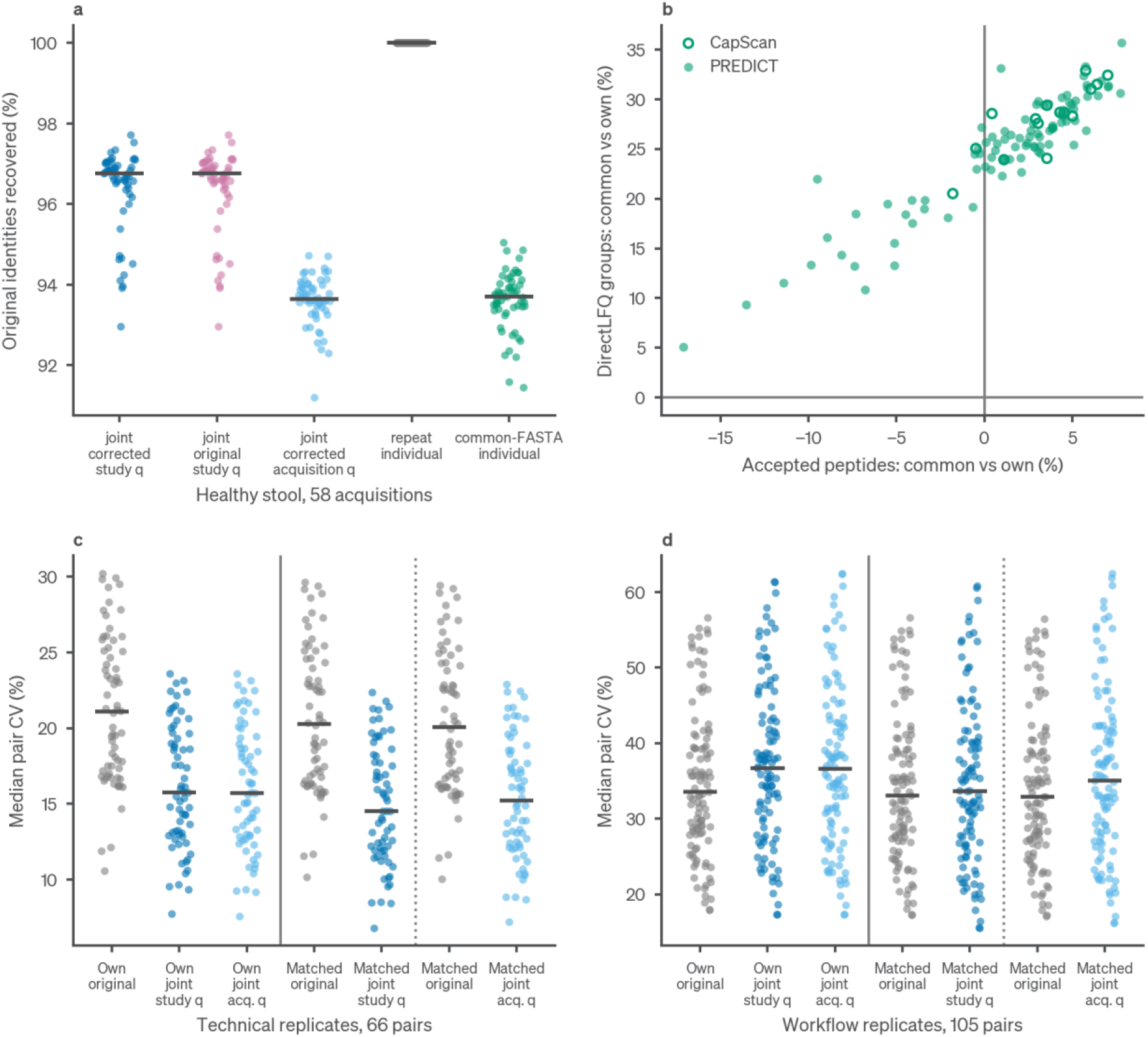
Common search databases: depth, identity retention and precision. **a**, Healthy-stool original peptide identities retained by the original and corrected joint searches with study-level confidence (study *q*), the corrected joint search with per-acquisition confidence (acquisition *q*), and repeat and common-FASTA individual searches. **b**, Clinical common-FASTA sensitivity analysis on the original 306 PREDICT and 331 CapScan acquisitions. The 16 participant-15 intestinal-fluid acquisitions added later are excluded; participant 15 contributes stool measurements. Participant-level changes in peptide counts and directLFQ protein groups for common-FASTA individual searches in PREDICT (93 patients) and CapScan (15 participants). Panels c,d use all accepted 5–50-residue peptides. **c,d**, Microbial protein-group precision under the original individual searches and the corrected joint search with study-level (study *q*) or per-acquisition (acquisition *q*) confidence: rows present in each arm separately (Own) versus rows matched across both arms (Matched), matched separately for each joint confidence definition. Each dot is an acquisition pair (66 technical or 105 preparation pairs). Pairs sharing an acquisition are dependent. Pair coefficient of variation (CV) uses population SD/mean. This differs from the separately supplied all-replicate, sample-SD CV. Bars are medians.

**Supplementary Figure 8.**
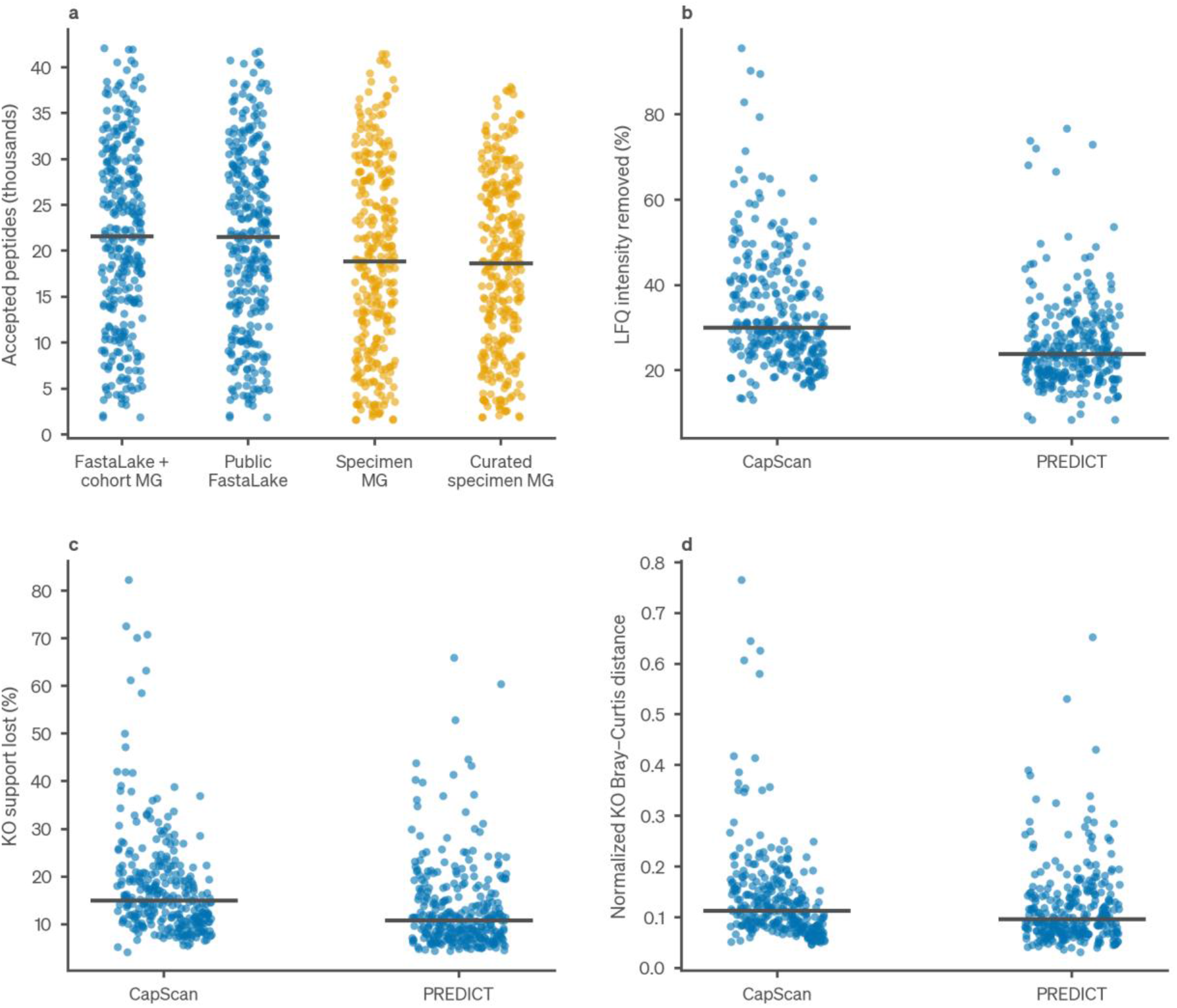
Clinical database comparators and evidence ablation. **a**, Accepted peptides in four PREDICT database arms on the same 296 acquisitions: full FastaLake including cohort metagenomes, public-sequence FastaLake, specimen-matched metagenome database (specimen MG) and curated specimen MG (FastaLake selection applied to the specimen metagenome). Database selection in panel a used predictions from the same spectra subsequently searched. The 5–50-residue results are provided in Source Data. **b–d**, Removing FastaLake-only peptide evidence from a fixed reporting dictionary: intensity removed, KEGG orthologs (KOs) losing all support, and Bray–Curtis distance between normalized all-evidence and shared-only KO profiles. Dots are acquisitions (296 PREDICT, 295 CapScan) and bars are medians. This ablation is conditional on the FastaLake dictionary and does not reconstruct a metagenome-only workflow.

**Supplementary Figure 9.**
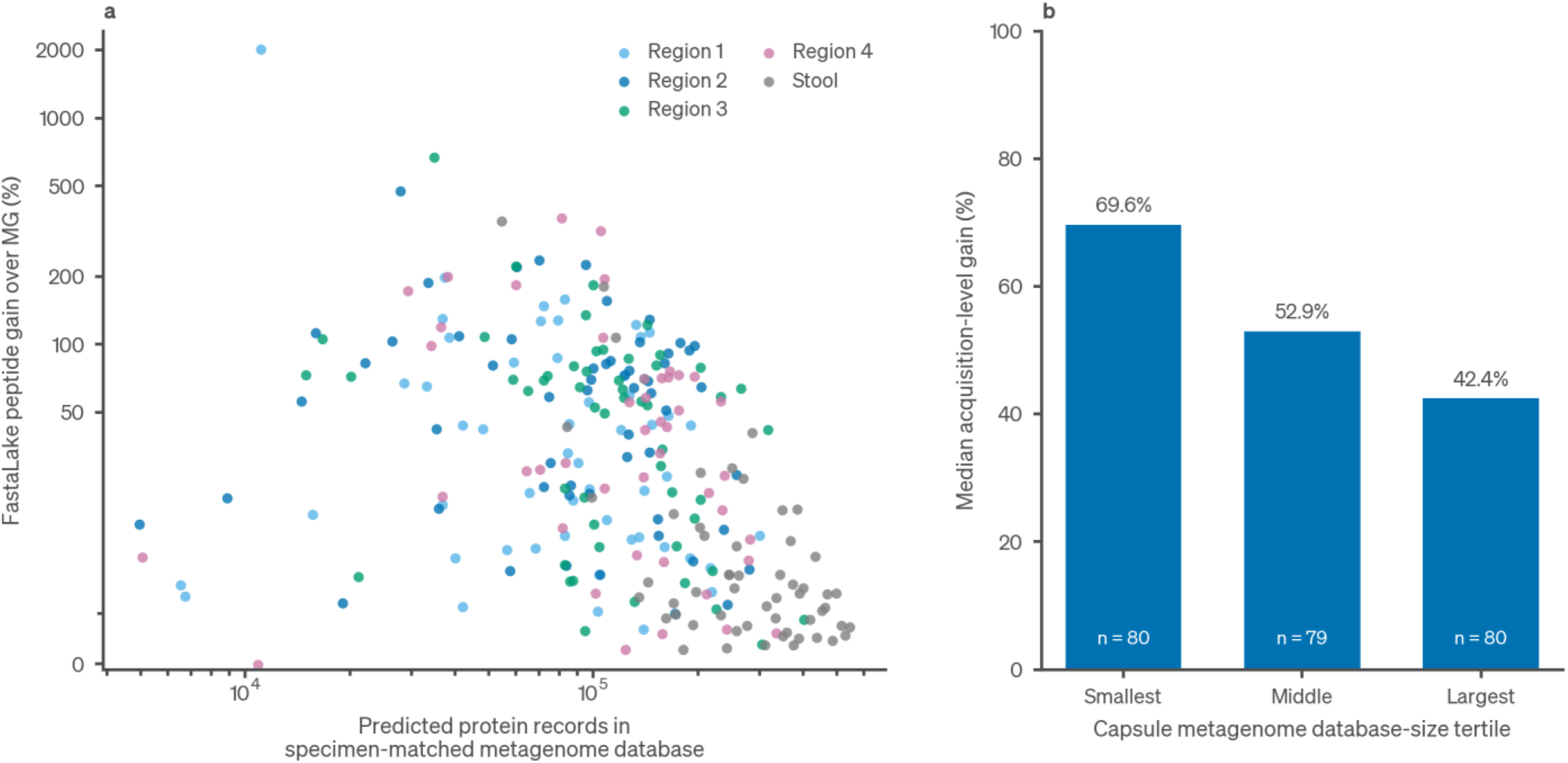
Predicted metagenome database size and FastaLake peptide gain. **a**, Each point is one of 263 matched specimens. Repeated acquisitions are summarized by their median gain and database size. Colors denote sampling region. Database size counts predicted-protein records retained in the specimen-matched metagenome FASTA after removing malformed records and duplicate accessions, before searching. Database size excludes added human reference proteins and is not a count of proteins identified by MS. The database-size axis is logarithmic. The gain axis is linear up to 50% and logarithmic above 50%. **b**, Median acquisition-level gain across increasing thirds of intestinal-fluid metagenome database size: 69.6%, 52.9% and 42.4% (80, 79 and 80 acquisitions). Specimens and participants may contribute repeated acquisitions. Gain is 100 × (FastaLake peptides / metagenome-search peptides − 1), with FastaLake databases selected from the scored spectra. Database size and gain also share the metagenome comparator.

**Supplementary Figure 10.**
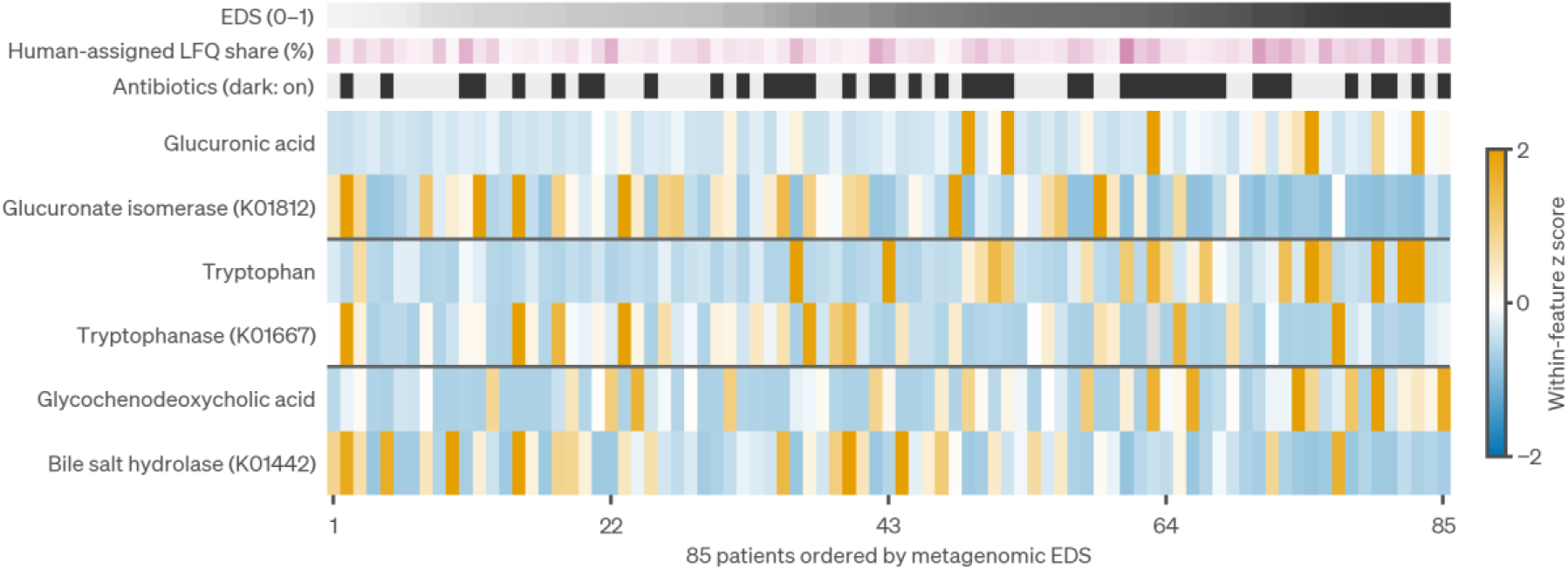
PREDICT metabolite–enzyme pairs ordered by Enterotype Dysbiosis Score (EDS). Three metabolite– enzyme pairs selected post hoc from the same-cohort screen, shown for 85 patients in EDS order. Positive values are standardized within feature and clipped at ±2. Gray indicates non-detection. Tracks above the heatmap show EDS (0–1), human-assigned LFQ share (%) and antibiotic exposure (dark, on). Supplementary Table 3 and the Source Data retain the original 20-feature, 259-observation selection, the focused measurements and complete association and sensitivity results.

**Supplementary Figure 11.**
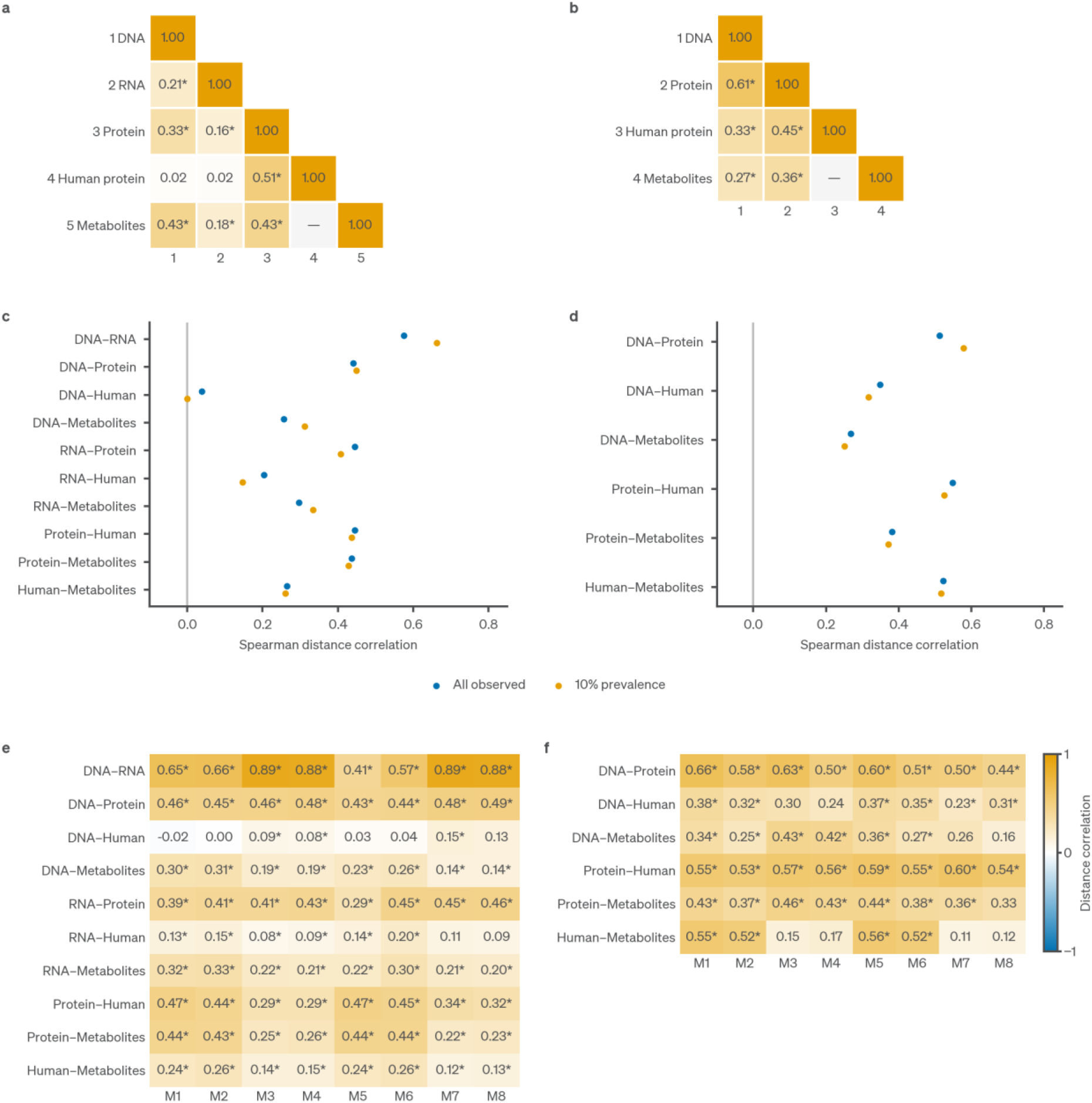
Pathway-level cross-omics profiles and sensitivity analyses. **a,b**, KEGG pathway distance correlations for 77 PREDICT patients (a) and the CapScan roster of 13 participants with complete DNA profiles in all five sampling groups (b). Native, eggNOG orthologous group (OG) and KO correlations are shown in Fig. 4i and Fig. 5g. All observed features are retained. Protein profiles exclude human-assigned signal and retain unresolved origin. **c,d**, Native-profile correlations with all observed features and with a ≥10% prevalence filter, for PREDICT (c) and CapScan (d). **e,f**, Controlled native-distance comparisons for PREDICT (e) and CapScan (f): M1–M4 use the ≥10% filter and M5–M8 all observed features. Each set shows Hellinger/Pearson, Hellinger/Spearman, rank/Pearson and rank/Spearman combinations. CapScan panels use region-adjusted profiles. Asterisks indicate *q*<0.05 within the complete comparison family after 1,999 design-preserving permutations. Source Data retain all native, OG, KO and pathway correlations, all 176 controlled estimates, including unadjusted CapScan comparisons, and all functional prevalence and population sensitivities.

**Supplementary Figure 12.**
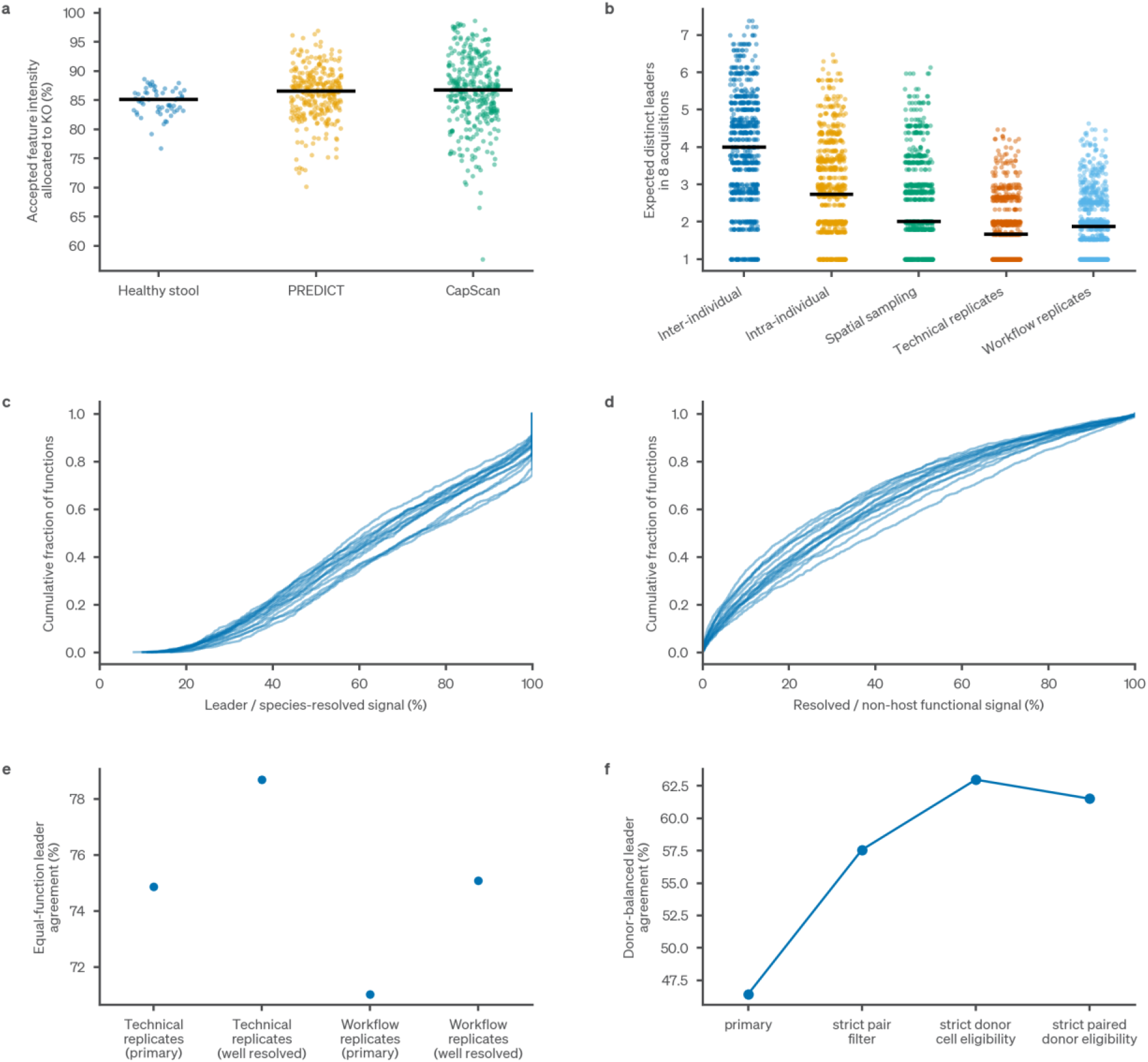
Functional allocation and leading-species attribution. **a**, Fraction of accepted quantified feature intensity allocated to a KEGG ortholog. The denominator includes the functionally unassigned remainder. Acquisitions are shown separately by cohort. **b, Expected leading-species diversity after rarefaction to eight resolved healthy-stool acquisitions, for the KOs eligible within each design (740–822 KOs). c,d**, Per-participant cumulative distributions across eligible CapScan functions distinguish leader share within species-resolved signal from species-resolved share of non-host functional signal (21,168 cells, 1,467 functions, 15 participants). **e,f**, Sensitivity of leading-species agreement to resolution rules in healthy stool (e) and eligibility rules in CapScan (f). All panels use direct peptide assignment. Species labels are representative labels retained only with a compatible same-KO carrier.

**Supplementary Figure 13.**
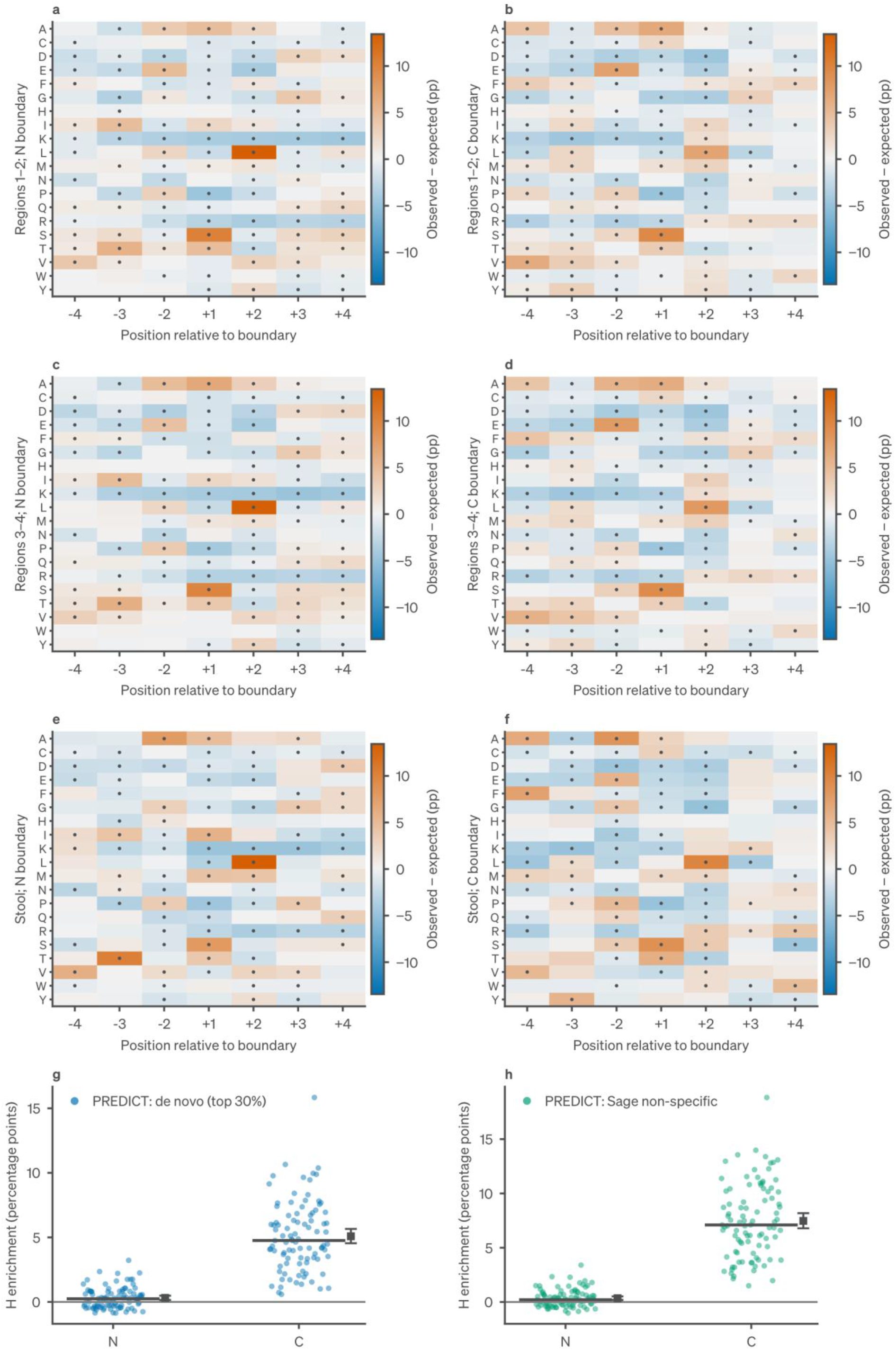
Histidine-boundary sequence context and PREDICT replication. **a–f**, CapScan histidine-conditioned motif deviations from source-protein opportunities for regions 1–2, regions 3–4 and stool, separately at N-terminal and C-terminal boundaries. Colors show participant-mean observed-minus-expected frequencies in percentage points. Dots mark *q*<0.05 in the full 32,400-test motif family, of which 840 cells are displayed. The conditioned position is omitted. **g,h**, PREDICT histidine enrichment for de novo and non-specific-search evidence, retaining both N-terminal and C-terminal boundaries. Dots are 93 patient means after averaging repeated acquisitions within specimens and specimens within patients. Bars are medians. Squares and intervals are means and 95% patient-bootstrap intervals.

**Supplementary Figure 14.**
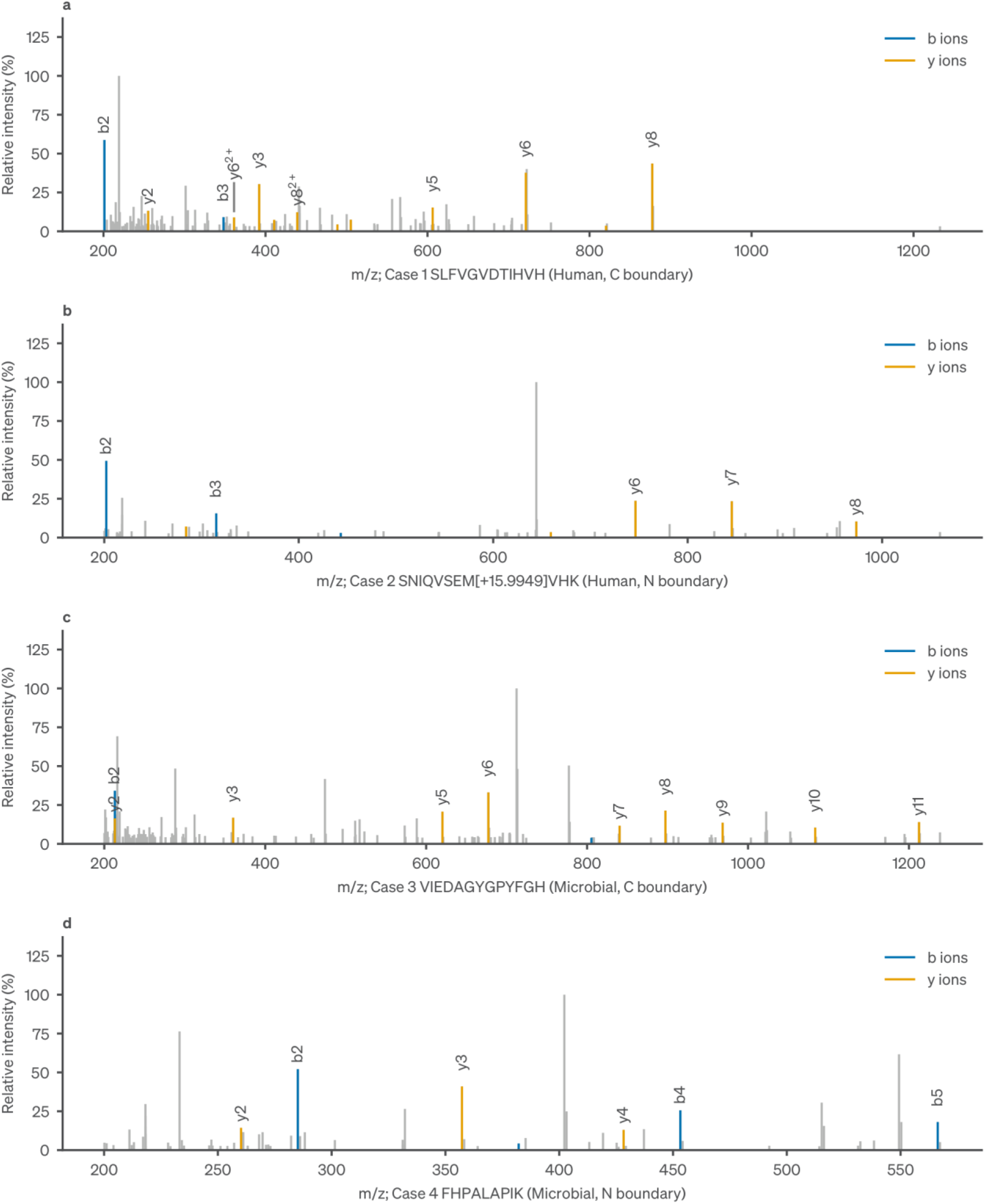
Measured spectra for selected PREDICT histidine-boundary peptides. Four selected PREDICT spectra from the non-specific search (cases 1–4 in a–d; Source Data case identifiers HIS_001–HIS_004). Gray peaks show the processed spectrum. Blue and orange peaks mark matched b and y ions. Intensities are normalized to each spectrum’s maximum processed peak. Thin gray leader lines connect displaced ion labels to their peak positions. Source Data include observed and processed peaks, all matched fragments, peptide–spectrum match (PSM) identifiers, source FASTA/RAW provenance and bounded sequence alternatives for every case. N-terminal-boundary histidine is inferred from the compatible source context, whereas C-terminal-boundary histidine belongs to the observed peptide. Selection is illustrative.

**Supplementary Figure 15.**
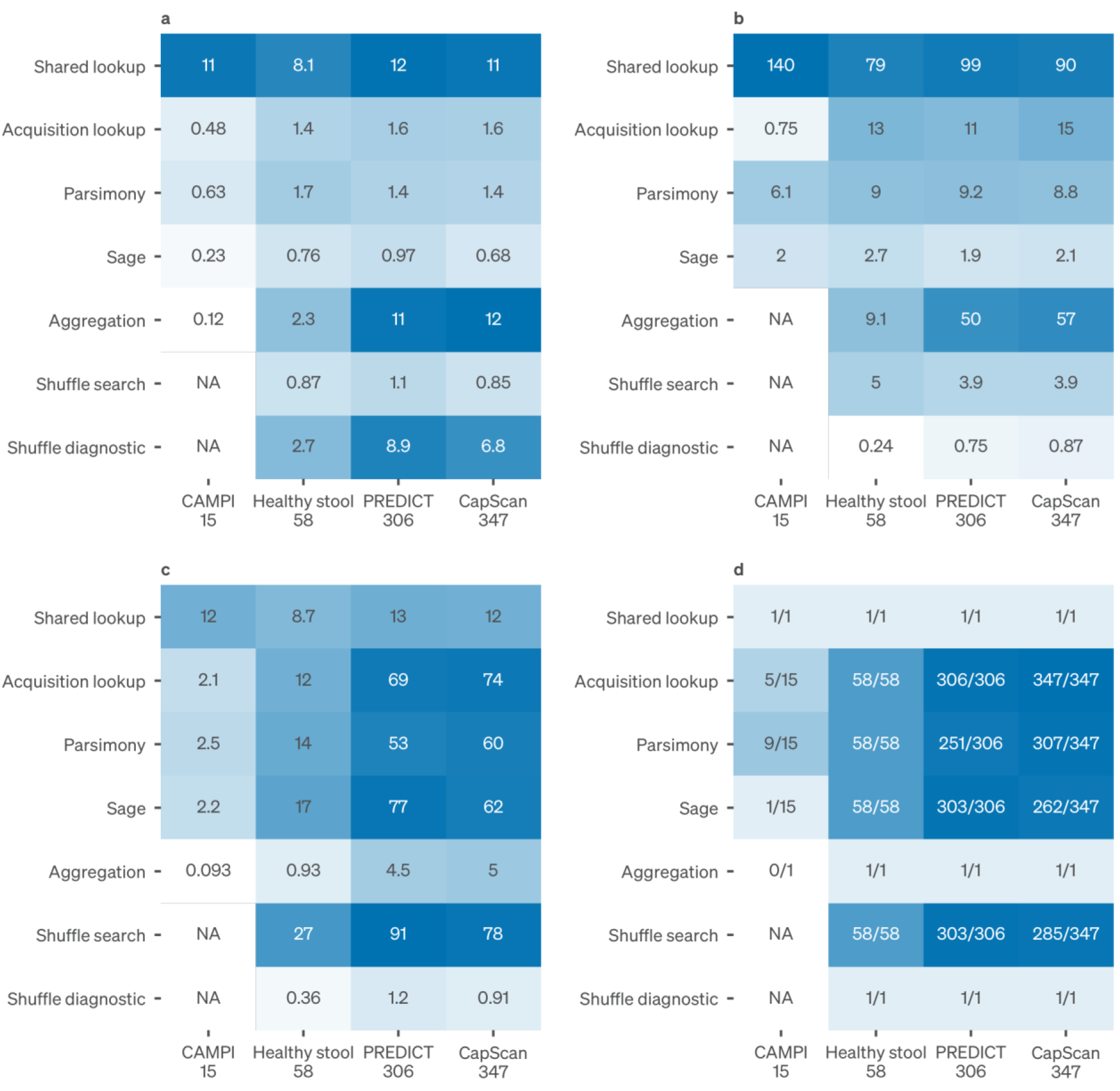
Measured computational resources. Execution and sampled-memory summaries for the CPU production workflow. Acquisition stages use 15 CAMPI, 58 healthy-stool, 306 PREDICT and 347 CapScan acquisitions. Shared healthy-stool and CapScan stages retain their actual 97- and 368-acquisition execution scopes. CAMPI timings come from the broad-reservoir production run. **a**, Median task execution minutes. **b**, Median usable sampled memory in GiB (2³⁰ bytes). **c**, Sum of allocated CPU-hours, distinct from consumed CPU time. **d**, Number of usable memory samples/number of tasks. Shading is log_10_(1+value). Printed values are untransformed. Queueing, conversion, de novo inference and downstream biological reanalysis are outside these stage timings. Available CAMPI inference records are supplied separately.

### Supplementary Tables

Supplementary tables are supplied as data files. Further numerical tables that support reported values are supplied as Source Data.

Supplementary Table 1 | Candidate-clustering search sensitivity at 99%, 97%, 95%, 90%, 75% and 50% sequence identity in 15 CAMPI acquisitions.

Supplementary Table 2 | PREDICT antibiotic endpoints under alternative handling of rifaximin for hepatic encephalopathy. Rows give the primary definition (42 patients), rifaximin for hepatic encephalopathy counted as exposure (40 patients), patients with any such record excluded (35 patients) and only off-state acquisitions with such a record excluded (38 patients). For human-assigned share and protein-group depth, the table gives patient and acquisition counts, state means, mean and median paired differences, 95% bootstrap intervals, sign-flip *P* values and *q* values.

Supplementary Table 3 | Enterotype Dysbiosis Score displays. Sheets All20_features_259_observations to 85_patient_z retain the earlier 20-feature, 259-observation selection and its 85-patient display (Fig. 4h). Sheets Focused_raw, Focused_features, Focused_pairs and Focused_adjustments give, for the three metabolite–enzyme pairs in Supplementary Fig. 10, the six displayed measurements and patient order, the feature associations, *q* values saved from the complete screen families, and all 192 descriptive adjustment estimates.

Supplementary Table 4 | Individual-function correlations between DNA, RNA and protein layers.

Supplementary Table 5 | Shuffled-query controls in the first 60 completed acquisitions and two seeds. At peptide *q*≤0.01, exact and bag routes yielded no accepted canonical peptide–acquisition–seed observations in these acquisitions. The mass route yielded 13,512 and the combined route 13,518. Selected protein entries summed across acquisition–seed runs were 55,229, 78,239, 561,126 and 561,331, respectively. In the extension to the full roster (645 of 726 acquisitions completed), exact and bag routes accepted peptides in one acquisition–seed only, where a shuffled 9-residue query selected human titin, an abundant protein in that sample; 185 of its 190 exact-route peptides were also accepted in the native search of that acquisition. Full acquisition and seed rows and the summary are in Source Data.

Supplementary Table 6 | Full-route shuffled-control acceptance and control exposure.

### Supplementary Data

Supplementary Data 1 | Sample_rosters.xlsx: analysis-specific acquisition rosters, recorded specimen and participant links, and exclusions for all 726 acquisitions (15 CAMPI, 58 healthy-stool, 306 PREDICT and 347 CapScan acquisitions).

## Supplementary Methods

These procedures describe the four study datasets, sequence selection and search, downstream quantification, and validation and sensitivity analyses. Numbered references refer to the Supplementary References at the end of this document.

### Supplementary Methods 1 | Study design, samples and experimental measurements

#### Study design and datasets

We analyzed four metaproteomic datasets: the CAMPI SIHUMIx benchmark [1], a healthy-stool variability series, the PREDICT liver-disease cohort [2], and the CapScan intestinal sampling study [3,4], all acquired by data-dependent acquisition (DDA).

Healthy-stool analyses used public-sequence FastaLake. PREDICT and CapScan biological analyses used assembly-inclusive FastaLake, in which proteins from all 296 and 335 available assembly FASTAs, respectively, were pooled with public-sequence resources in each cohort reservoir. The 335 CapScan assembly FASTAs represent 335 sequenced specimens. Of these, 301 had a completed public shotgun run that supplied the DNA profiles (one run lacking its second read file was excluded); 245 of these runs came from the 13 participants with complete DNA profiles in all five sampling groups and from specimens that also had protein and metabolite profiles, and the 295 matched-benchmark acquisitions (263 specimens) are the acquisitions of the 347-acquisition roster whose specimen had one of the 301 completed runs; three participant-15 intestinal-fluid acquisitions have no specimen metagenome. The matched-metagenome benchmark (Fig. 4b–d) instead used public-sequence FastaLake, comprising the Unified Human Gastrointestinal Genome (UHGG), the gut subsets of the Global Microbial Gene Catalog (GMGC) and the Global Microbial smORFs Catalog (GMSC) and human Swiss-Prot, without study-specific assembly FASTAs. PREDICT and CapScan specimen-matched metagenome comparators used the assembly of the corresponding specimen plus human Swiss-Prot. CAMPI used the broad-UniProt reservoir and post-selection reference additions described below.

We used 15 of the 21 CAMPI SIHUMIx acquisitions: S01–S10, the four S11 fractions and S12 [1]. Source records identified five laboratories. We distinguished gradients, fractions and technical measurements from biological replicates. The 58-acquisition healthy-stool series comprised 12 technical replicates of one preparation, 15 independent preparations of one stool sample, 10 spatially separated aliquots from one stool sample, 11 longitudinal samples from one donor, and 10 inter-individual measurements from different donors. The technical-replicate, preparation and spatial experiments used material from the longitudinal donor. We excluded from this variability series the 39 differential-centrifugation acquisitions that contributed to the original upstream sequence selection. The supplied protocol links the spatial aliquots to longitudinal sample 2. This donor is also one of the 10 inter-individual donors. PREDICT comprised 306 acquisitions, of which 302 were linked to 93 patient identifiers. Analysis-specific populations included 296 matched search comparisons, 91 visit-1 records, 42 patients observed in both antibiotic states, 85 earliest matched proteomic–metabolomic visits and 77 earliest visits with complete molecular measurements. The CapScan study [3,4] contributed 347 acquisitions from 308 specimens and 15 participants, counted from the deposited files.

The 16 participant-15 intestinal-fluid acquisitions were linked to their specimens through the source-study workbook, because the sample overview listed supernatant rather than pellet identifiers for this participant. Region assignments were retained from the source study. Sixteen acquisitions, 14 of them in this roster, carry different specimen numbers in file names and instrument records. Metadata joins, preparation relationships and analysis populations are provided in Source Data and Supplementary Data 1 (Sample_rosters.xlsx). (analysis: fastalake-paper, pipelines/sample_rosters_workbook and capscan_participant_roster)

#### Experimental procedures and mass spectrometry

##### Healthy-stool reproducibility series

Healthy-stool and PREDICT samples (50 mg stool) were lysed without prior pelleting in 200 µl PreOmics LYSE buffer by two cycles of 10 min at 100 °C followed by 20 min at −80 °C and by Bioruptor sonication (30 cycles of 30 s), and nucleic acids and cellulose were removed with benzonase and cellulase. Protein concentration was measured by absorbance at 280 nm (NanoDrop). Proteins were digested overnight with trypsin and Lys-C, each at an enzyme:protein ratio of 5:100. After centrifugation (10 min, 16,000 × *g*), digestion was quenched with TFA, the sample was diluted in 5% TFA in 2-propanol, and approximately 20 µg of peptides was loaded onto two-plug SDB-RPS StageTips, washed twice with 5% TFA in 2-propanol and once with 0.2% TFA in water, and eluted in 80% acetonitrile and 1% ammonium hydroxide. Peptides were concentrated by vacuum centrifugation, resuspended in 2% acetonitrile and 0.1% TFA and quantified again by NanoDrop. Approximately 200 ng of peptides was separated at 60 °C on a 40-cm in-house pulled and packed column (75 µm inner diameter; ReproSil-Pur C18-AQ 1.9 µm resin, Dr. Maisch GmbH) with buffer A (0.1% formic acid, 5% DMSO) and buffer B (0.1% formic acid, 5% DMSO, 80% acetonitrile) on an EASY-nLC at 350 nl/min with a 150-min method, increasing buffer B from 2% to 12% over 70 min, to 30% at 140 min and to 98% at 145 min, and analyzed on a Q Exactive HF-X. Full MS scans covered 300–1,650 *m/z* at a resolution of 60,000 (AGC target 3 × 10⁶, maximum injection time 25 ms). The 15 most intense precursors with charge 2–5 were isolated with a 1.4 Th window and fragmented by HCD at a normalized collision energy of 27. MS/MS spectra were recorded at a resolution of 15,000 (AGC target 5 × 10⁴, maximum injection time 25 ms), with dynamic exclusion for 30 s.

##### CAMPI SIHUMIx

CAMPI sample preparation and LC–MS/MS acquisition were performed as described in the original CAMPI study [1], whose methods and supplementary data files give the workflow-specific parameters. All used tryptic digestion, with sample preparation, LC gradients and mass-spectrometer configurations differing between the contributing laboratories. The analyzed files were acquired by DDA on Q Exactive HF-X, Q Exactive HF or Q Exactive Plus instruments.

##### PREDICT

PREDICT stool specimens were collected in the MUCOSA-PREDICT sub-study of the PREDICT cohort [2,5], within the MICROB-PREDICT consortium. Clinical metadata were obtained from the PREDICT study [2]. Sample preparation was as described above. PREDICT samples were analyzed on a Q Exactive HF-X using a 150-min LC program at 350 nl/min, increasing buffer B from 2% to 12% over 70 min and to 30% over the next 70 min, followed by an increase to 98% over 5 min and a 5-min hold. Full MS scans covered 300–1,650 *m/z* at a resolution of 60,000, with an AGC target of 3 × 10⁶ and a maximum injection time of 25 ms. The 15 most abundant eligible precursors were selected for HCD MS/MS with a 1.4 Th isolation window, normalized collision energy of 27, resolution of 15,000, an AGC target of 5 × 10⁴ and a maximum injection time of 25 ms. Dynamic exclusion was 30 s. Column, LC solvents and peptide amount (approximately 200 ng) were the same as for the healthy-stool samples.

##### CapScan

CapScan intestinal-fluid and stool samples were prepared and analyzed by DDA LC–MS/MS as described by Shalon et al. [3]. The analyzed files comprised intestinal fluid from sampling regions 1–4 and stool. These regions do not establish anatomical localization for individual specimens.

Full MS scans covered 300–1,650 *m/z* at a resolution of 60,000, with a normalized AGC target of 300% and a maximum injection time of 25 ms. Top-15 HCD MS/MS used a 1.4 Th isolation window, normalized collision energy of 30, resolution of 15,000, a normalized AGC target of 100% and a maximum injection time of 28 ms. Dynamic exclusion was 40 s. The embedded methods of all 347 original RAW files record a normalized HCD energy of 30, in agreement with the converted mzML values. This value and the MS1 maximum injection time of 25 ms differ from the 27 and 20 ms reported in the source publication [3]. Method records and the discrepancy are in Source Data.

#### Metagenomics and metatranscriptomics

PREDICT DNA and RNA were extracted from stool samples using the Qiagen AllPrep PowerFecal DNA/RNA Kit. Metagenomic sequencing libraries were prepared with the NEBNext Ultra II DNA Library Prep Kit, with a targeted insert size of 350–400 bp and dual-index multiplex oligos, on an automated liquid-handling system (Beckman Coulter i7 Series), and sequenced on an Illumina HiSeq 4000 with 2 × 150 bp paired-end reads. Metagenomes were assembled and genes predicted with the SPIRE pipeline (Methods). The resulting predicted-protein FASTAs were supplied by EMBL and used as inputs to FastaLake. Specimen-matched files provided the metagenome search comparators.

For CapScan, specimen metagenomes were assembled as described in the source study [3], with the software versions given in Methods.

CAMPI metagenome-derived proteins were supplied by the source study. FastaLake did not rerun assembly or protein prediction. The CAMPI metagenome pipeline and comparators are described in Supplementary Methods 3.

Ribosomal RNA was depleted from the extracted RNA with the NEBNext Bacteria rRNA Depletion Kit (New England Biolabs). Metatranscriptomic libraries were prepared and pooled with the NEBNext Ultra II Directional RNA Library Prep Kit (New England Biolabs) and sequenced single-end on an Illumina NextSeq 500. PREDICT metatranscriptomic profiles were supplied as companion molecular measurements for the cross-omics analyses. Their quantification and normalization will be described upon publication.

#### Metabolomics

Companion metabolite measurements were used for the PREDICT and CapScan functional cross-omics analyses. CapScan metabolomics was described in the source studies [3,4]. Cohort matching and profile transformations are described in Supplementary Methods 8. The PREDICT metabolomics protocol will be made available upon publication. Ethics approvals and consent are stated in Methods.

### Supplementary Methods 2 | Reservoirs, de novo prediction and exact database selection

#### Protein sequence reservoirs

FastaLake begins from a protein sequence reservoir, defined as the complete collection of protein sequences available for de novo-guided lookup. Reservoirs combined public microbial protein catalogs, human reference sequences and, where specified, proteins predicted from study-specific metagenome assemblies.

Reservoir compositions and sizes are given in Methods. Reservoir cleaning converted letters to uppercase and removed all other characters, including stop symbols, while preserving I/L distinctions. Each exact sequence was written once under the accession of its first source in the priority order given in Methods; alternative source accessions were retained in a provenance table. Source catalogs retained their own upstream processing.

In the assembly-inclusive PREDICT and CapScan reservoirs, the assembly-derived sequence space of an acquisition was not restricted to that participant’s or specimen’s own metagenome. Assembly protein names were not stable across specimens. Across the specimen metagenome FASTAs of the 296 PREDICT matched comparisons, 4,817,481 of 44,750,815 distinct sequences (10.8%) carried different names in different specimens, and 4,797,277 of the 4,918,663 names shared between specimens (97.5%) denoted different sequences. In the 295 CapScan comparisons, 2,393,928 of 29,163,863 distinct sequences (8.2%) occurred in more than one specimen under different names, and 2,494,126 of 2,494,236 shared assembly-local names (more than 99.9%) denoted different sequences. Each assembly-derived protein was therefore named PREDICT_MG_ or CAPSCAN_MG_ followed by the SHA-256 digest of its sequence. After renaming, no accession denoted two different sequences in either cohort, and the 53 sequences in each cohort that carried more than one accession were duplicate entries of the appended human Swiss-Prot (header audit tables in Source Data).

#### AlphaNovo de novo prediction

Archived top-ranked AlphaNovo predictions were used as sequence evidence [6]. The checkpoint was model-epoch12-step058020-peprecall0.5156.ckpt (SHA-256 68ff1bfa6e20a2eab7a32f2589d6354e4a0d9e7d41d38ff7dfcf999ca863eacd). None of the analyzed spectra were in the AlphaNovo training data. To estimate AlphaNovo accuracy, each reference-database Sage identification in the 15 CAMPI acquisitions (target, rank 1, peptide *q*≤0.01) was paired by scan number with the top-ranked AlphaNovo prediction and scored as correct when the unmodified prediction equalled the reference peptide, treating I and L as equivalent. Permuting predictions across spectra within each acquisition reduced the match rate to 0.006%. Archived inference code retained peaks at 50–2,500 *m/z*, removed peaks within 2 Da of the precursor, retained up to 800 peaks by intensity, sorted by *m/z*, square-root transformed intensities and applied L2 normalization. Constrained decoding generated 10 beam candidates. Only the top-ranked candidate was exported. FastaLake read peptide_prediction_detokenized_unmodified and score. Modified prediction strings were not additional lookup queries.

#### De novo prediction filtering and exact sequence lookup

Lookup eligibility required a positive score and 9–50 residues. The low-complexity filter removed sequences in which one residue exceeded 60%, sequences of at least 12 residues with at most two residue types, and exact alternating two-residue repeats of at least 10 residues. Complexity was assessed before I/L collapsing. The top 30% of eligible prediction rows were retained per acquisition, including score ties, and the retained query sequences were deduplicated with I/L equivalence. Exact lookup retrieved complete proteins and recorded query–protein–position hits, including overlapping matches, while retaining the original headers and I/L residues. The nine-residue minimum, top-30% fraction and complexity rules are configured analysis choices.

#### Relaxed lookup: order and mass ambiguity

De novo predictions carry residual order ambiguity that AlphaNovo records as bags, and near-isobaric composition ambiguity that a swap table can enumerate. We tested whether admitting either at the lookup step improved the final identification list. Over 48 acquisitions (15 CAMPI, 32 healthy stool and one CapScan), admitting recorded bags added 6,961 peptide–acquisition events and removed 5,941 (net +1,020, positive in 38 of 48). Restricting the bag route to predictions with at most 8 orderings, which covers 908,037 of 976,861 eligible prediction rows (93.0%), gave the largest net gain among the tested settings (+5,051, −3,318, net +1,733, positive in 37 of 48). Restricting it to at most 2 orderings was net negative (−793), so tighter bounds did not monotonically improve yield. Admitting mass-equivalent substitutions from the full swap table added 31,111 and removed 33,306 events (net −2,195, positive in 13 of 48). In the 32 healthy-stool acquisitions the mass arm was net −5,242 and positive in 3. In the first 60 completed acquisitions and two seeds, shuffled-query controls yielded no accepted peptides through the exact or bag route and 13,512 peptide–acquisition– seed observations through the mass route. Across the 645 of 726 acquisitions tested, the exact and bag routes accepted peptides in a single acquisition–seed, where a shuffled query selected abundant human titin (Supplementary Table 5). The separate matched comparison and fragment-audit protocol are detailed below and in Supplementary Fig. 2e,f.

#### Hierarchical FastaLake database construction

Sequence selection had three levels: reservoir, study-supported proteins and acquisition-specific candidates. Retained queries pooled across the declared study acquisitions retrieved the study-supported subset of the reservoir. Each acquisition’s retained queries then retrieved its candidates from that subset. Because its queries were in the pooled lookup, exact matching against this subset yields the same candidates as direct matching against the full reservoir under the same filtering rules. Selection parsimony reduced each candidate set to a FASTA of complete proteins for searching the original acquisition’s spectra. Both levels ran the same extractor with identical eligibility, top-30% and I/L settings. (software: MannLabs/fasta-lake, rust/fasta_extractor_v2/src/main.rs; analysis: fastalake-paper, pipelines/healthy_stool_stage1_lookup and healthy_stool_97_study_lookup) For the 58 healthy-stool acquisitions, re-derivation using only the final analysis population produced a study-supported protein space of 18,007,876 proteins. The original production run pooled 97 acquisitions, including the 39 excluded differential-centrifugation acquisitions, and produced 26,881,038 study-supported proteins. This re-extraction also produced identical acquisition-specific selected FASTAs for all 58 acquisitions. We report the 18,007,876-protein space for the 58-acquisition analysis.

#### FastaLake selection parsimony

Selection parsimony used the evidence defined in Methods, counting each I/L-equivalent peptide once. Unique support meant occurrence in exactly one protein of the complete candidate set. Unique and total distinct-peptide support were computed once for each protein and remained fixed. Evidence peptides were processed in decreasing mapping multiplicity, with lexical sequence ties. Each peptide was then assigned by the ranking given in Methods. The peptide–protein graph was obtained by matching an Aho–Corasick automaton of all eligible acquisition predictions against the I/L-normalized candidate sequences, recording overlapping occurrences. Under the accession-hash rule each assignment depends only on the fixed counts and the accessions, so the processing order does not change the selected proteins. The 64-bit FNV-1a hash is implemented in the code rather than taken from the Rust standard library, whose hash function may change between releases, so tie resolution is reproducible across builds. (software: MannLabs/fasta-lake, rust/parsimony_engine/src/main.rs, map_peptides_parallel, run_razor and fnv1a64) Ranking by unique evidence first assigns shared evidence to proteins already required by unique peptides. Hash tie-breaking avoids the catalog preference that lexical accession ordering alone could introduce.

A parser audit compared all 726 retained prediction files (114,971,345 rows) using the archived and current Rust parser functions and found identical eligible lookup rows and selection evidence within each stage. Three lookup sequences absent from selection evidence had no matches in the saved candidate FASTAs. Acquisition and cohort summaries and all eligible-row parser differences are in Source Data. (analysis: fastalake-paper, pipelines/prediction_parser_audit)

#### Bag-aware retrieval and validation

An AlphaNovo bag represents a predicted residue multiset whose internal order is unresolved in the prediction’s fragment-based representation. Our experimental matcher uses ordered query segments as anchors to find candidate protein locations, then verifies the ordered segments by exact sequence comparison and each bag by residue-multiset equality. A two-residue bag can therefore match either ordering without treating the alternatives as independent peptide observations. Evidence from matching routes is merged at the query level before the existing protein-selection procedure. This implemented bag-aware route differs from treating every possible ordering as a separately enumerated literal query. Mass-equivalent substitutions are evaluated separately because they can change residue composition as well as order. None of these routes changes the exact-lookup production analyses.

The 48-acquisition comparison contained seven search arms per acquisition: exact lookup, bags restricted to at most two, four or eight orderings, unrestricted recorded bags, mass substitutions, and the combined bag/mass route. Selection was rerun separately for each arm using the fixed-count procedure. Searches used the same spectra and Sage settings. Gains and losses were counted against the exact arm at nominal target-peptide *q*≤0.01 after canonical peptide deduplication within each acquisition. Summed counts are peptide– acquisition identifications. All 336 arm–acquisition rows, including losses, are in Source Data. The CapScan acquisition used the assembly-inclusive reservoir. The 48-acquisition panel and the separately selected six-acquisition fragment panel were not pooled.

The fragment audit compared gained bag-arm sequences with the best reported rank-one target sequence from the exact-search arm for the same spectrum. Among the composition-preserving rearrangement pairs, a diagnostic ion was a Sage-annotated b/y fragment spanning a residue-order difference whose mass did not match any modeled alternative-sequence b/y ion within 20 ppm, considering charges one and two and collapsing charge duplicates. Of 215 comparison pairs across six acquisitions (one CAMPI, one CapScan, three healthy-stool and one PREDICT), 193 had no such ion and 22 had at least one. (analysis: fastalake-paper, pipelines/bag_mass_lookup_search and bag_mass_paired_nulls)

### Supplementary Methods 3 | Reference databases and matched comparisons

#### CAMPI reference and metagenome databases

The supplied SIHUMIx reference and multi-omic co-assembly (SIHUMIx_MO, hereafter MG) databases were obtained from the CAMPI study [1]. The reference database combined UniProt proteomes selected using prior knowledge of the community. This database represented *Blautia producta* using proteins from other members of the genus, as a UniProt reference proteome for this species was unavailable when the database was constructed. The source study built MG from co-assembled metagenomic and metatranscriptomic reads. The source-study pipeline used IMP preprocessing, MEGAHIT 1.2.4, MetaBAT 2.12.1, MaxBin 2.2.6, DASTool 1.1.2 and Prodigal 2.6.3 [1]. No separate SIHUMIx RNA expression dataset was analyzed here.

The searched target FASTAs contained 29,673 reference and 19,435 distinct MG records, each including 116 contaminants. We removed supplied decoys before searching, and Sage generated new decoys. For FL + Reference, the complete reference target database was appended to each selected acquisition-specific FastaLake FASTA after selection parsimony. For FL + Reference + MG, the complete MG target database was subsequently added to this union. All 15 acquisitions were searched with the settings used for FL and the standalone reference and MG searches.

For each of the eight supplied member reference proteomes (29,557 member proteins), we assessed exact presence in the 375,333,654-sequence FastaLake reservoir with I and L treated as equivalent. Tryptic peptides of 9–50 residues without missed cleavages were additionally checked for presence in tryptic context.

#### Clinical matched-metagenome comparisons

The primary clinical database benchmark compared public-sequence FastaLake with specimen-matched metagenome databases. Both databases are defined in Supplementary Methods 1.

The 296 matched acquisitions were searched independently using the same standard Sage settings. We summarized search performance by canonical peptide counts and peptide-identity recovery, because independently inferred protein groups are not equivalent.

For the 295 matched CapScan acquisitions, the public-sequence FastaLake-to-matched-metagenome peptide-count ratio was summarized overall and separately for regions 1–4 and stool. Gains were calculated as in PREDICT (Supplementary Methods 5), not as ratios of cohort-median counts. The region breakdown was a descriptive follow-up of the public-sequence comparison, using canonical 5–50-residue peptides and their nested 9–50-residue subset at peptide *q*≤0.01 and the region labels of the analysis roster (Supplementary Methods 1). Source Data retain acquisition counts, peptide-identity recovery and a sensitivity summary that first averages ratios within specimen and then within participant and region. Repeated acquisitions were not treated as independent participants for inference. (analysis: fastalake-paper, pipelines/campi_comparator_fastas, campi_reservoir_containment, clinical_public_only_comparison, predict_curated_mg_comparator and capscan_region_mixed_model) For the primary regional inference, the outcome was log(FLᵢ/MGᵢ) from pooled cross-fitted counts in the 295 matched acquisitions from 15 participants. A linear mixed model included sampling group as a fixed effect, with stool as reference, and a participant random intercept, fitted by maximum likelihood with statsmodels 0.14.5. The mean of the four region coefficients defined the intestinal-fluid-versus-stool contrast, tested by a two-sided Wald z test; exponentiating its estimate and Wald interval gave the reported fold difference. Equality of the four region coefficients was tested with three independent region contrasts and a three-degree-of-freedom Wald chi-square test. Neither reported test received multiple-testing adjustment. The same-spectrum fit was retained separately. Candidate competition used Sage counts of candidates with at least one fragment match during preliminary scoring, summed over precursor-charge and isotope hypotheses and target/decoy labels. Counts were paired on identical rank-one spectra and summarized as cohort medians of acquisition medians.

### Supplementary Methods 4 | Database searching and peptide acceptance

Original mass spectra were searched using the official Linux release binary of Sage v0.14.7 [7], which reports its version as 0.14.6. Standard search settings are given in Methods. The preliminary fragment index used b/y ions with neutral theoretical masses of 200–2,000 Da and ion indices greater than two. The retained healthy-stool production configuration additionally records precursor charge range 2–4, fragment charge at most 2, minimum/maximum spectrum peak counts of 15/150, retention-time prediction enabled, and deisotoping, chimeric searching and wide-window mode disabled. Five PREDICT mzML files (MicrobPred_13, _44, _90, _115 and _181) had been truncated on disk. We reconverted them from the instrument RAW files with ThermoRawFileParser 2.0.0.0 after checking the RAW SHA-1. The main repaired searches used these reconversions with unchanged settings. The spectrum-split and 10-fold cross-fitting analyses used complete ThermoRawFileParser 1.3.4 replacements for the same five acquisitions (Supplementary Methods 7). The second-engine comparison excluded these five acquisitions. Standard Sage decoys used the rev_ protein-label prefix.

The exact representative configuration is supplied in Source Data. Other analysis routes have their own retained configurations. Label-free quantification (LFQ) used Hybrid peak scoring, Sum integration, a spectral-angle setting of 0.7, 5 ppm tolerance and combined charge states. Semi-enzymatic searches used the same configuration except for semi-enzymatic cleavage. Sage regenerated the configured ion series when scoring shortlisted candidates. The preliminary index mass and ordinal restrictions do not describe the instrument acquisition window or exclude those ions from final PSM scoring.

We retrospectively checked input completeness for the 653 public-sequence searches of 306 PREDICT and 347 CapScan acquisitions and the 58 acquisition-specific healthy-stool searches. Each mzML had to end with its closing tag, and the highest scan scored by Sage had to reach at least 90% of the scan range declared in the mzML. All passed (lowest coverage 99.6%).

The additional-confidence analysis retained the fixed 296-PREDICT and 295-CapScan matched acquisition rosters and all 30 broad-reservoir CAMPI fusion searches (15 acquisitions × two seeds). Added PSM posterior error probability (PEP) gates of 0.01 and 0.001 were evaluated on the same PSM as the peptide *q* gate before canonical union, and spectrum *q*≤0.01 and peptide *q* thresholds 0.005 and 0.001 were secondary sensitivities. Results were reported for lengths 5–50, 5–8 and 9–50. The stored posterior_error=1.0 default is an unavailable estimate rather than log_10_(PEP). The three PREDICT FastaLake acquisitions with such PSMs had zero PEP-filtered yields and remain in summaries. Undefined FL/MG ratios are kept in the roster without imputation. Primary identification and LFQ analyses retain their original acceptance rules (Source Data). Cross-fitted, joint healthy-stool and CAMPI member-specific PEP results are in Source Data. Across all 591 matched public-sequence FastaLake/metagenome acquisition pairs, the added PEP≤0.01 gate retained a median 85.7% of 5–50-residue FastaLake peptides in PREDICT and 84.7% in CapScan. At 5–50 residues, the cohort-median gains increased from 18.8% to 23.5% in PREDICT and from 49.3% to 59.9% in CapScan, but gains decreased in 11 PREDICT and 10 CapScan acquisitions. At 9–50 residues, cross-fitted gains rose from −3.5% to 1.7% in PREDICT, from −5.5% to −1.1% in CapScan stool and from 26.2% to 34.7% in intestinal fluid. (software: MannLabs/fasta-lake, fasta_lake/search.py, _dda_config; analysis: fastalake-paper, pipelines/sage_input_completeness and predict_truncated_mzml_repair)

### Supplementary Methods 5 | Protein inference, quantification and annotations

#### Study-level protein grouping and quantification

Post-search protein grouping started from the accepted PSM-reported peptide–protein membership graph, restricted to proteins in the FASTAs searched for the declared study and database arm. Grouping did not first remap every peptide to every protein containing that sequence. The later functional allocation used full peptide containment. The greedy cover and direct peptide assignment followed Methods. Each representative was checked to contain its assigned peptides, with I and L treated as equal. Local search members, representative presence in the acquisition FASTA and source provenance were retained separately. Representative-labeled group intensities are not estimates of uniquely identified proteins.

Analyses comparing database strategies used independently inferred study dictionaries constructed from identical acquisition rosters: 296 acquisitions for matched PREDICT and 295 for matched CapScan. The empty CapScan acquisition was retained as an explicit matrix column but excluded from analyses requiring a non-empty profile. Regional and functional quantitative summaries used the 346 non-empty CapScan acquisitions.

Protein-group counts from different database strategies are not one-to-one counts of the same biological proteins. For matched PREDICT and CapScan comparisons, percentage gains in peptide counts were calculated as the median within-acquisition percentage change, 100 × (FLᵢ/MGᵢ − 1), across the 296 and 295 matched acquisitions, respectively.

#### Origin composition

Origin classes were assigned per accepted LFQ feature from the annotated origins of all compatible proteins of its canonical peptide in the searched-target union plus the full local UHGG catalog. Origins came from UHGG genome lineages (GTDB release 202), from GMGC NCBI taxa and GMSC GTDB release 95 lineages mapped to release 202, and from UniProt organism identifiers mapped through NCBI taxonomy (snapshot of 30 September 2026). An NCBI taxon was mapped to the common ancestor of all GTDB release 202 genomes below it, no deeper than the rank of the source taxon. A release 95 lineage was mapped only when all of its genomes were present in release 202. Proteins from cohort or CAMPI assemblies, and catalog proteins that could not be mapped, had no origin annotation. For each feature, the origin was the deepest lineage shared by its annotated compatible proteins. Proteins without an annotation were ignored, and a feature without any annotated protein had unresolved origin. A feature was human-assigned when this lineage reached *Homo sapiens* at species rank. The other classes were microbial at species rank, microbial above species rank (Bacteria or Archaea), other annotated origins, and unresolved origin, which includes features whose annotated proteins disagreed at domain level. All five classes were retained in the total-intensity denominator. Fig. 5b combines the two microbial classes. Because unannotated proteins were ignored, 0.12% (healthy stool), 0.61% (CapScan) and 1.96% (PREDICT) of total LFQ intensity, pooled over all acquisitions, was human-assigned although some compatible proteins lacked an origin annotation. Acquisition-level fractions were averaged within patients or participants according to the experimental design. The protein-group comparisons in Fig. 3f use group-level classes, in which each feature was classed as host, microbial, mixed or unclassified from the protein members reported in the acquisition-level search. Relative LFQ shares describe measured signal.

The leading-species analyses in Supplementary Fig. 12b–f used a separate rule. Species–function allocation started from the species label of the study-group representative. This label was retained for an allocated KO share only if at least one protein containing the same peptide and carrying that KO resolved to the same species through the UHGG accession-to-species lookup. If no such carrier supported the representative label, the share was assigned to the unresolved category. Other compatible carriers, which could have different or missing species labels, did not by themselves veto a supported representative label. This is a carrier-supported representative label.

#### Functional annotation and profile construction

Functional annotations were indexed by full-length protein sequence, with I and L treated as equivalent, and combined from four sources. The first was eggNOG-mapper 2.1.12 in DIAMOND mode [8,9] (eggNOG database 5.0.2, DIAMOND 2.0.11, default settings) run on the searched proteins of each study in August and September 2026. The second was the MGnify per-genome eggNOG tables of the UHGG v2.0 species catalog (4,744 species representatives, with no record of the eggNOG-mapper version), joined by UHGG gene identifier and used only when DIAMOND aligned the protein to the annotation’s seed ortholog with at least half of the reported bitscore, because these tables list gene identifiers without sequences. The third comprised earlier eggNOG-mapper 2.1.12 runs on GMGC and GMSC catalog proteins, joined after sequence verification, for which the eggNOG database version was not recorded. The fourth was an eggNOG-mapper 2.1.12 run on 17,935 otherwise unannotated CAMPI reference proteins. Of 9,368,708 MGnify annotations, 372,277 (4.0%) failed this check, including 191,565 (2.0%) with no alignment. Most failures involved proteins shorter than half the seed ortholog or a seed shared with the adjacent gene. Values before and after this check are provided in Source Data. A protein kept every distinct annotation set from these sources. MGnify tables supplied annotations for 88–95% of compatible proteins, depending on the study. Each accepted LFQ feature’s canonical peptide defined its compatible proteins in the searched-target union plus the full local UHGG catalog. The five CAMPI arms shared one union. Distinct terms from all compatible proteins were collected at the requested level: KEGG orthologs matching K\d{5}, bacterial/archaeal scoped orthologous-group tokens, or pathways matching map\d{5}, excluding map011 global maps but retaining map012 overview maps. The feature intensity was divided equally among distinct eligible terms. Repeated support for the same term did not increase its share, and compatible proteins without an annotation did not remove terms supported by other proteins. Features without an eligible term remained functionally unassigned. In profiles excluding human-assigned signal, human-assigned features were withheld as a separate category. Allocated, unassigned and withheld intensity was checked against input intensity for every acquisition and level. Microbial single-OG profiles used only annotation sets that consisted of a single bacterial or archaeal scoped OG. A peptide could still have several eligible protein/term matches. The unassigned and withheld remainders were reported but excluded from functional distance calculations. As a sensitivity analysis, we allocated intensity only from features with a single eligible term, only from features whose compatible proteins were all annotated, or both. In profiles excluding human-assigned signal, these rules allocated a median of 7–61% instead of 62–81% of total intensity, depending on cohort and level. All 15 CapScan participants remained closer to their own stool (*P=1 × 10*⁻⁵), and PREDICT protein profile distances remained correlated with DNA, RNA and metabolite distances (*ρ*=0.16–0.64 against 0.16–0.51 with equal allocation, all *P*≤0.026), but their correlations with human-assigned protein profile distances fell from 0.43–0.51 to 0.28–0.35 (*P*≤0.002). In healthy stool, KO correlations still decreased from technical replicates (*r*=0.89–0.91) to different individuals, but the inter-individual value fell from 0.81 to 0.69–0.73, below the within-donor value, so the equal-allocation value for this design partly reflects terms shared through multi-term allocation. We calculated these sensitivity values before the seed-ortholog check, so they include the MGnify annotations that failed it.

The fixed-dictionary common-FASTA leading-species diagnostic is a separate group-based calculation. Each compatible protein group contributes its full intensity to each KO assigned to it, so totals across KOs are non-additive. This rule does not describe the PREDICT selected-feature heatmaps, whose microbial pathway rows use peptide-based equal allocation (Supplementary Methods 8). The clinical evidence-removal diagnostic also used a fixed group dictionary. Group intensity was partitioned into features with an accepted same-acquisition matched metagenome peptide and FastaLake-only features, then each partition was divided equally across the group’s distinct KO terms. Group-unannotated intensity remained outside normalized KO profiles but inside total intensity. KO support and normalized Bray–Curtis distance were compared before and after removal (Supplementary Fig. 8b–d). (software: MannLabs/fasta-lake, fasta_lake/study.py, group_searches; analysis: fastalake-paper, pipelines/peptide_centric_downstream, peptide_centric_functional_allocation, functional_allocation_sensitivity, mgnify_seed_ortholog_check, annotation_repair_coverage, campi_supplemental_eggnog, fixed_database_evidence_ablation, healthy_stool_common_fasta, healthy_stool_study_grouping and origin_annotation_share)

### Supplementary Methods 6 | Validation, error controls and spectrum inspection

#### Sequence controls and lookup specificity

We scrambled residues within each prediction of the representative healthy-stool acquisition, preserving length and composition, using seeds 20260901, 20260913, 20260914, 20260915 and 20260916. Authentic and scrambled queries were matched against the same 233,497,702-sequence reservoir. For each length, the numerator was distinct queries matching at least one protein and the denominator was all distinct queries in that arm/seed/length. All five seed curves and denominators are supplied. The full workflow on the representative acquisition propagated seed 20260901 through selection and the final search against the same spectra. Distinct-query counts were allowed to differ after scrambling and deduplication.

The original all-position shuffle did not fix the C-terminal residue. Its matched terminal-preserving counterpart fixed the last amino acid and shuffled the remaining residues while preserving full composition, length, row IDs, prediction scores and other fields, with the same five seeds. A fresh authentic workflow and all 10 shuffled workflows completed selection, selection parsimony and Sage with matched settings. Selected FASTAs contained 499–557 proteins for all-position shuffles and 489–536 for terminal-preserving shuffles, compared with 12,606 authentic proteins. Each shuffled arm yielded zero canonical target peptides at peptide *q*≤0.01. The authentic arm yielded 29,663. Both length-specific lookup denominators and final-search outcomes are retained separately (Source Data). Terminal-preserving query-match fractions were 8.11–8.65% at 9 residues and 0.72–1.03% at 10 residues, and no terminal-preserving query matched at lengths ≥13 in these realizations.

An adjacent-residue-swap sensitivity analysis tested every single adjacent swap of the distinct predictions from the representative acquisition against the full reservoir. This analysis used a separately documented scrambled control. The swap-aware lookup recovered additional authentic predictions but produced a disproportionate increase in candidate retrieval for scrambled queries. Full query and candidate counts are in Source Data. A separate candidate-clustering sensitivity used MMseqs2/Linclust at 99%, 97%, 95%, 90%, 75% and 50% sequence identity, with 80% bidirectional coverage and cluster mode 2, outside production selection.

#### Selected-database size and sequence support

We counted selected-database sizes as FASTA entries for all 726 primary acquisitions and compared with the 20,431 canonical entries of the reviewed human Swiss-Prot proteome (release 2026_03, isoforms excluded).

For the full-reservoir context, we used the same 726 selected FASTAs and exact reservoir entry counts (Supplementary Fig. 1 and Source Data).

We counted support for each selected protein separately from the top-30% production lookup queries and from the broader eligible predictions available to parsimony. Both sets used positive-score 9–50-residue predictions passing the complexity filter, with distinct I/L-equivalent sequences. Single-query support meant exactly one production query occurred in the protein. Corroboration meant that a protein contained a broader-set predicted peptide also accepted in the same acquisition’s final database search.

Selection sensitivity used the same 58 healthy-stool acquisitions for 18 distinct settings: minimum evidence lengths of 6–12 at top 30% and score fractions of 5–100% at minimum length nine, which share the nine-residue, top-30% setting; that setting was also run with the earlier 97-acquisition upstream pool. All 1,102 acquisition–setting observations, FASTA sizes and accepted counts are supplied. Here, peptide counts ignore modifications but do not merge I/L and are therefore not directly comparable with the primary peptide counts (Supplementary Fig. 2c,d). The nine-residue point reuses the production top-30% run.

#### CAMPI community recovery and peptide-loss analysis

We pooled accepted canonical peptides from each CAMPI search arm as set unions across the 15 acquisitions. A peptide was assigned to a community member when its I/L-equivalent sequence occurred in at least one protein of that member’s supplied reference proteome. A peptide was defined as member-specific when it occurred in exactly one of the eight supplied member proteomes. The 116 contaminant entries were excluded from community membership.

For every search arm and each of the eight members, we report the pooled member-specific peptide count, all per-acquisition counts including zeros, and the number of acquisitions with at least one such peptide. Both the accepted 5–50-residue population and its nested 9–50-residue subset are reported, retaining the original search *q* values. No minimum pooled peptide count is used to classify a member as recovered (Supplementary Fig. 3a,b).

Within each acquisition, the canonical peptide set from a search arm was intersected with the accepted reference set from the same acquisition and divided by the size of that reference set. For the pooled comparison, accepted peptide sets were united over all 15 acquisitions before calculating overlap with the 76,127 distinct pooled reference peptides (63,348 at 9–50 residues). For the CAMPI acquisition-depth analysis, we calculated the expected number of distinct accepted peptides when drawing 38,254 MS2 spectra without replacement from each acquisition, matching the smallest of the 15 acquisitions. We used each peptide’s observed number of accepted spectra to calculate its probability of appearing in the subsample, then summed these probabilities. These hypergeometric calculations kept the original identifications and *q* values fixed. Database selection and searching were not repeated.

To find where unrecovered reference peptides were lost, we traced every reference peptide accepted in an acquisition but not in its FastaLake search through database construction, counting each as one peptide– acquisition observation. We scanned candidate and final selected FASTAs for the exact I/L-equivalent sequence and checked each occurrence against the production tryptic search rules. Observations were classified into mutually exclusive states according to whether the sequence was absent from the candidate set, present only in candidate proteins removed during selection parsimony, present only in a non-compatible peptide context, or present in a compatible context in the final FASTA but not accepted by the final search. The complete stage counts and direct-prediction availability are in Source Data. For each acquisition, we compared the original FastaLake search with searches after adding the reference proteomes, with or without MG. For every gained peptide, we checked whether its sequence was absent from the original selected database, present only in a context incompatible with the digestion rules, or already searchable. For peptides quantified only in the expanded search, we recorded whether they were newly identified, previously identified but newly quantified, or absent from that search’s accepted peptide set. We retained gains and losses as peptide–acquisition observations (Source Data).

We matched each retained top-ranked CAMPI prediction to the accepted reference identification of the same spectrum, checking precursor masses and reconstructing the production query set. Where they disagreed, we recorded whether the reference peptide remained searchable in the final FASTA, was accepted elsewhere in the acquisition or on the same spectrum, and whether the discrepant prediction itself retrieved a reference-compatible annotated family. Unannotated candidates were not classified as unrelated. These endpoints distinguish retention through combined evidence from retrieval by a discrepant prediction.

#### Candidate clustering sensitivity

To test the effect of reducing candidate sequence diversity before FastaLake selection, we clustered candidate proteins from each CAMPI acquisition with MMseqs2/Linclust at six sequence-identity thresholds with 80% bidirectional coverage. Cluster representatives replaced the original candidate set before otherwise unchanged FastaLake selection parsimony. Clustered and unclustered databases were searched with the same Sage configuration. Accepted canonical peptides were compared at peptide *q*≤0.01. KO recovery was the fraction of KOs compatible with the unclustered accepted peptides that remained compatible with the clustered accepted peptides, using one fixed peptide-to-KO mapping. Candidates were sorted by accession before MMseqs2 createdb (--shuffle 0), and linclust used --min-seq-id 0.99, 0.97, 0.95, 0.90, 0.75 or 0.50, -c 0.8, --cov-mode 0 and --cluster-mode 2. All six clustering arms were compared with the same fixed contemporaneous unclustered baseline. Accepted identities, all 105 acquisition–arm configurations and paired counts are supplied in Supplementary Table 1.

#### Second search engine

We also searched the 15 acquisition-specific FastaLake, reference and MG databases with MSFragger 4.1 followed by Percolator 3.06.5. Digestion, peptide length and mass bounds, mass tolerances, isotope errors, fixed and variable modifications and the minimum number of matched fragments were matched as closely as possible to the Sage configuration. Mass calibration, deisotoping and MSFragger’s default N-terminal acetylation were disabled. One full-length reversed decoy was supplied per target, and peptide confidence was estimated independently per acquisition and engine at *q*≤0.01. Comparisons used canonical 9–50-residue peptide sequences, with 5–50-residue values in Source Data. Percolator used target–decoy competition with seed 913. This analysis tested whether the selected FASTA transferred to a second search workflow. The same MSFragger and Percolator workflow was applied to the matched public-sequence FastaLake and metagenome searches of both clinical cohorts and to the healthy-stool FastaLake searches.

#### Spectrum-level comparison

For Fig. 4d, the same-spectrum public-sequence FastaLake and matched metagenome databases were searched again for 5–50-residue peptides with Sage v0.14.7 using the production settings, quantification disabled and five candidates reported per spectrum. For each spectrum, we selected the highest-hyperscore reported target row. A spectrum was classified as FastaLake-only when the selected FastaLake target had peptide *q*≤0.01 and the selected metagenome target was absent or had peptide *q*>0.01. Spectra with accepted targets in both searches were excluded from this denominator irrespective of whether the peptide identities agreed. The metagenome database was additionally digested in silico under the standard cleavage, missed-cleavage, length and mass rules (Supplementary Methods 4), including fixed carbamidomethylation of cysteine but without expansion of variable modifications. Protein termini were permitted.

FastaLake-only spectra were classified into mutually exclusive categories according to whether the FastaLake peptide sequence was absent from the metagenome FASTA, present in a metagenome protein but outside the theoretical digest, present in the digest with no reported metagenome target, or present in the digest with the best reported metagenome target representing either the same or a different peptide sequence. In the latter two target-containing categories, the selected metagenome target had peptide *q*>0.01. Presence in the theoretical digest establishes sequence availability under these rules but does not show that the FastaLake peptide was generated, retained or scored as a candidate for that spectrum.

#### Entrapment and selection-exposure controls

Because FastaLake uses spectral evidence to select the database subsequently searched against the original spectra, we introduced controls at different workflow stages and analyzed them separately.

##### CAMPI target–entrapment fusion control

The defined CAMPI community was used for an artificial target–entrapment fusion control following the linked-retention principle of Yi and Fu [10]. For the broad-UniProt CAMPI benchmark, every target protein in the 375,333,654-sequence reservoir was paired with a seeded control in which residues were permuted within each tryptic segment, keeping K, R, P and the first residue of each segment in place. Target and control entered before FastaLake sequence selection and were retained together through cohort selection, acquisition selection and parsimony before being searched as separate records using the production Sage configuration and its internal decoys. This linked-retention control characterizes error behavior in the artificial fusion system and does not directly estimate the false-discovery proportion in ordinary cohort searches.

At each *q* threshold and length range, the fusion numerator comprised accepted paired entrapments E, accepted entrapments whose paired target was not accepted E₀, twice the accepted paired entrapments with a better or tied peptide *q* value than their accepted target B, and all accepted sequences outside the audited one-to-one paired subset U. The diagnostic was (E + E₀ + 2B + U)/N, where N was the accepted canonical sequence count. Unresolved, shared and non-bijective cases contributed to U rather than disappearing. Score-ordering sensitivity was retained separately. Cohort summaries sum acquisition-level numerators and denominators before division, rather than pooling distinct-peptide unions. Both seeds, nested 9–50 and 5–8 populations, *q* thresholds and components are supplied in Supplementary Fig. 5. At peptide *q*≤0.01 across the two seeds, the pooled diagnostic was 2.59–2.79% for 5–50 residues and 1.55–1.56% for the nested 9– 50-residue subset. The diagnostic is the sum of a paired contribution, (E + E₀ + 2B)/N, and an unresolved contribution, U/N. For peptides of 5–8 residues, the paired contribution was 3.09–3.38% and the unresolved contribution 6.35–8.30%, giving 9.72–11.39% in total. Unresolved pairing accounted for 65–73% of that total. At 9–50 residues, the paired contribution was 1.10–1.20% and the unresolved contribution 0.36–0.45%. Unresolved pairing can reflect shared target/control strings.

The healthy-stool, PREDICT and CapScan fusion controls used their own cohort reservoirs and the same linked-retention design. They are compared descriptively with the broad-reservoir CAMPI control.

##### Reservoir occurrence of accepted control sequences

Accepted entrapment-only sequences from the CAMPI fusion searches were restricted to peptide *q*≤0.01, lengths 5–50, and deduplicated across acquisitions and seeds after treating I and L as equivalent. Exact substring occurrence was scanned across all 375,333,654 original reservoir proteins, treating I and L as equivalent on both sides. Composition-preserving shuffled sequences were generated from this unique accepted-control set with seed 20260925. All 4,000 positive sequences sampled from original reservoir proteins were recovered. Among 5–8-residue sequences, 1,517/1,544 accepted controls (98.25%) and 1,511/1,543 shuffled sequences (97.93%) occurred in the reservoir. At 9–50 residues the corresponding counts were 480/3,177 (15.11%) and 350/3,177 (11.02%).

Additional-confidence fusion summaries used the same accepted-PSM filters and canonicalization as the ordinary-search sensitivity, retaining the two seeds separately. The weighted paired numerator, accepted unresolved identities and full augmented canonical denominator were recomputed at each fixed reporting gate. Conditioning on the added PEP gate changes the discovery set (Source Data). Peptides of 5–8 and 9– 50 residues are reported separately because short sequences recur more readily in large reservoirs and complicate target–control pairing. With the added PEP≤0.01 gate, the diagnostic across the 30 CAMPI fusion-control searches decreased to 0.666–0.686% for 9–50 residues but remained 5.34–6.90% for 5–8 residues.

##### Donor-selected entrapment

The donor-selected assay evaluated the search step after selecting the database independently from the spectra being evaluated. Each of the 58 healthy-stool acquisitions was searched against a database selected from another acquisition of the same experimental design. This transferred database was fixed before control construction. FDRBench v0.0.4 [11] generated composition-matched shuffled entrapment peptides and explicit shuffled decoys after in silico digestion of the transferred database. Sage decoy generation was disabled for this control, and the supplied FDRBench decoys were identified using the rev_ tag. Searches otherwise used the production mass-spectrometric settings with digestion disabled because explicit peptide records were supplied. At peptide *q*≤0.01, the paired estimator was calculated according to FDRBench as (E + E₀ + 2B)/(T + E), where T and E are accepted targets and entrapments, E₀ denotes accepted entrapments whose paired target was not accepted, and B denotes entrapments ranked above their accepted target by peptide *q* value and then PEP [11]. No entrapment peptide received exactly the same score as its paired target, so tie counting did not affect these estimates.

The 9–50-residue analysis was a post hoc restriction of the accepted 5–50-residue search and retained the *q* values assigned in that search. Three seeds were averaged within target acquisition before acquisition-level summaries. The five self-selected controls remain a separate stratum.

##### Restricted-reference simulation

Whole-procedure error was counted directly only in a restricted simulation in which generating peptide sequences were known. AlphaPeptDeep-predicted spectra for 2,048 generating peptides per acquisition were noise-injected to create 12 baseline and 12 stress mixtures. Each mixture was searched against a fixed reference, a database selected from the measurement being searched and a database selected from an independently noised replicate. Generating peptides were 7–30 residues and originated from an approximately 30,000-protein CAMPI reference. Searches accepted peptides of 5–50 residues. For each target canonical-peptide list passing peptide *q*≤0.01, false discoveries were counted directly against the known generating sequences. The false-discovery proportion was the number of false discoveries divided by the larger of the total number of discoveries and one. Means and 95% bootstrap intervals were calculated over whole mixtures, separately for the 12 units in each regime (20,000 bootstrap resamples). These simulations are controlled stress tests. All 72 ordinary-search outcomes were retained and form a defined subset of 288 total simulation outcomes, which also include fragment interventions. Mean false-discovery proportions at the nominal 1% peptide threshold were 1.25% for the fixed-reference search, 1.06% with selection from the replicate and 1.01% with selection from the measurement being searched under baseline conditions, and 1.50%, 1.20% and 1.91%, respectively, under stress conditions. Stress conditions combined increased spectral noise with greater reference incompleteness: 50% of generating peptides were deliberately absent from the search reference, compared with 25% at baseline. Under stress, selection from the measurement being searched increased the proportion by 0.71 percentage points relative to selection from the replicate (95% bootstrap interval, 0.35–1.12) and by 0.41 percentage points relative to the fixed-reference search. (analysis: fastalake-paper, pipelines/database_selection_specificity, cohort_database_sizes, scramble_end_to_end_workflow, cohort_prediction_support, healthy_stool_selection_sweep, campi_reference_gap, campi_variability, campi_near_miss_audit, campi_candidate_clustering, campi_second_engine, predict_fl_only_spectrum_audit, campi_fusion_375m, cohort_fusion_controls, healthy_stool_donor_entrapment and restricted_reference_simulation)

### Supplementary Methods 7 | Transfer, pooling and common study search spaces

#### Healthy-stool database transfer and pooling

For each of the 58 healthy-stool acquisitions, we selected transferred databases from the predictions of another acquisition. A same-design donor was selected from the other acquisitions belonging to the target’s experimental design group (Supplementary Methods 1). An inter-individual control donor was selected from the 10 inter-individual acquisitions. Fig. 3b,c labels these arms Same series and Inter-individual series. For the 10 inter-individual targets, both arms used another participant’s database, and the two donor assignments differed for nine targets and coincided for one. Each transferred database was selected using the donor acquisition’s retained predictions against the healthy-stool study-supported protein space, followed by the production FastaLake selection-parsimony rule, and was then searched against the target acquisition’s spectra with the standard Sage settings and quantification disabled. Count retention and identity recovery (Methods) used canonical target peptides passing peptide *q*≤0.01, reported at 9–50 residues (5–50 residues in Source Data).

For technical-replicate pooling, selected FASTAs were merged by accession, requiring repeated accessions to carry identical sequences and headers. For each target acquisition, the other-11 database contained the union of selected FASTAs from the remaining 11 technical replicates and excluded the target’s own final FASTA.

Artificial shuffled proteins used to assess control acceptance entered the reservoir before sequence selection and underwent lookup and parsimony with ordinary reservoir sequences. A shuffle-only accepted peptide was one assigned exclusively to shuffled-control proteins and without an exact I/L-equivalent occurrence in the original reservoir. Counts were summarized as peptide–acquisition observations because the same sequence identified in two acquisitions contributed twice. Shuffle-only peptides accounted for 2 of 376,412 accepted peptide–acquisition observations with acquisition-specific databases and 12 of 411,354 with the other-11 unions (Supplementary Table 6).

For the 58 healthy-stool acquisitions, we built KO profiles from accepted quantified peptide features. Intensity was allocated equally among compatible KO annotations as described in Supplementary Methods 5. We generated profiles with all origins and profiles excluding human-assigned signal. Human assignment followed Supplementary Methods 5. The second profile retained unresolved origin. We recorded signal without eligible KO annotations separately and excluded it from profile comparisons.

We compared acquisition pairs within each healthy-stool sampling design using KOs positive in both acquisitions, without imputation. For each pair, we calculated Pearson correlation on log_2_ intensities and the median per-KO CV, defined as the population standard deviation of the two raw intensities divided by their mean and multiplied by 100. Fig. 3f reports medians across pairs. Coverage, intensity-conservation checks, feature gains and losses, and both comparisons are in Source Data.

#### Common study search space

To test a common search space across the healthy-stool series, we deduplicated the 58 acquisition-selected FASTAs by exact full-length sequence to give a union of 153,267 protein sequences. This union of selected search FASTAs is distinct from both the upstream reservoir and the study-supported protein space. In the acquisition-specific healthy-stool analysis, 43.2% of quantified peptide features, counted per acquisition, were assigned to a representative accession absent from that acquisition’s own FASTA, although the supporting peptide was identified in that acquisition. Shared protein-group labels therefore need not reflect identical sequence spaces across acquisitions. The clinical common-FASTA sensitivity used the original 306 PREDICT and 331 CapScan acquisitions, with separate per-acquisition searches and downstream directLFQ. The 331-acquisition CapScan union contained 452,255 proteins and included participant 15’s stool measurements but excluded that participant’s 16 intestinal-fluid acquisitions, which were linked later. The expanded 347-acquisition union contained 460,977 proteins and was not the database used for this sensitivity. Clinical primary analyses retained acquisition-specific FASTAs; the common-search comparison was not extrapolated to the expanded roster.

#### Spectrum-split selection sensitivity

The fixed roster comprised 58 healthy-stool, 15 CAMPI, 306 PREDICT and 347 CapScan acquisitions. We divided predictions by native scan-number parity. Scan IDs, precursor *m/z* and charge were verified against mzML spectra. Separate cohort evidence caches were rebuilt from the original full reservoirs for each half, with the unchanged top-30% eligibility procedure, 9–50-residue lookup and I/L equivalence. The all-prediction arm used the all-prediction selected FASTA, the same-half arm used odd predictions and the disjoint arm used even predictions. All three Sage v0.14.7 searches scored the same odd MS2 subset while retaining MS1 spectra. Confidence was estimated separately in each search.

Rank-one targets with native peptide *q*≤0.01 were collapsed by modification-stripped I/L-equivalent peptide sequence. Count and identity-overlap endpoints were reported for 5–50 and nested 9–50 residues without re-estimating *q* values. A participant-weighted sensitivity used the median ratio within each participant and then the median across participants: 93 PREDICT patients (302 mapped acquisitions, four missing IDs) and 15 CapScan participants (all 347 acquisitions).

For both the spectrum-split experiment and 10-fold cross-fitting, five truncated PREDICT inputs were replaced with complete ThermoRawFileParser 1.3.4 files carrying matching RAW SHA-1 declarations. Their peak arrays differ from both the original ProteoWizard 3.0.24025 files and the ThermoRawFileParser 2.0.0.0 reconversions used for the main searches (Supplementary Methods 4). Every comparison arm for a recovered acquisition used the identical replacement. A sensitivity analysis excluding these five acquisitions is in Source Data.

Native parity does not ensure equal numbers of selection predictions, repeated peptide observations can occur in both halves, only even-select/odd-score was tested and clinical arms are public-sequence. Thirteen Sage arm searches across eight acquisitions used heuristic scoring after model-fit failure. Two low-yield CapScan disjoint arms had zero accepted peptides. Both are retained. Excluding fallback-affected acquisitions barely changes cohort medians.

The Source Data contain the acquisition-level counts and identity overlaps, cohort and participant summaries, input-balance strata, conversion and fallback sensitivities. Results are shown in Supplementary Fig. 6.

#### Joint healthy-stool search

The three healthy-stool search designs are defined in Methods. Peptide-count comparisons required a rank-one target peptide accepted at peptide *q*≤0.01 in the corresponding acquisition. For per-acquisition confidence, the joint PSMs were divided by acquisition. Within each acquisition, the best discriminant score was taken for each modified peptide sequence and label, *q* was calculated as the number of decoys divided by the number of targets at or above each score and made monotone, and rank-one target peptides with *q*≤0.01 were accepted. Applied once across all 58 acquisitions together, the same estimator gave a median gain of 32.0%, compared with 32.9% from Sage, for peptides of 5–50 residues (34.2% from Sage at 9–50 residues). In one permutation of target and decoy labels within each acquisition, it accepted at most 17 peptides in any acquisition. With one added to the decoy count, it accepted none. Quantified features were then gated by these per-acquisition identifications in the same way as for study-level confidence. Quantitative comparisons likewise required a positive finite LFQ feature with an accepted canonical peptide in that acquisition.

The joint analysis used Sage v0.14.7 rebuilt from source with one correction in the precursor-level LFQ false-discovery step: *q* values were stored by precursor and target/decoy status rather than by precursor alone. Without this correction, a target precursor and its decoy share one entry, so the target can receive the decoy’s *q* value. This can only raise target LFQ *q* values. Among the existing LFQ outputs it changes only the q_value column. Intensities and PSM, peptide and protein *q* values are unchanged. Reported identifications and quantitative matrices were filtered on Sage peptide *q* values from the same acquisition, not on protein or LFQ *q* values. Sage protein *q*≤0.01 was applied only in the separate comparison of native Sage protein sets. With study-level confidence, the original and corrected joint searches produced identical peptide counts, identity-retention, completeness and reported precision results.

All-replicate CV was sample standard deviation divided by mean intensity across 12 technical or 15 workflow replicates, restricted to features complete in both arms. Pairwise protein-group CV used population standard deviation divided by mean raw intensity, and Pearson correlation used log_2_ intensity. Primary pairwise precision compared groups quantified in both acquisitions under both strategies without imputation; group CVs were summarized by the median within each acquisition pair and then across pairs. The 66 technical and 105 preparation pairs shared acquisitions and were not treated as independent biological replicates. For all-replicate peptide-feature CV, charge states of the same modified peptide were combined. The own-row definition required positivity in both acquisitions under the arm being scored. The matched definition additionally required positivity under both arms in both acquisitions. Matched group labels need not have identical peptide contributions. Both definitions and all-replicate CV are retained in Supplementary Fig. 7 and its full table. Using microbial protein groups observed under both analysis strategies within each acquisition pair, the joint search reduced median pairwise CV between technical replicates from 20.1% to 15.2% and increased Pearson correlation from 0.83 to 0.87. Among independently prepared samples, however, CV increased from 32.9% to 35.1% and correlation decreased from 0.76 to 0.72. With study-level confidence, the corresponding changes were from 20.3% to 14.5% and 0.83 to 0.87 for technical replicates, and from 33.1% to 33.6% and 0.75 to 0.74 for independent preparations (Supplementary Fig. 7c,d and Source Data). We evaluated leading-species agreement with study-level confidence. Using a fixed functional annotation dictionary and matched resolved functions, function-averaged pairwise agreement of the leading annotated species remained 74.9% among technical replicates under both strategies, but decreased from 70.8% to 66.5% among independent preparations.

#### 10-fold spectrum cross-fitting

Fold membership was assigned deterministically from the native scan number (Methods); the exact fold key is given in the code and Source Data. Folds held 9.9–10.2% of MS2 spectra in the six pilot acquisitions. For each cohort and fold, out-of-fold prediction tables of all 306 PREDICT and 331 CapScan acquisitions were written (the 16 participant-15 intestinal-fluid acquisitions added later were outside this original analysis roster; participant 15’s stool acquisitions were included) and used to rebuild the cohort evidence cache from the original reservoir with the unchanged top-30% eligibility, 9–50-residue lookup and I/L equivalence. Per-acquisition selection used selection parsimony on the acquisition’s out-of-fold predictions. For the extension to 295 matched CapScan acquisitions, the 282 original cross-fitted results were retained and the 13 added acquisitions used fold caches built from all 347 acquisitions; in a control, caches built from 331 and 347 acquisitions gave identical pooled selections apart from Sage per-fold numerical variation (Source Data). No prediction from a fold-k spectrum entered the fold-k selection. FastaLake, all-prediction and matched metagenome databases were searched with Sage v0.14.7 and the archived per-acquisition configuration, changing only database, input and output paths.

Because each Sage search fits its own discriminant, per-fold scores cannot be pooled directly. For each acquisition and database, we therefore pooled the fold PSMs and rescored them with a reimplementation of the Sage v0.14.7 source (version string 0.14.6). The reimplementation recomputed Sage’s 20 PSM features, including a kernel-density precursor-mass error probability, fitted Fisher’s linear discriminant between the pooled target and decoy PSMs with Sage’s Gauss–Jordan solver and power-iteration eigenvector, oriented it so that targets scored above decoys, and assigned picked-peptide *q* values to the pooled scores. Peptides were counted as rank-one targets with pooled peptide *q*≤0.01, modification-stripped and I/L-equivalent, at 5– 50 and nested 9–50 residues. On 36 archived unsplit searches, the *q* value reimplementation reproduced Sage’s accepted sets in 36 of 36 and the full rescoring in 31 of 36. With one fold (no cross-fitting), selection reproduced the archived databases in six of six pilot acquisitions, and Sage reproduced the original accepted peptide sets in 24 of 24 comparisons (two databases and two length ranges). Through the pooled rescoring, 22 of 24 were identical. The two exceptions were one PREDICT metagenome search, with 3,047 against 3,100 peptides at 5–50 residues and 2,361 against 2,429 at 9–50 residues. Across all 591 acquisitions, the fold-pooled all-prediction database reproduced the original gains over the metagenome search, with median differences of at most 0.04 points in every cohort stratum.

Sage’s discriminant solve is numerically unstable on these data, and a one-ulp perturbation of its inputs can change scores. The five differences in the rescoring check and the two in the one-fold control arise from this instability. Pooled rescoring ran in Python 3.10.9 with NumPy 1.26.4. Two one-ulp perturbations of the pooled inputs changed stratum medians by at most 0.16 points but changed 25 and 28 of 591 acquisitions (4–5%) by more than 0.5 points at 5–50 residues. In three acquisitions (two PREDICT and one CapScan region 3), the pooled fit was singular and Sage’s heuristic score was used for the FastaLake arm, and in the two PREDICT acquisitions also for the all-prediction arm, whereas the metagenome arm used the fitted discriminant. An independent closed-form discriminant, which reproduced the accepted sets exactly in five checked acquisitions without fallback, gave gains of 75.1%, 13.3% and 11.7% at 5–50 residues in these three acquisitions instead of 31.5%, −10.7% and −74.3%. Substituting these values changed the PREDICT medians from 6.50% to 6.55% at 5–50 residues and from −3.54% to −3.49% at 9–50 residues and left every CapScan median unchanged. The reported values keep the fallback scores. Gains are acquisition medians with 95% intervals from 10,000 participant-level bootstrap resamples (seed 20261004), in which the four PREDICT acquisitions without a patient ID were resampled as separate units. Sampling-region intervals come from a separate bootstrap stream with the same seed, so CapScan stool has a slightly different 9–50-residue interval there (−8.4 to −2.5% instead of −8.4 to −2.3%). The union of per-fold accepted peptides with native per-fold *q* values changed the cohort, stool and intestinal-fluid medians by at most 1.4 points and single sampling-region medians by up to 2.9 points. In two CapScan acquisitions every fold’s metagenome search accepted no peptides, so their union gain is undefined and they are excluded from that sensitivity. Repeated observations of a peptide in other folds can still support its selection, the pooled rescoring uses retention-time features fitted per fold, and the matched metagenome searches had no separate error control. With the cross-fitted databases, more FastaLake than metagenome peptides were identified at 9–50 residues in 106 of 296 PREDICT, 16 of 56 CapScan stool and 213 of 239 intestinal-fluid acquisitions. On the same spectra, the cross-fitted databases identified a median 9.8% (PREDICT) and 14.1% (CapScan) fewer 5–50-residue peptides than the all-prediction databases, a difference that combines the same-spectrum advantage with the withheld tenth of the selection evidence. An earlier comparison that selected databases from half of the predictions, on the other half of the spectra, gave lower gains of −1.0%, −10.5% and 21.7% at 5–50 residues (Source Data). Selection from odd-scan predictions, which also used half of the predictions but came from the scored spectra, gave median gains of 12.4% in PREDICT, 11.6% in CapScan stool and 50.0% in intestinal fluid at 5–50 residues. For the same-spectrum comparison, requiring peptides of at least seven residues, with confidence re-estimated in every search, reduced the median gains to 14.5% (PREDICT) and 43.8% (CapScan), while identity recovery remained 85.4% and 84.0% (Supplementary Fig. 4). Peptides of 5–8 residues, which had the highest fusion-control diagnostic (Supplementary Fig. 5), showed gains of 67% and 94%. (analysis: fastalake-paper, pipelines/healthy_stool_database_transfer, healthy_stool_pooling_controls, healthy_stool_common_fasta, healthy_stool_joint_search_q, healthy_stool_ko_precision_refit, cohort_selected_fasta_unions, cohort_spectrum_split_selection, metagenome_split_spectra_comparison, capscan_crossfit_10fold and minlen7_search_sensitivity)

### Supplementary Methods 8 | Clinical, regional and cross-omics analyses

#### Leading functional contributors

For healthy stool, a function was eligible for leading-species analysis when a leading species was resolved in at least 80% of acquisitions in every experimental design. Unresolved observations were not counted as a species. For the common-FASTA comparison, leading-species agreement was evaluated with the fixed study annotation dictionary, requiring resolved leaders in at least 80% of acquisitions in each of the two compared arms within the experimental design. Pairwise agreement was calculated from acquisitions resolved in both arms for each eligible KO, then averaged equally over KOs. Comparisons under native search-engine protein inference are also in Source Data.

For CapScan, function intensities were normalized to acquisition total LFQ, averaged within specimen and then within participant and region. A function had a resolved leader when species-annotated non-host intensity was positive. The leading species was the largest supported species-assigned contribution, with lexical tie-breaking. Eligibility required a resolved leader in at least 12 of the 15 participants. Species-label coverage was assessed separately across 21,168 eligible function–participant cells spanning 1,467 functions under direct peptide assignment. The denominator retained unresolved functional allocation. Label coverage is therefore the fraction of functional intensity carrying a supported species annotation. Intestinal fluid–stool leading-species agreement was additionally evaluated on function–participant–region comparisons with a resolved leader in both the intestinal region and stool. A stricter subset required at least 20% species-resolved signal and a leading-species margin of at least five percentage points of resolved intensity in both observations.

The Fig. 5e mosaic displays 24 functions selected previously by abundance for visualization. Among these 24 displayed functions and the 15 CapScan participants, a participant–function profile was evaluated when stool and at least one intestinal sampling region had a resolved leader. Persistence was defined as the stool-leading species remaining the leader in every resolved intestinal sampling region. This 30.8% therefore refers only to the 24 displayed functions.

#### PREDICT biological and clinical analyses

For biological analyses, we used the assembly-inclusive PREDICT production searches, whose reservoir is described in Methods and Supplementary Methods 1. These analyses are distinct from the public-sequence benchmark.

Human-assigned and other origin fractions of total LFQ intensity were calculated per acquisition as described in Supplementary Methods 5. An acquisition was on antibiotics if any core antibiotic field or antibiotic-treatment field was positive and off if all nine core antibiotic fields (prophylaxis for spontaneous bacterial peritonitis, variceal bleeding or an intervention, rifaximin, norfloxacin, cephalosporins, penicillins, other antibiotic and antibiotic for infection) were negative. Other records were retained as unknown. A separate field recording rifaximin for hepatic encephalopathy was not used in the primary definition. In a sensitivity analysis it was counted as exposure, which reclassified 12 acquisitions from seven patients from off to on and changed the paired population to 40 patients. A second sensitivity excluded all patients with any such record, leaving 35 of the 42 paired patients, and a third set only the 12 off-state acquisitions with such a record to unknown, leaving 38 patients. All used the same resampling settings and seeds as the primary analysis (Supplementary Table 2). Counting rifaximin for hepatic encephalopathy as exposure, the human-assigned share was 11.5 percentage points higher during exposure (95% CI 5.7–17.7, *q*=4.8 × 10⁻⁴) and quantified protein groups were 2,378 fewer (95% CI 1,140–3,651, *q*=5.4 × 10⁻⁴).

Patient-bootstrap intervals (Methods) used seed 20260907, and sign-flip tests used seed 20260907 plus the endpoint index. The two-test Benjamini–Hochberg [12] family was applied separately for the primary definition and each sensitivity definition.

#### Enterotype Dysbiosis Score and clinical heatmap

The supplied metagenome-derived Enterotype Dysbiosis Score (EDS), enterotype labels and membership-strength fields were retained following Keller et al. [13]. The molecular-feature overview (Fig. 4h) retains a fixed list of 20 features from an earlier cohort screen; HV366 had no detections in the 85 displayed patients and was omitted, leaving 19 rows. The retained feature list, current input matrices and deterministic display code reproduce its displayed values. For both patient-level displays and the subsequent biochemical screen, the earliest matched proteome–metabolome observation was retained for each of 85 identified patients. Display columns were ordered by continuous EDS. KO and pathway signals came from the assembly-inclusive peptide profiles, with intensity divided equally among distinct eligible annotations from compatible proteins in the searched-target union plus the full local UHGG reference. Human-assigned signal was excluded and unresolved origin was retained. Features required positive finite measurements in at least 43 patients for continuous association testing. Non-detections were not imputed. We tested Spearman associations with EDS and between metabolites and proteins linked by compound–reaction–KO or compound–pathway annotations. Sensitivity analyses used partial rank correlations adjusted for human-assigned LFQ share, log_2_ protein-group depth, antibiotic status and clinical group, and protein signals normalized to the total annotated signal at the same functional level. A further paired-feature sensitivity also adjusted for EDS. Two-sided partial-correlation tests used residual degrees of freedom. Benjamini–Hochberg correction was applied separately within each layer, normalization and model to all eligible features or linked pairs, not only heatmap features. Literature-supported and stereochemistry-dependent follow-up links were marked post hoc and included in the augmented linked-pair families. Full tests, missing targets, identity qualifications and sensitivity results are in Source Data. In a post hoc decomposition of adjustment sensitivity, reported in Source Data only, we examined five metabolite–KO or metabolite–pathway pairs from an earlier follow-up of this screen using all 16 combinations of the four sensitivity covariates. Within each pair, every model used the same positive-measurement, complete-covariate patient subset. A secondary analysis used the 84 patients shared by all five pairs. Both native and annotated-relative protein signals were evaluated. Continuous variables were ranked within each analysis subset. We used 1,000 patient bootstrap resamples, shared across models and normalizations within each pair, to calculate pointwise 95% percentile intervals for correlations and their changes from the unadjusted estimate. All model combinations were retained. After inspection of the complete refitted screen, three annotated metabolite–KO pairs were selected post hoc for Supplementary Fig. 10: glucuronic acid–glucuronate isomerase (HILIC_FT23003, K01812, KEGG reaction R01482), tryptophan–tryptophanase (HILIC_FT24070, K01667, R00673), and glycochenodeoxycholic acid– bile salt hydrolase (C18_F2423, K01442, R03975). The candidate set comprised 21 native-signal pairs with inverse metabolite–KO associations, positive metabolite–EDS associations and negative KO–EDS associations, each passing *q*<0.05 in its complete unadjusted screen family. The three illustrated pairs were chosen for interpretable annotated reaction links and distinct biological topics. No significance threshold or testing family was redefined around the displayed pairs. None retained *q*<0.05 after joint adjustment for all four covariates. For these three pairs, a further descriptive decomposition retained all 16 combinations of those covariates under native and annotated-relative normalization, on pair-specific complete subsets (85, 84 and 85 patients) and the common 84-patient subset (192 estimates). This additional decomposition reports point estimates without new *P* values or bootstrap intervals. Full-family *q* values are retained from the original screen (Source Data).

The focused heatmap (Supplementary Fig. 10) displays six rows: three measured metabolites and their annotated enzyme-associated KO signals, arranged as pairs. KO rows used peptide-based functional allocation excluding human-assigned signal, with unresolved origin retained. Metabolite measurements came from the companion dataset. In Fig. 4h, human-protein rows used human-assigned peptide signal within retained reporting-group coordinates, and pathway rows used the revised peptide-based functional allocation. The original supplementary sample view and source measurements retain all 20 features across 259 matched observations, including repeated visits, with positive finite values standardized within feature. Neither display used imputation, and the selected-feature list and all previous biochemical follow-up results remain in Source Data.

Exploratory protein-based EDS prediction used the fixed 85-patient roster, one observation per patient and the revised peptide-based OG profiles, with quantified-group counts under direct peptide assignment as the depth covariate. OG features with positive intensity in at least 10% of training observations were transformed to square-root relative abundance. Models used protein features alone, log_10_ protein-group depth plus human-assigned intensity fraction alone, or both. Baseline variables were standardized using training-fold means and population standard deviations. Ridge penalties 0.01, 0.1, 1, 10 and 100 were selected by mean squared error in three inner folds within each of five shuffled patient-separated outer folds. Three outer split seeds (20260915–20260917) were retained. A separate distance score averaged the five nearest training profiles among those in the training EDS lower quartile. All filtering, transformation estimates, penalty choices and low-EDS reference selection were confined to training data. For the first outer split, 199 label permutations repeated model selection or low-EDS reference selection.

Two-sided Spearman permutation probabilities used the plus-one correction and Benjamini–Hochberg adjustment across the four model tests. Other split seeds assessed sensitivity. Two thousand patient resamples (seed 20260918) of fixed held-out predictions provided 95% intervals for correlation and baseline-minus-model mean absolute error. These intervals condition on the fitted folds. All held-out predictions, folds, model metrics, selected penalties, permutation outcomes and depth/host controls are supplied.

#### Native cross-omics profiles and sequence coverage

The native pyramids within Figs. 4i and 5g used native gene-sequence transcripts per million (TPM), protein-group intensity and metabolite features without OG, KO or pathway summation. PREDICT retained one earliest complete-stack stool specimen per patient (77 patients), with repeated protein acquisitions averaged within the specimen roster. CapScan retained 13 participants with complete DNA profiles in all five sampling groups, using 245 DNA runs. Protein profiles were normalized within acquisition, collapsed by median within specimen and averaged within participant–region. DNA runs were averaged within specimen, normalized and averaged within participant–region. Metabolite profiles (log10) were averaged within participant–region. Exact per-layer rosters are in Source Data.

DNA/RNA coordinates were identical full encoded amino-acid sequences after removal of terminal stop symbols. Only exact sequences were combined. Reused sample-local gene identifiers were not matched across unrelated assemblies. This representation combines synonymous nucleotide variants but applies no similarity clustering or functional grouping. No gene was selected by de novo support, protein detection or annotation. Protein measurements used the reporting groups under direct peptide assignment. Within each group, human-assigned peptide features (Supplementary Methods 5) formed the human-assigned layer. The remaining peptide signal formed the other/unresolved layer. The two layers conserved each group’s measured intensity. A group could contribute different peptide features to both layers.

Upstream sequence-resolution coverage was quantified as the retained TPM fraction for every DNA/RNA source. Unresolved sequence identifiers were excluded from native coordinates but their TPM mass was retained in coverage tables. Feature-prevalence filtering is a separate operation on the sequence-resolved profiles. All native, functional and cross-omics profile comparisons retain every feature observed in the analysis population. A companion analysis restricted gene and protein features to those present in at least 10% of profiles. Because this retained a median of 26.5% and 21.0% of sequence-resolved PREDICT DNA and RNA abundance, the filtered analysis is reported only in Supplementary Fig. 11c,d and Source Data.

Gene and protein layers were normalized to within-profile relative abundance and square-root transformed. Euclidean distances then represent Hellinger geometry [14]. Metabolites retained features finite in at least 70% of profiles, used log_10_ intensity, feature-median imputation and standardization after removing near-constant features. In CapScan, each transformed feature was residualized against average region effects within the 13-participant complete population. Spearman correlations compared the upper triangles of layer-specific distance matrices. Two-sided tests used 1,999 patient-label permutations in PREDICT or whole-participant-block permutations in CapScan, with seed 20260907 and plus-one corrections. Benjamini– Hochberg correction used complete families of 10 PREDICT or 6 CapScan native-profile comparisons. Functional-profile families are specified below. These profile comparisons do not require matching feature identities between molecular layers.

#### Controlled distance sensitivity

The controlled comparison crossed Hellinger/Euclidean and rank-profile distances with outer Pearson or Spearman correlations while holding native input features and metabolite preprocessing fixed. Both all-observed and ≥10% prevalence inputs were retained. The unadjusted rank-profile dissimilarity was one minus within-layer Spearman correlation. For regional adjustment, each profile’s feature ranks were centered and normalized to unit length. Half their squared Euclidean distance equals one minus Spearman. Region means were subtracted in this rank-feature space, followed by half squared residual Euclidean distance. This adjusted quantity was labeled region-adjusted rank-profile distance.

PREDICT was unadjusted. CapScan retained both adjusted and unadjusted variants. Every method/adjustment/input variant used the same design-preserving 1,999 permutations and its own 10- or 6-comparison Benjamini–Hochberg family. All 176 estimates from these controlled distance comparisons were retained (Supplementary Fig. 11e,f).

Functional and native distances measure concordance between overall profiles. Individual-function correlations compare matching functions across layers. Under controlled rank distances, 9 of 24 region-adjusted CapScan comparisons in the 13-participant population did not meet the significance threshold: one of four non-host protein–metabolite, all four host protein–metabolite, two of four DNA–metabolite and two of four DNA–host protein comparisons. All 24 Hellinger-distance comparisons met it.

#### PREDICT functional-profile comparisons (Fig. 4i and Supplementary Fig. 11a)

Cross-omics analyses compared between-profile geometry rather than treating individual pairwise patient distances as independent observations. DNA, RNA and protein profiles were represented at orthologous-group, KEGG ortholog and pathway resolution. Protein functional profiles excluded human-assigned peptide signal and retained unresolved origin. The separate human layer used human-assigned signal within fixed reporting groups and metabolites retained their native feature space. Profile transformations followed the native comparisons. Concordance between layers was the Spearman correlation between upper triangles of the Euclidean distance matrices. Two-sided significance was assessed using 1,999 patient-label permutations, permuting rows and columns of one distance matrix together and using a plus-one correction. Benjamini–Hochberg correction was applied across the 28 PREDICT profile-level comparisons, which include pathways although the main panels display OG and KO. The host-protein–metabolite comparison is counted once, under OG. A sensitivity restricting DNA, RNA and protein features to those positive in at least 10% of profiles is in Source Data. For PREDICT functional profiles, DNA and RNA abundances were gene-level TPM values summed separately within each sample by bacterial/archaeal eggNOG orthologous group. For KEGG ortholog and pathway profiles, each eligible gene’s TPM was divided equally among its distinct annotations at that level before summation. Protein functional profiles instead summed quantified peptide-feature intensities allocated equally as described in Supplementary Methods 5. Signals were aggregated within each sample and molecular layer before normalization and between-patient distance calculation. Across the same 77 native profiles, 5,391,526 encoded-sequence features had positive RNA abundance, compared with 14,822,078 for DNA.

Individual-function correspondence was analyzed separately. For each functional representation and pair of microbial DNA, RNA and protein layers, profiles were normalized over function identifiers shared by that layer pair. Spearman correlations were then calculated across patients for each individual function, requiring at least 35 patients with positive measurements in both layers and a finite correlation statistic. A function tested in more than one layer pair contributed separately to each analysis. Benjamini–Hochberg correction was applied separately within each of the nine representation-by-layer-pair families. Under the revised allocation, 30,581 function-by-layer-pair comparisons met these criteria. Here, the revised allocation is the functional allocation described in Supplementary Methods 5 and used for all other functional results. The previous allocation is the earlier allocation of the same PREDICT profiles, made before those annotation sources and that reference scope were adopted, under which the common functions had median DNA–protein and RNA– protein correlations of 0.10–0.12. To assess dependence on feature eligibility, we also summarized the revised correlations for functions eligible under both the previous and revised allocations, matching by functional representation, molecular-layer pair and function identifier. We retained the original eligibility rules and did not renormalize profiles, refit correlations or redefine multiple-testing families for this subset. Common, newly eligible and lost-feature counts and the corresponding median correlations are provided in Supplementary Table 4 and Source Data.

#### CapScan functional-profile comparisons (Fig. 5g and Supplementary Fig. 11)

CapScan functional cross-omics concordance used complete participant-by-region profiles spanning regions 1–4 and stool. All 16 comparisons among microbial DNA, microbial proteins, host proteins and metabolites used the same 13 participants with complete DNA profiles (65 profiles), matched at participant–region. For DNA, gene TPM values from 301 completed metagenome runs were allocated to bacterial or archaeal OGs, KOs and pathways using eggNOG annotations, splitting each gene’s TPM equally among its distinct annotations within each representation. Only genes with a single bacterial or archaeal root OG contributed, and map011 global maps were excluded. Run profiles were averaged within specimen, renormalized over annotated functions and averaged within participant and region. Protein and DNA features were transformed to square-root relative abundances. Metabolites were preprocessed as for the native profiles.

Region effects were removed separately for each transformed feature using ordinary least squares with an intercept and four region indicators within the 13-participant population. In this balanced design, this corresponds to subtracting each sampling group’s mean transformed profile (regions 1–4 and stool) from every profile in that group. Concordance was the Spearman correlation between upper triangles of the Euclidean distance matrices of region-adjusted profiles. Two-sided *P* values were calculated using 1,999 whole-participant block permutations, preserving the five-region structure within each participant. Benjamini– Hochberg correction was applied jointly across all 16 functional-profile comparisons: OG/KO in Fig. 5g and pathways in Supplementary Fig. 11b, with all 16 in Source Data.

Two sensitivities are retained in Source Data: the same comparisons with a ≥10% feature-prevalence filter, and a CapScan family in which seven non-DNA comparisons used 15 participants (75 profiles) and nine DNA comparisons used 13. Across these filters and populations, no comparison changed classification at adjusted *q*<0.05, although coefficients changed.

The Source Data contain the 120 functional and 32 native correlation results, the functional and native distance matrices, participant and specimen rosters, eligibility tables, all plotted cross-omics values and the 176 controlled estimates. All prevalence, population and distance-method results are retained.

#### CapScan regional analyses

Fifteen participants contributed paired intestinal-fluid and stool measurements across all five sampling groups. For human-assigned-share comparisons, mean sampling-region versus stool differences were estimated across participants, with 95% intervals from 2,000 participant-bootstrap resamples (seed 20261003). Two-sided paired sign-flip tests enumerated all 32,768 possible sign patterns across the 15 participants, with Benjamini–Hochberg correction across the four regional contrasts.

#### Participant-specific microbial functional profiles

For the own-stool comparison in Fig. 5c, we built peptide-based microbial orthologous-group profiles (microbial single-OG profiles, Supplementary Methods 5). We withheld human-assigned signal and retained signal of unresolved origin. Each acquisition’s annotated OG profile was normalized to its sum and then averaged within specimen and participant–region, yielding 75 profiles from the 15 participants sampled in all five regions. All 73,316 orthologous groups observed in these profiles were retained. Restricting to OGs present in at least 10% (53,023 OGs) or 20% (44,330 OGs) of profiles gave the same result, with every participant closer to their own stool and *P=1 × 10*⁻⁵. Profiles were square-root transformed and compared using Euclidean distance. The comparator used distances to each other participant’s stool, not a pooled stool centroid.

Significance was assessed by the permutation test described in Methods, using a plus-one *P* value. Participant and region contributions to functional-profile variation were summarized descriptively using an additive participant-plus-region model in the same transformed feature space. Sums-of-squares shares for participant, region and residual variation were reported as variance partitions. To distinguish the intestinal fluid–stool contrast from variation among intestinal sampling regions, we further decomposed the sampling-group sum of squares in the balanced 15-participant, five-group analysis. With transformed group-mean profiles ordered as regions 1–4 and stool, the unit-length contrast (1,1,1,1,−4)/√20 defines the capsule–stool component. Its squared vector norm was multiplied by 15. The among-capsule component was 15 times the sum of squared deviations of the four capsule-region means from their capsule mean. These orthogonal components sum to the original sampling-group sum of squares. Both were divided by the same total sum of squares of the 75 transformed participant–group profiles about their grand mean, yielding 10.7% and 4.2%, respectively. The separate intestinal-fluid-only partition was calculated from 60 profiles and uses its own total sum of squares. Separately, we quantified protein-signal evenness using Pielou’s index, J = −Σᵢ pᵢ ln(pᵢ)/ln(S), where pᵢ is the relative KO-annotated protein signal assigned to named species i and S is the number of species with positive signal. This analysis used the 346 non-empty CapScan acquisitions and excluded host, mixed host/microbial and unresolved species assignments before normalizing the named-species contributions to unit sum. We calculated evenness for each acquisition and compared, for each participant, the median over all intestinal-fluid acquisitions (regions pooled) with the median over stool acquisitions, using a two-sided exact Wilcoxon signed-rank test. This measures the distribution of assigned protein signal. (analysis: fastalake-paper, pipelines/predict_clinical_refit, predict_rifaximin_sensitivity, predict_eds_screen, predict_eds_feature_selection, predict_eds_focused_pairs, predict_eds_pair_adjustments, predict_eds_heldout_prediction, predict_crossomics_refit, native_crossomics_profiles, crossomics_distance_sensitivity, predict_function_correlations, peptide_centric_downstream, capscan_crossomics, capscan_metagenome_profiles, capscan_functional_profiles and capscan_dominance_evenness)

### Supplementary Methods 9 | Non-specific searches and histidine boundaries

#### Non-specific searches and peptide termini

Non-specific searches used Sage v0.14.7 with no protease cleavage rule. Search scope, peptide gates and the non-K/R definition are given in Methods. To measure the additional candidate burden, we compared archived non-specific and tryptic searches of the same assembly-inclusive acquisition FASTAs in all 347 CapScan acquisitions. Sage scored_candidates counts were paired on native scans reported at rank one by both searches, without a q-value filter, and include target/decoy candidates summed over precursor-charge and isotope hypotheses. We calculated the median within each acquisition and then the median across acquisitions; fold changes were paired ratios of acquisition medians. The medians were 20,247.5 and 218 candidates per shared spectrum, respectively, with a median paired ratio of 89.84. These are scored-candidate counts, not counts of unique peptide sequences. Semi-tryptic and non-specific searches used their corresponding Sage cleavage rules.

#### Peptide-origin classification

Peptide-origin labels were assigned from all compatible parent proteins in the acquisition’s selected FASTA under I/L-equivalent sequence containment and the cleavage rules of the corresponding analytical route. Proteins with recognizable human accessions were classified as human and other readable microbial accessions as microbial. Under the conservative primary rule, peptides compatible with both classes were classified as mixed, and peptides involving unreadable hashed accessions were classified as unknown. Fig. 6b includes only peptides classified as human or microbial. Mixed and unknown assignments were retained separately in Source Data.

Acquisition-level terminal fractions were averaged within specimen and then within participant and region. The region 1–2 intestinal-fluid summary comprised the mean of regions 1 and 2, and the region 3–4 summary comprised the mean of regions 3 and 4 in participants with the corresponding measurements. For each participant, the intestinal fluid–stool difference was the mean of the four sampling-region values, each region weighted equally, minus that participant’s stool value, and paired tests assessed whether the mean of these differences across the 15 participants differed from zero.

#### Internal-boundary enrichment

Boundary analyses retained peptides with a uniquely localized internal boundary in the selected source proteins. Unique localization refers to one compatible protein-sequence/start/end location under the route-specific mapping rules and does not require that only one spectrum or peptide observation support that boundary. For each observed internal non-K/R boundary, the source expectation was conditioned on the compatible source protein and analytical route. Histidine enrichment and its source background were defined as in Methods. Three analytical routes were evaluated for CapScan: top-30% de novo predictions, semi-tryptic Sage searches and non-specific Sage searches. The database-search routes required the specified Sage confidence gates. De novo predictions did not.

PREDICT boundary analyses used de novo predictions and non-specific Sage searches. No cohort-wide semi-tryptic PREDICT boundary series was used. Observed-minus-expected values were averaged within participant and region, or within PREDICT patient across available specimens. Paired or sign-flip inference was performed within the analysis-specific multiple-testing families defined in Source Data. Peptide-only routes are retained separately as sensitivities. Internal-boundary and complementary-product intervals use 20,000 whole-participant or whole-patient bootstrap resamples. Source-enrichment tests require at least eight units and enumerate all signs when *n*≤16. Larger groups use 200,000 sign assignments and a plus-one correction. Fig. 6c residue-composition intervals use 2,000 participant-bootstrap resamples. The broader non-K/R fraction intervals use 50,000.

#### Complementary peptide products

A complementary product was defined as a distinct observed peptide, supported by a distinct spectrum, whose boundary mapped to the opposite side of exactly the same localized protein boundary within the same acquisition and analytical route. Histidine boundaries were compared with eligible non-histidine boundaries matched within the same source protein for theoretical peptide availability on each side and observed spectrum-depth stratum. The complementary-product analysis was performed separately for de novo, semi-tryptic and non-specific routes. Because these routes share spectra, participants and FastaLake-selected sequence spaces, they were treated as complementary analytical evidence rather than independent replication. Concordant enrichment across the three routes was interpreted specifically for CapScan intestinal fluid. Sequence-context analysis retained the full residue-position matrices. Tests of flanking motifs excluded the fixed histidine-conditioning P1 position. The CapScan motif family included 32,400 comparisons across the retained routes, sides, origins, conditioning groups, flanking positions and residues. Negative complementary-product contrasts in stool de novo predictions, strongest in PREDICT, were retained in the complete source tables.

#### Selection and inspection of measured spectra

Measured-spectrum examples were selected within 12 cohort/compartment/origin/boundary strata: eight CapScan strata (intestinal fluid or stool, human or microbial, N-terminal or C-terminal histidine) and four PREDICT stool strata. Eligible cases were 9–50-residue peptides absent from the same acquisition’s accepted tryptic set, with a uniquely localized internal histidine boundary and rank-one target evidence passing both peptide and spectrum *q*≤0.01. Acquisitions with positive candidate counts were ordered by count and sample identifier. The middle acquisition was selected, with a deterministic nearest-rank fallback if no candidate passed the stricter PSM gates. Within an acquisition, canonical peptides were represented by lowest peptide *q*, then highest hyperscore and lexical scan/peptide order. Cases were ordered by matched-intensity percentage, canonical peptide and scan identifier, and the middle case was chosen. Selection denominators and empty strata were retained.

Actual mzML peaks were extracted and b/y annotations independently checked against Sage matched-ion counts. Local-FASTA substitutions and adjacent-residue swaps were evaluated within the permitted cysteine/methionine modification and isotope-error settings. All eligible alternatives, including equal or better fragment matches, were retained. This was a bounded fragment-count diagnostic. Same-scan calls from the existing routes were retained regardless of acceptance. A preceding histidine for an N-terminal boundary is inferred from protein context, not observed within the peptide. (analysis: fastalake-paper, pipelines/capscan_nonspecific_search, predict_nonspecific_search, terminal_route_counts, terminal_participant_inference, denovo_threshold_routes, terminal_histidine_inference, histidine_boundary_localisation, histidine_spectrum_evidence and predict_histidine_case_audit)

### Supplementary Methods 10 | Statistical analysis

Analysis units, test sidedness, resampling and multiple-testing conventions are given in Methods. Function cells were also not treated as independent biological replicates. Positive-tail *P* values retained for the distance-method sensitivity are labeled separately.

#### Testing-family map

Benjamini–Hochberg correction families were (i) two PREDICT antibiotic endpoints, separately for the primary and each rifaximin sensitivity definition, (ii) four CapScan regional human-assigned-share contrasts, (iii) 28 PREDICT and 16 CapScan functional-profile contrasts, (iv) 10 PREDICT and 6 CapScan native-profile contrasts separately for each feature/distance/outer-correlation/adjustment variant, (v) nine representation-by-layer-pair families for individual-function correlations, (vi) EDS association screens within each layer, normalization and model, and (vii) the four held-out EDS model tests. The seven CapScan non-DNA comparisons repeated in all 15 participants are a separate sensitivity. Histidine analyses retained complete families: (i) CapScan terminal fraction 4,536, (ii) residue composition 13,608, (iii) internal source residue 9,000 CapScan and 1,560 PREDICT, (iv) complementary products 90 and 12, and (v) CapScan spatial histidine 2,520 and motifs 32,400. Unavailable planned hypotheses and non-significant tests remain in the supplied families. No correction family was limited to displayed or significant comparisons.

The panel/source map identifies the numerical table, row selection, columns, analysis unit, producing code and manuscript explanation for each panel. Source tables retain negative results, unresolved categories and non-displayed sensitivity strata. (analysis: fastalake-paper, text_statistics and figures/<figure>/panels.py; docs/STATISTICS.md)

### Supplementary Methods 11 | Computational resources and provenance

#### Sequencing cost and computational performance

Sequencing cost and turnaround comparisons use dated provider examples, retained in the computational source table [15–18]. They are not measured study expenditure or a standardized complete workflow. Costs depend on read depth, library preparation, batch and tax, and may omit extraction, quality control, assembly and gene prediction.

Acquisition-level CPU stages were restricted by array-index/sample mapping to the 15 CAMPI, 58 healthy-stool, 306 PREDICT and 347 CapScan manuscript acquisitions. Shared healthy-stool and CapScan jobs retain their actual 97- and 368-acquisition execution scopes. The CAMPI sequence-selection and search chain used the broad UniProt reservoir. Wall minutes are job execution elapsed time. Allocated CPU-hours are distinct from consumed CPU time and end-to-end workflow time. Peak memory is expressed in GiB (2³⁰ bytes). Memory summaries report the number of usable samples. Acquisition medians are not added to estimate workflow turnaround. Queueing, raw conversion, de novo inference and downstream biological analyses are outside these CPU-stage summaries. Available CAMPI prediction logs retain a separate execution span and hardware record. The recorded CAMPI reservoir build took 92 min with 64 threads on a 512-GB AMD EPYC node, and pooled extraction took 11 min. Median acquisition-level extraction, selection and Sage/LFQ times were 0.48, 0.63 and 0.23 min; AlphaNovo inference had a recorded median of 28 min on an NVIDIA L40S. Chunked public-reservoir workflow measurements used an 11-GiB allocation: two healthy-stool acquisitions took 38 min 52 s with an 8.4-GiB largest-process peak, and 24 took 3 h 35 min with a 10.2-GiB peak. Separately, the archived non-specific CapScan searches had a median whole-process wall time of 25.6 min and median maximum resident set size of 208.6 GiB across 347 acquisitions. These non-specific searches did not use the 11-GiB chunked workflow.

#### Software and data provenance

The principal software comprised AlphaNovo [6], Sage (v0.14.7 release binary, reporting version 0.14.6, and the corrected build in Supplementary Methods 7) [7], eggNOG-mapper 2.1.12 [8] in DIAMOND mode [9] with eggNOG database 5.0.2 and DIAMOND 2.0.11, ProteoWizard 3.0.24025 for the original mzML conversions, ThermoRawFileParser 2.0.0.0 and 1.3.4 for replacement mzML files, MSFragger 4.1 [19], Percolator 3.06.5 [20], MMseqs2/Linclust (commit 76da68ad7577378410c075049e18666fcc94f8d1) [21,22], FDRBench v0.0.4 [11], AlphaPeptDeep [23] and directLFQ 0.3.3 [24]. directLFQ was used for the common-study quantification sensitivity in Supplementary Fig. 7. FastaLake’s Rust binaries were compiled with the pinned Rust toolchain 1.91.1. Origin annotation used GTDB release 202 [25] and the NCBI taxonomy snapshot of 30 September 2026. The principal downstream statistical environment used Python 3.12.4, NumPy 2.2.6, SciPy 1.16.3, pandas 2.3.3 and statsmodels 0.14.5; the separate pooled cross-fitting rescoring environment is specified in Supplementary Methods 7.

Study-level protein groups were produced by study.py run through run_cohort.py. The released module (MannLabs/fasta-lake, fasta_lake/study.py at commit 9ad09c9) reproduced the production healthy-stool grouping exactly, with identical output tables for the 58 acquisitions, including 59,428 quantified protein groups and 1,909,973 quantified peptide features. (software: MannLabs/fasta-lake, fasta_lake/study.py; analysis: fastalake-paper, pipelines/campi_simplified_grouping/run_cohort.py and common_resource_accounting)

